# MtvS1 and MtvS2 Interact with RNA Polymerase to Regulate the *Francisella* Type V-A CRISPR-Cas System

**DOI:** 10.1101/2024.09.12.612765

**Authors:** Maj Brodmann, Christian F. Baca, Joshua Chandanani, Elizabeth A. Campbell, Luciano A. Marraffini

## Abstract

Bacteria and archaea often harbor multiple CRISPR-Cas loci to defend against mobile genetic elements. Little is known, however, about whether and how different CRISPR-Cas systems are differentially regulated, in many instances due to the impossibility of studying CRISPR immunity in native hosts. Here we investigated the regulation of the endogenous type II-B and type V-A CRISPR-Cas systems present in the opportunistic human pathogen *Francisella novicida* U112. We found that while the type II-B system is constitutively expressed, the type V-A system is differentially expressed at stationary phase and high cell density. We identified MtvS1 and MtvS2 as factors required for this regulation, as well as for the modulation of many additional genes in stationary phase, some of which are required for *Francisella* virulence. Both *Francisella* MtvS proteins bind to RNA polymerase. MtvS1 is predicted to interact with the β’ subunit of the RNA polymerase, and MtvS2 with multiple RNA polymerase subunits as well as MtvS1. We propose that MtvS1 and MtvS2 constitute noncanonical alternative sigma factors involved in the regulation of the expression of the type V-A CRISPR locus and other genes in *Francisella*. Last, we show that the MtvS1 homolog YgfB is required for expression of the type I-E CRISPR-Cas system in *E. coli*, a result that suggests a broader role in gene regulation for these alternative sigma factors.

## Introduction

CRISPR (Clustered regularly short interspaced palindromic repeats)-Cas (CRISPR associated genes) systems protect bacteria and archaea against foreign nucleic acids such as phages and genetic mobile elements^1,2^. CRISPR loci are composed of an array of short (30-40 bp) repetitive sequences separated by equally short invader-derived sequences known as spacers. CRISPR-Cas systems provide adaptive immunity through the acquisition of spacer sequences during infection that are stored as immunological memories of previous encounters with mobile genetic elements^3–5^. CRISPR arrays are transcribed and processed into small CRISPR RNAs (crRNAs) that use the spacer sequences to guide Cas effectors to complementary sequences present on invading foreign genomes^6,7^. Commonly, Cas effectors are crRNA-guided nucleases that cleave the infecting DNA to stop infection ^8–13^. Prokaryotic organisms often encode multiple CRISPR-Cas loci, which may be differently regulated^14^.

Regulation of the CRISPR-Cas defense is important to balance an effective immune response that minimizes autoimmunity and limits, but does not completely abrogates, potentially beneficial mobile genetic elements^1,15–18^. Previous studies have shown that regulation of the CRISPR-Cas systems can be affected by secondary metabolites and nanoRNAs^19–24^, H-NS-mediated chromosome condensation and membrane stress^19,20,25–27^, environmental cues sensed by two component signaling systems^28,29^ and cell density or virulence^30–33^. Many CRISPR-Cas systems can be autoregulated by repression of their own transcription through the Cas effectors^34–38^. Other CRISPR-Cas systems encode their own specific transcriptional regulator or putative DNA-binding proteins within or nearby the CRISPR-*cas* locus^34,39–43^. Detailed investigation of regulatory networks governing CRISPR expression, however, has been difficult due to the lack of natural hosts that can be used as experimental systems to study the CRISPR-Cas response.

Here we studied the endogenous type II-B and type V-A CRISPR-Cas systems in the opportunistic human pathogen *Francisella tularensis* subspecies *novicida* Utah 112 (here *F. novicida* U112) (Supplementary Fig. 1a-b)^13,44,45^. It has been shown previously that both *F. novicida* U112 CRISPR loci are functional and capable of restricting foreign DNA^46,47^. Interestingly, *F. novicida* U112 Cas9 is also required for virulence. It can use a special guide RNA known as the scaRNA to repress expression of a lipoprotein with unknown function that enhances membrane stability during *Francisella* colonization ^48–51^. However, little is known about how these two systems are transcriptionally controlled, especially in the light of the limited understanding of transcription regulation in *Francisella*^52^. For example, *Francisella* genomes encode only two canonical sigma factors (σ^70^ and σ^32^) that dictate promoter specificity and initiation of transcription^52,53^. *Francisella* also has a unique RNA polymerase composed of two heterologous α subunits (instead of two identical subunits) which may also impact promoter recognition^54–56^.

Using fluorescence microscopy, we found that the two *F. novicida* U112 CRISPR-Cas systems are differentially regulated. While the type II-B CRISPR-Cas system is constitutively expressed, the type V-A CRISPR-Cas system displays significantly higher expression during stationary phase. Using mass spectrometry and genetics, we identified MtvS1 and MtvS2 (Modulators of the type V-A CRISPR-Cas system expression in stationary phase, FTN_1238 and FTN_1519), proteins with previously unknown function in *Francisella* species, as factors required for the differential expression of the type V-A CRISPR-Cas locus. Additionally, we showed that type V-A CRISPR-Cas system expression is induced upon introduction to spent overnight medium suggesting that *F. novicida* U112 is capable of quorum sensing despite lacking canonical quorum sensing genes. We demonstrated that both MtvS proteins can bind and pull-down RNA polymerase. AlphaFold3 structural predictions suggest interactions between MtvS1 and the β’ subunit and between MtvS2 and both β and β’ subunits, as well as MtvS1. Since both MtvS proteins affect not only the expression of the type V-A CRISPR-Cas locus but also many additional genes in stationary phase, we hypothesize that MtvS1-2 may act as noncanonical sigma factors. While MtvS2 is only found within *Francisella* species, our bioinformatics analysis revealed that MtvS1 homologs are enriched with type I and type IV CRISPR-Cas systems, and we demonstrated that the *E. coli* K-12 *substr*. BW25113 (here *E. coli*) homolog YgfB is important for upregulation of the *E. coli* type I-E CRISPR-Cas system in stationary phase. Last, we showed that MtvS1 and YgfB are functionally redundant, suggesting a broader role for this protein family in modulating bacterial transcription in stationary phase.

## Results

### The F. novicida U112 type V-A CRISPR-Cas locus is upregulated during stationary phase

To investigate the expression pattern of the type II-B and type V-A CRISPR-Cas systems in *F. novicida* U112, their RNA-guided nucleases were C-terminally tagged to generate Cas9-msfGFP and Cas12-mCherry2, respectively. The fluorescent tags did not affect their ability to prevent plasmid conjugation (Supplementary Fig. 1c). Fluorescence live-cell imaging of exponentially growing bacteria and of cells in stationary phase revealed that fluorescence intensity was comparable in both growth phases for Cas9-msfGFP while fluorescent intensity for Cas12-mCherry2 was higher in stationary phase (Fig. 1a). Relative quantification of the untagged expressed proteins in the parental strain revealed broad changes in protein expression between exponential and stationary phase cells (Fig. 1b, Supplementary Data 1, tabs A-B). Gene set enrichment (GSE) analysis (Supplementary Data 1, tab B) suggested that proteins involved in DNA replication and cell envelope biogenesis were expressed at significantly higher levels in exponential growing cells. On the other hand, DNA repair and anti-stress proteins were more prominent in stationary phase cells. More important, this untargeted proteomics approach corroborated that Cas12 expression was increased ∼1.8-fold in stationary phase and showed no phase variation for Cas9 (Fig. 1b, Supplementary Data 1, tab A). To investigate whether these distinct expression patterns of Cas9 and Cas12 originate from different transcription levels, we performed RNA-seq. Similar to the proteomics data, comparison of RNA levels of genes between exponentially growing and stationary phase bacteria revealed broad changes in gene expression between the two growth conditions. Genes required for protein synthesis were expressed higher in cells growing exponentially and metabolic genes were upregulated in stationary phase (Fig. 1c, Supplementary Data 1, tabs C-E). Importantly, *cas12* and *cas2_type V-A_* RNA levels were significantly higher during stationary phase than in exponential growth phase. In comparison, components from the type II-B CRISPR-Cas system, except *cas2*, had comparable RNA levels in the two tested conditions. Targeted measurement of RNA levels of *cas9* and *cas12* by RT-qPCR confirmed the invariability of RNA levels of *cas9* in the two growth conditions and the distinct temporal expression of *cas12* in stationary growth phase (Figs. 1d-e). Together these results show that the two effector nucleases from the *Francisella* type II-B and type V-A CRISPR-Cas systems have different expression patterns and that *cas12* expression levels significantly increase in stationary phase.

**Figure 1.**
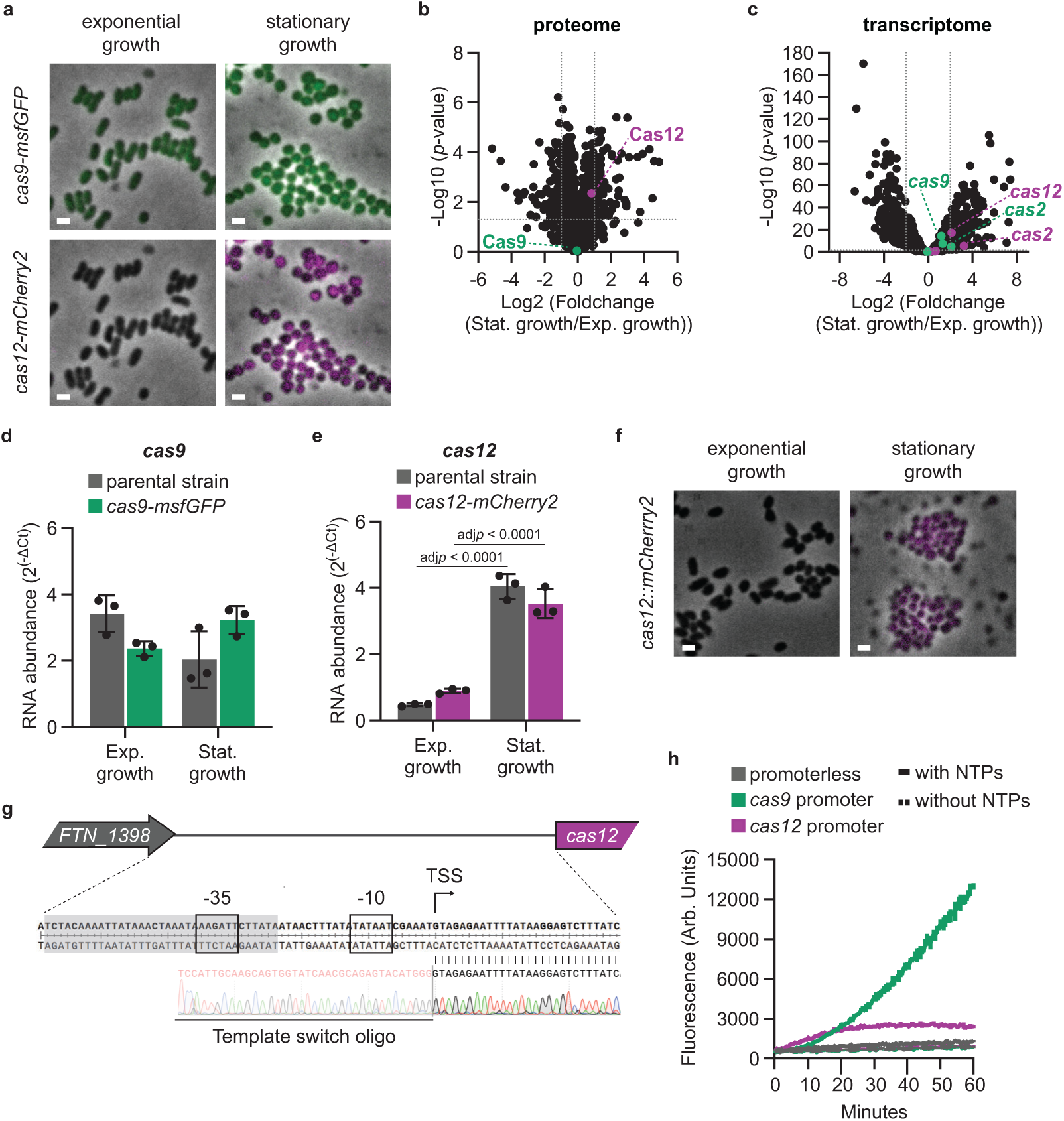
CRISPR nucleases Cas9 and Cas 12 are differentially expressed during exponential and stationary growth. **a** Fluorescence microscopy of Cas9-msfGFP and Cas12-mCherry2. Fields of view of 13 x 13 μm are shown. Upper panel is a merge of phase contrast and GFP channel, lower panel is a merge of phase contrast and RFP channel. Scale bars represent 1 μm. Representative images are shown (n = 3 biological replicates). Source data are provided as a Source Data file. **b** Proteomics data of differentially expressed proteins in stationary and exponential growth phase for parental strain. Green: type II-B CRISPR-Cas proteins. Magenta: type V-A CRISPR-Cas proteins. Dotted horizontal line: *p*-value of 0.05 (two-sample *t*-test). Dotted vertical lines: 2-fold difference. n = 3 biological replicates. **c** RNA-seq data of differentially expressed genes in stationary and exponential growth phase for parental strain. Green: *type II-B CRISPR-Cas* genes. Magenta: *type V-A CRISPR-Cas* genes. Dotted horizontal line: *p*-value of 0.05 (Wald test). Dotted vertical line: 4-fold difference. n = 3 biological replicates. **d-e** Determination of mRNA levels for *cas9* **(d)** and *cas12* **(e)** by RT-qPCR. Grey bars: parental strain. Green bars: *cas9-msfGFP*. Magenta bars: *cas12-mCherry2*. Means with standard deviations are displayed (n = 3 biological replicates). Significant differences were determined with a two-way ANOVA and Tukey’s multiple comparison test (95 % confidence interval). Source data are provided as a Source Data file. **f** Fluorescence microscopy of mCherry2 in a *cas12::mCherry2* strain. A merge of phase contrast and RFP channel in 13 x 13 μm fields of view are shown. Scale bars represent 1 μm. Representative images are shown (n = 3 biological replicates). Source data are provided as a Source Data file. **g** Promoter region of *cas12*. Transcription start site (TSS) determined by 5’ RACE. Black boxes: predicted canonical -10 element and non-canonical -35 element. Light grey box: putative single repeat unit. n = 3 biological replicates. **h** Fluorescence based *in vitro* transcription assay with *E. coli* RNA polymerase holoenzyme and promoterless (grey), *cas9* promoter (green) and *cas12* promoter (magenta) aptamers. A representative assay (mean and standard deviation, n = 3 technical replicates, repeated twice) with (solid line) and without (dashed line) NTPs added is shown. Source data are provided as a Source Data file.

### MtvS1 and MtvS2 are required for upregulation of the type V-A CRISPR-Cas locus in stationary phase

In other organisms, type V-A CRISPR-Cas systems have been shown to be autorepressed by Cas12 loaded with *cas*-regulating RNAs^36^. Moreover, *F. novicida* U112 *cas12* promoter sequence contains a predicted single repeat unit (SRU, with a similar but not identical sequence to the canonical CRISPR repeat, Supplementary Fig. 1b), which was proposed to have a regulatory function^35^. To test if Cas12 is responsible for the upregulation of the type V-CRISPR-*Cas* locus we replaced *cas12* with *mCherry2*. However, the fluorescence pattern observed in Fig. 1a (lower panel) did not change in absence of *cas12* (Fig. 1f). Next, we determined the exact transcription start site by 5’ RACE to identify the *cas12* promoter and its regulatory sequences. The bacterial transcription specificity and initiation factor σ recognizes two conserved core promoter elements, located at approximately −10 and −35 relative to the transcription start site^57–59^. The transcription start site map via 5’ RACE experiments suggested the presence of a putative promoter sequence with a canonical -10 sequence (TATAAT), but a divergent -35 sequence (AAGATT, canonical sequence: TTGACA) that lies within the predicted SRU (Fig. 1g). We then used this promoter sequence to perform *in vitro* transcription assays with *E. coli* RNA polymerase holoenzyme. Using a fluorescence assay (see Methods), we detected only background levels of aptamer transcription with the *cas12* promoter, indicating that it is not recognized by *E. coli* σ^70^ (Fig. 1h). In contrast, an aptamer with the *cas9* promoter was transcribed. In conclusion, these results suggest that *Francisella cas12* requires additional, noncanonical transcription factors for expression.

To identify such factors, we performed a protein pull-down using as baits the *cas12* promoter DNA sequence and an intragenic *cas12* sequence as a control. Lysates from exponentially growing and stationary bacteria were probed and pulled down proteins were identified by LC-MS/MS. All identified proteins are listed in Supplementary Data 1, tabs I-J. An enrichment score was calculated by comparing protein abundance in pull-downs from the *cas12* promoter sequence to the control sequence. In exponentially growing cells, one of the most significantly enriched proteins was the LysR-family transcriptional regulator FTN_0392 (Fig. 2a, Supplementary Data 1, tab I). However, neither Cas12-mCherry2 expression nor type V-A-mediated plasmid targeting was decreased in a *FTN_0392* deletion mutant (Supplementary Fig. 2a-b). On the other hand, in the stationary growth sample, FTN_1238 and FTN_1519, which we renamed *mtvS1* and *mtvS2* respectively (see below), were significantly enriched (Fig. 2b, Supplementary Data 1, tab J). Importantly, there was no enrichment for both proteins when using a lysate from exponentially growing bacteria. *mtvS1* is the first of a five-gene operon also harboring genes encoding for respiratory chain proteins (*ubiH*, *ubiF*), a putative phage protein (*FTN_1235*) and a tRNA modification enzyme (*queA*) (Supplementary Fig. 2c). *mtvS2* is the second gene of a predicted operon also containing *FTN_1520* (amino acid transporter), *RNase P* RNA and *relA* (Supplementary Fig. 2d). Both proteins have no assigned function in *Francisella* but FTN_1238 is structurally homologous to YgfB (Supplementary Fig. 2e), which is conserved in many γ-proteobacteria, including *E. coli* and *Haemophilus influenzae*^60^.

**Figure 2.**
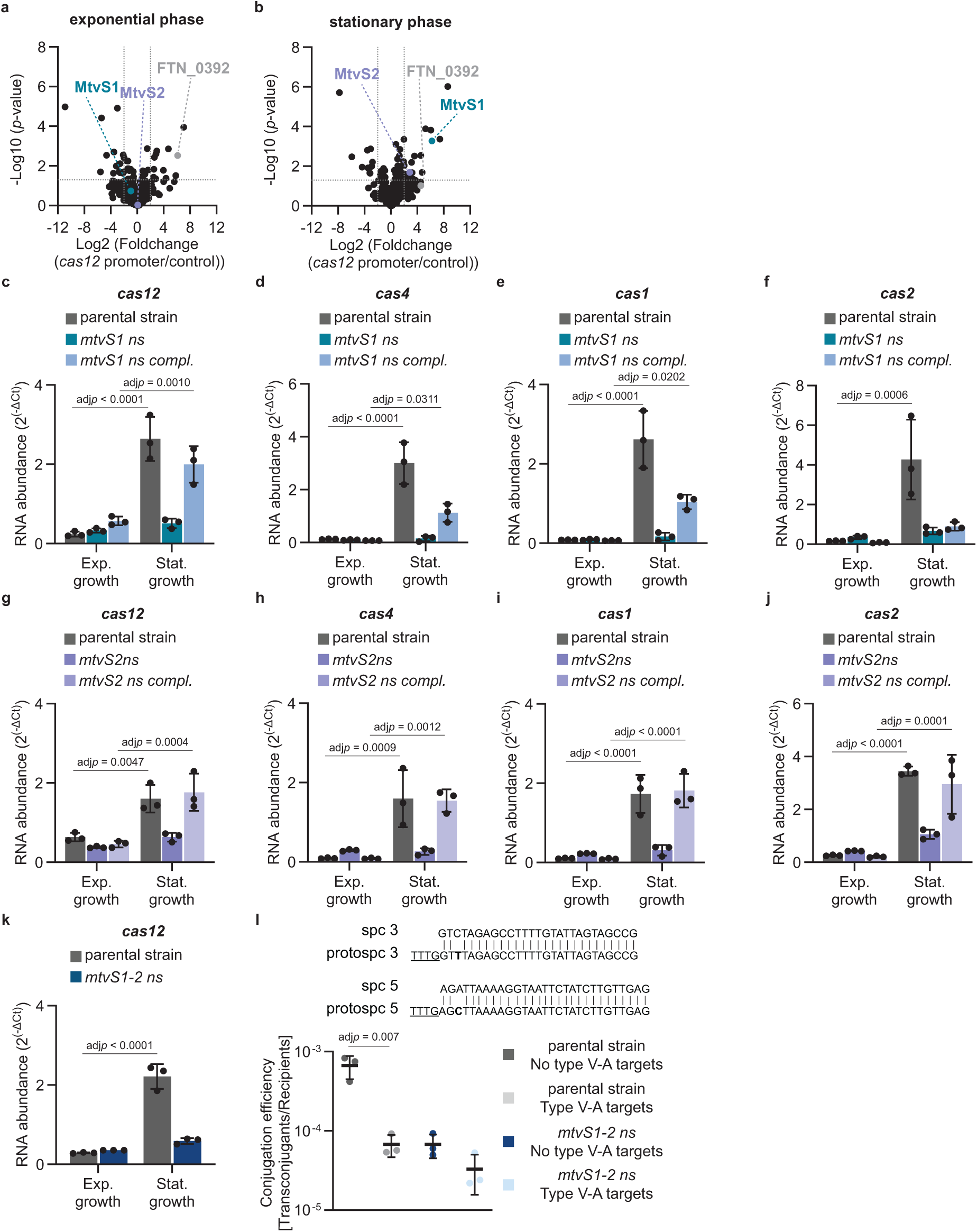
MtvS1 and MtvS2 modulate expression of *cas12* and the *type V -A CRISPR-Cas* system. **a-b** Pull-down of proteins from exponential growth phase **(a)** and stationary phase **(b)** lysate of parental strain binding specifically to the *cas12* promoter DNA sequence. Turquoise circle: MtvS1. Purple circle: MtvS2. Grey circle: FTN_0392. Dotted horizontal line: *p*-value of 0.05 (two-sample *t*-test). Dotted vertical line: 4-fold difference. n = 3 biological replicates. **c-k** Determination of mRNA levels for *cas12* **(c, g, k)**, *cas4* **(c, h)**, *cas1* **(d, i)** and *cas2* **(e, j)** by RT-qPCR. Grey bars: parental strain (with empty plasmid for **g-j**). Turquoise bars: *mtvS1 nonsense* mutant. Light blue bars: *mtvS1 nonsense* mutant with wild-type *mtvS1* at Tn7 insertion site. Purple bars: *mtvS2 nonsense* mutant with empty plasmid. Light purple bars: *mtvS2 nonsense* mutant with wild-type *mtvS2* expressed from plasmid. Blue bars: *mtvS1-2 nonsense* mutant. Means with standard deviations are displayed (n = 3 biological replicates). Significant differences were determined with a two-way ANOVA and Tukey’s multiple comparison test (95 % confidence interval). Source data are provided as a Source Data file. **l** Conjugation efficiency and Cas12-dependent targeting of a plasmid without type V-A targets and a plasmid carrying two protospacers partially homologous to *spacer 3* and *spacer 5* with seed mutations (in bold) in the native CRISPR array. PAM is underlined. Grey circles: parental strain and plasmid without type V-A target. Light grey circles: parental strain and plasmid with type V-A targets. Blue circles: *mtvS1-2 nonsense* mutant and plasmid without type V-A target. Light blue circles: *mtvS1-2 nonsense* mutant and plasmid with type V-A targets. Means with standard deviations are displayed (n = 3 biological replicates). Significant differences were determined a one-way ANOVA and Tukey’s multiple comparison test (95 % confidence interval). Source data are provided as a Source Data file.

To test if these two candidates were involved in regulation of *cas12* expression, we made nonsense mutations by introducing three stop codons after the amino acid at position 10 of both genes individually or combined and analyzed *cas12* RNA levels by RT-qPCR. Importantly, these individual mutations did not affect the growth rate of *F. novicida* U112 (Supplementary Fig. 2f-g). Lack of MtvS1 or MtvS2 significantly affected the RNA levels of all type V-A CRISPR-Cas system genes in stationary phase (Fig. 2c-j). We restored MtvS1 expression in the *mtvS1 nonsense* mutant by integrating a copy of the gene, including its promoter sequence, at the Tn7 integration site downstream of gene *glmS* (*FTN_0485*). With the exception of *cas2*, the transcripts for all the *cas* operon genes were increased (Fig. 2c-f). A possible explanation for lack of restoration of *cas2* transcription is the level of expression of MtvS1 from its alternative chromosomal location in the complemented strain, which could affect the DNA accessibility and transcription dynamics of the integrated copy of *mtvS1*, and thus lower the expression of this gene. If this were the case, given that *cas2* is the last gene of the operon and could display low RNA levels due to transcription polarity, or if the *cas2* transcript is unstable, rescue of the RT-qPCR values could require a higher concentration of MtvS1 than present in the complemented strain. Expression of MtvS2 was restored with a plasmid-borne copy of the *mtvS2* gene that also included its promoter, which was sufficient to increase the RT-qPCR values of all *cas* genes in the operon to parental levels (Fig. 2g-j). The *cas12* phenotype for the *mtvS1-2* double mutant was comparable to the individual mutants (Fig. 2k). In contrast, *cas9* RNA levels were not significantly decreased in the individual mutants in comparison to the parental strain (Supplementary Fig. 2h-i). Interestingly, despite being only enriched in the pull-down with the *cas12* promoter and stationary phase lysate, *mtvS1* RNA levels were comparable in both growth phases while *mtvS2* RNA levels were significantly higher in stationary phase (Supplementary Fig. 2j-k, Supplementary Data 1, tab C and F). Fluorescence microscopy suggests the possibility of higher proteins level for MtvS1-mCherry2 in stationary phase, however no increase of protein levels was observed with LC-MS/MS (Supplementary Fig. 2l, Supplementary Data 1, tab A). It is possible that the fluorescence tag affected MtvS1 aggregation and/or stability properties^61–63^.

Next, we assessed whether MtvS1-2 dependent regulation of the type V-A CRISPR-Cas locus is important for plasmid restriction. Plasmid transfer into recipients with *mtvS1* and/or *mtvs2* deletions carrying a targeting spacer was not detected, the same as in the case of conjugation into parental recipients (Supplementary Fig. 2m-n). This result prevented us from determining whether the mutations affect plasmid targeting. Therefore, we decided to increase our level of detection of transconjugants to be able to detect differences in anti-plasmid CRISPR immunity, if any. First, we deleted the two restriction enzyme endonucleases, *FTN_0710* and *FTN_1487*, from the parental and mutant strains, which contribute to a low conjugation efficiency. Additionally, we decreased Cas12’s targeting activity by introducing mutations in the seed sequence of the protospacer^45^. Since the presence of a single target with a seed sequence mutation completely eliminated immunity, we integrated two protospacers into the conjugative plasmid, matching spacers 3 and 5 of the native type V-A CRISPR array and containing a mismatch at position 3 within each seed sequence (Fig. 2l). In this condition, the conjugation efficiency of a non-target plasmid into the parental strain was increased from 10^-5^/10^-4^ to ∼10^-3^, and the transfer of a target plasmid produced detectable transconjugants, but with a reduction in conjugation efficiency of one order of magnitude compared to the non-target control (Fig. 2l). In contrast, plasmid targeting did not affect conjugation efficiency into *mtvS1-2 nonsense* mutant recipients (Fig. 2l), a result that indicates that these genes are important for the type V-A CRISPR immune response, presumably due to their effects on *cas12* regulation. Interestingly, conjugation efficiency of the non-target plasmid into *mtvS1-2 nonsense* mutant recipients was reduced by an order of magnitude when compared to the parental strain, a finding that suggests the involvement of these genes in the regulation of other genes outside of the type V-A *CRISPR-cas* operon (see below) that affect conjugation, possibly involved in the generation of cell wall structures.

In summary, our results indicate that FTN_1238 and FTN_1519 are modulators of the type V-A CRISPR-Cas system expression in stationary phase, and therefore we renamed these genes *mtvS1* and *mtvS2*.

### MtvS1 and MtvS2 modulate expression of numerous genes in stationary phase

To investigate if MtvS1 and MtvS2 act on other genes beyond the type V-A operon, we performed RNA-seq of the *mtvS1* and *mtvS2 nonsense* mutants using RNA extracted from bacteria grown to exponential and stationary phase (Supplementary Data 1, tabs C-H). Comparison of RNA levels revealed that expression of the type V-A CRISPR locus is comparable in both growth conditions in these mutants, a result that confirms the requirement of MtvS1 and MtvS2 for increased expression of this locus in stationary phase (Fig. 3a-b Supplementary Data 1, tabs C-H). Intriguingly, principal component analysis and clustering of genes with similar expression patterns revealed differences between the parental strain and the mutants in particular in stationary phase samples (Fig. 3c-d, Supplementary Fig. 3a-b). In exponential phase, only a few genes with differential expression patterns between the parental strain and the two mutants were observed (Supplementary Data 1, tab C and F), GSE analysis suggested that in the *mtvS1 nonsense* mutant, downregulated genes are involved in cell envelope biogenesis and tRNA modification while genes involved in protein unfolding are upregulated (Supplementary Data 1, tabs C and E). For the *mtvS2 nonsense* mutant, no significantly down regulated pathways in exponential phase were detected, while genes involved in energy production and amino acid and coenzyme transports were upregulated (Supplementary Data 1, tabs F and H). Differences between the parental strain and the *mtvS1 nonsense* mutant were more profound than with the *mtvS2 nonsense* mutant in stationary phase, with more genes being significantly differentially expressed (Supplementary Data 1, tab C and F). For the *mtvS1 nonsense* mutant, especially genes involved in energy production and conversion as well as cell envelope biogenesis genes were negatively affected while transport genes were upregulated (Supplementary Data 1, tabs C and E). In general, many of the genes were affected more severely than *cas12* (Supplementary Data 1, tab C). On the other hand, for the *mtvS2 nonsense* mutant, different transport genes were downregulated while genes involved in translation were upregulated (Supplementary Data 1, tabs F and H). Interestingly, the expression of the *Francisella* pathogenicity island encoding the type VI secretion system, fundamental for this organism’s virulence^64,65^, a pantothenate biosynthesis operon, an operon containing DNA repair protein RecO and the ATP-dependent proteasome HlsVU are decreased in both mutants.

**Figure 3.**
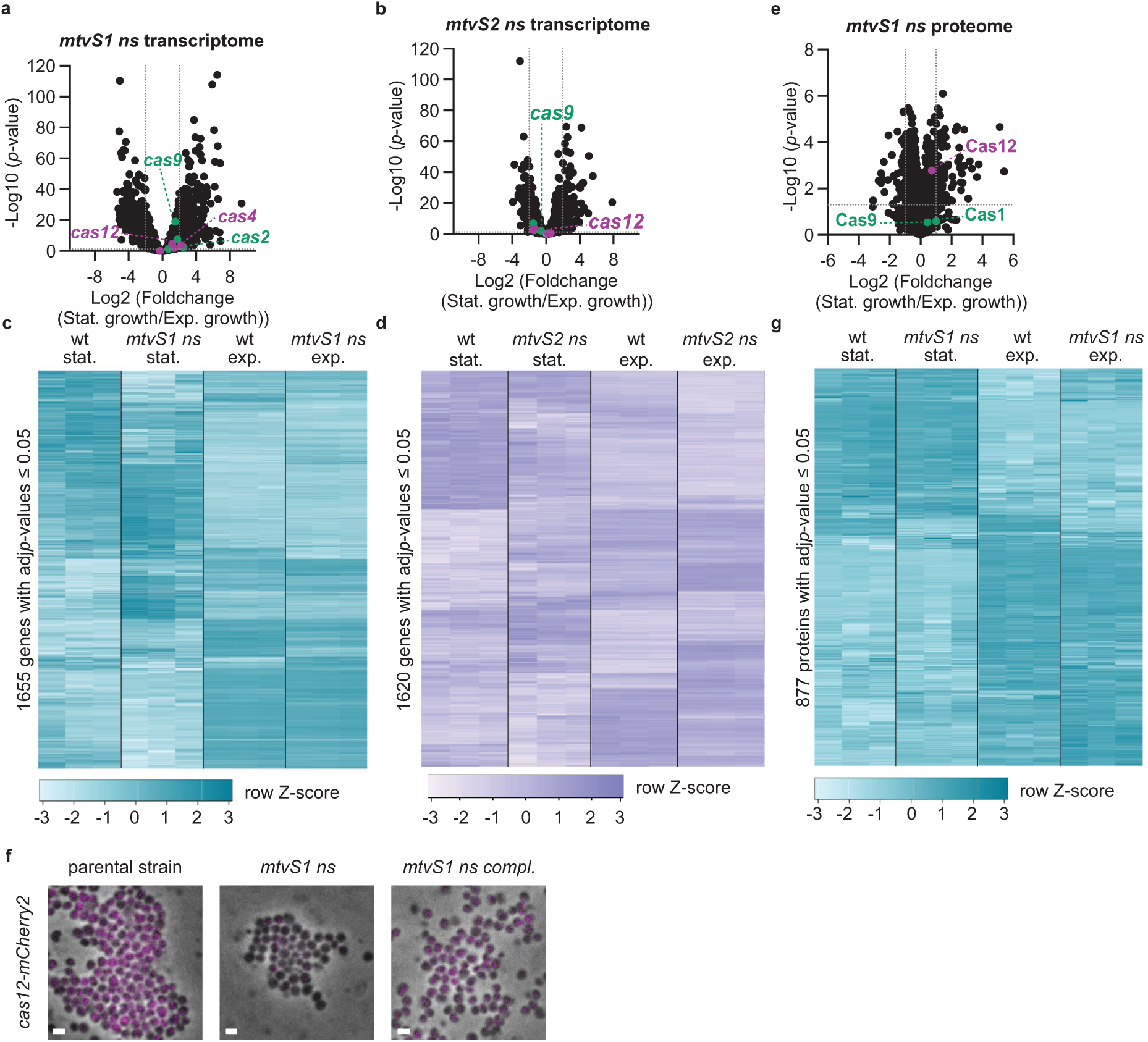
MtvS1 and MtvS2 modulate expression of numerous genes in stationary phase. **a-b** RNA-seq data of differentially expressed genes between stationary and exponential phase in *mtvS1 nonsense* **(a)** and *mtvS2 nonsense* **(b)** mutants. Green: *type II-B CRISPR-Cas* genes. Magenta: *type V-A CRISPR-Cas* genes. Dotted horizontal line: *p*-value of 0.05 (Wald test). Dotted vertical line: 4-fold difference. n = 3 biological replicates. **c-d** Clustered heatmaps of genes with an adj*p*-value ≤ 0.05 (Wald test with Benjamini-Hochberg correction) in RNA-seq data set showing differential expression of several genes in exponential (exp) and stationary (stat) phase between parental strain (wt) and *mtvS1 nonsense* **(c)** and *mtvS2 nonsense* **(d)** mutants. Z-scores represent the standard deviation of the actual value from the mean value calculated for each gene across samples. n = 3 biological replicates. **e** Proteomics data of differentially expressed proteins between stationary and exponential phase for the *mtvS1 nonsense* mutant. Green: type II-B CRISPR-Cas proteins. Magenta: type V-A CRISPR-Cas proteins. Dotted horizontal line: *p*-value of 0.05 (two-sample *t*-test). Dotted vertical line: 2-fold difference. n = 3 biological replicates. **f** Fluorescence microscopy of Cas12-mCherry2 in parental strain, *mtvS1 nonsense* mutant and *mtvS1 nonsense* mutant with wild-type *mtvS1* at Tn7 insertion site. *mtvS1 nonsense* mutant has decreased fluorescence intensity in stationary phase compared to the parental strain and rescued mutant. Merge of phase contrast and RFP channel in 13 x 13 μm fields of view are shown. Scale bars represent 1 μm. Representative images are shown (n = 2 biological replicates). Source data are provided as a Source Data file. **g** Clustered heatmap of proteins with an adj*p*-value < 0.05 (two-sample *t*-test with FDR correction (FDR = 0.05)) in proteomics data set showing differential expression of several proteins in exponential (exp) and stationary (stat) phase between parental strain (wt) and *mtvS1 nonsense* mutant. Z-scores represent the standard deviation of the actual value from the mean value calculated for each gene across samples. n = 3 biological replicates.

To find the putative -35 sequence associated with MtvS1-2 regulation, we analyzed the promoter sequences of the operons that were significantly downregulated in stationary phase in either of the two mutants, compared to the parental strain (Supplementary Data 1, tab E and H). Given that some of these genes may not be directly regulated by these factors (their expression could be a response to change in other factors that are directly regulated by MtvS1/2), we decided to filter the promoter sequences for the presence of the canonical -10 sequence (TATAAT), which is present in the *cas12* promoter used as bait to identify MtvS1 and MtvS2 (Fig. 2b, Supplementary Data 1, tab J). We found that 61 % of the promoters of downregulated genes (34/56) encoded a TATAAT sequence. In contrast, only 35 % of promoters (14/37) of upregulated operons in stationary phase contained the canonical -10 element (Supplementary Data 1, tab E and H). We believe that this enrichment is in agreement with the hypothesis that the canonical -10 sequence is important for MtvS1-2 dependent gene expression. Finally, we aligned the filtered promoters and detected a putative -35 sequence starting with a highly conserved A (Supplementary Fig. 3c).

Last, to corroborate the observed changes in transcription, we performed a proteomic analysis of the parental and *mtvS1 nonsense* mutant strains in both growth phases. We focused on MtvS1 rather than MtvS2 since other homologs exist in several γ-proteobacteria. The increase in Cas12 protein levels observed in the parental strain when cells entered stationary phase, was decreased in the absence of MtvS1 (Fig. 1b and 3e, Supplementary Fig. 3d, Supplementary Data 1, tab A). Although the decrease of Cas12 levels in the mutant strain is not statistically significant, this result is consistent with the decrease in the Cas12-mCherry2 fluorescence observed using microscopy (Fig. 3f). In spite of this decrease, microscopy showed residual fluorescence signal in the *mtvS1 nonsense* mutant, which was also confirmed by whole-cell proteomics results that revealed a small but detectable increase of Cas12 protein levels in stationary phase compared to exponential growth in this mutant (Fig. 3e-f, Supplementary Fig. 3d, Supplementary Data 1, tab A). This result provides a possible explanation for the high Cas12-dependent plasmid targeting levels detected for the *mtvS1 nonsense* mutant strain (Supplementary Fig. 2m). Other, statistically significant, changes in protein levels were observed in the absence of MtvS1 (Fig. 3e and g, Supplementary Fig. 3d). The GSE analysis suggests an upregulation of the UMP biosynthesis proteins in exponential growth phase in the *mtvS1 nonsense* mutant. In stationary growth phase, in the absence of MtvS1 the levels of proteins participating in bioenergetic processes were significantly decreased and some cell envelope proteins were more abundant, compared to the parental strain. (Supplementary Data 1, tab B). These changes in membrane and cell wall components, also detected in RNA-seq experiments, may contribute to the decrease in conjugation efficiency of the *mtvS1-2* double mutant recipient cells (Fig. 2l).

### Quorum sensing signals participate in the regulation of the type V-A CRISPR-Cas locus through MtvS1

Given that the *Francisella* type V-A CRISPR-Cas system is upregulated in stationary phase, when cell density is high, we investigated whether not only the stationary condition but also cell density may affect expression of the type V-A CRISPR locus. To test this hypothesis, we used fluorescence microscopy and exponentially growing cells in microfluidic plates. We observed that indeed Cas12-mCherry2 is expressed within densely populated areas but not in less dense regions present in the same flow chamber (Fig. 4a, Supplementary Movie 1-2 Supplementary Data 2). This density-dependent regulation of Cas12-mCherry2 was abrogated in *mtvS1 nonsense* mutant bacteria, a finding that confirmed that MtvS1 is required for this phenotype (Fig. 4b, Supplementary Movie 3-4, Supplementary Data 2)^60^. These results led us to hypothesize that quorum sensing signals, which are secreted to the medium and reach a high enough concentration when bacterial cultures reach high density, could play a role in regulating the type V-A CRISPR-Cas system. Therefore, we tested if the addition of spent overnight medium could increase Cas12-mCherry2 expression in exponential cultures of *F. novicida* U112. We replaced the medium of exponentially growing cells with spent overnight medium (filtered with 0.2 µm filters) in microfluidic plates and monitored individual cells by microscopy. We observed changes in cell shape from rod to coccus, as well as an increase in Cas12-mCherry2 fluorescence (Fig. 4c, Supplementary Movie 5, Supplementary Data 2).

**Figure 4.**
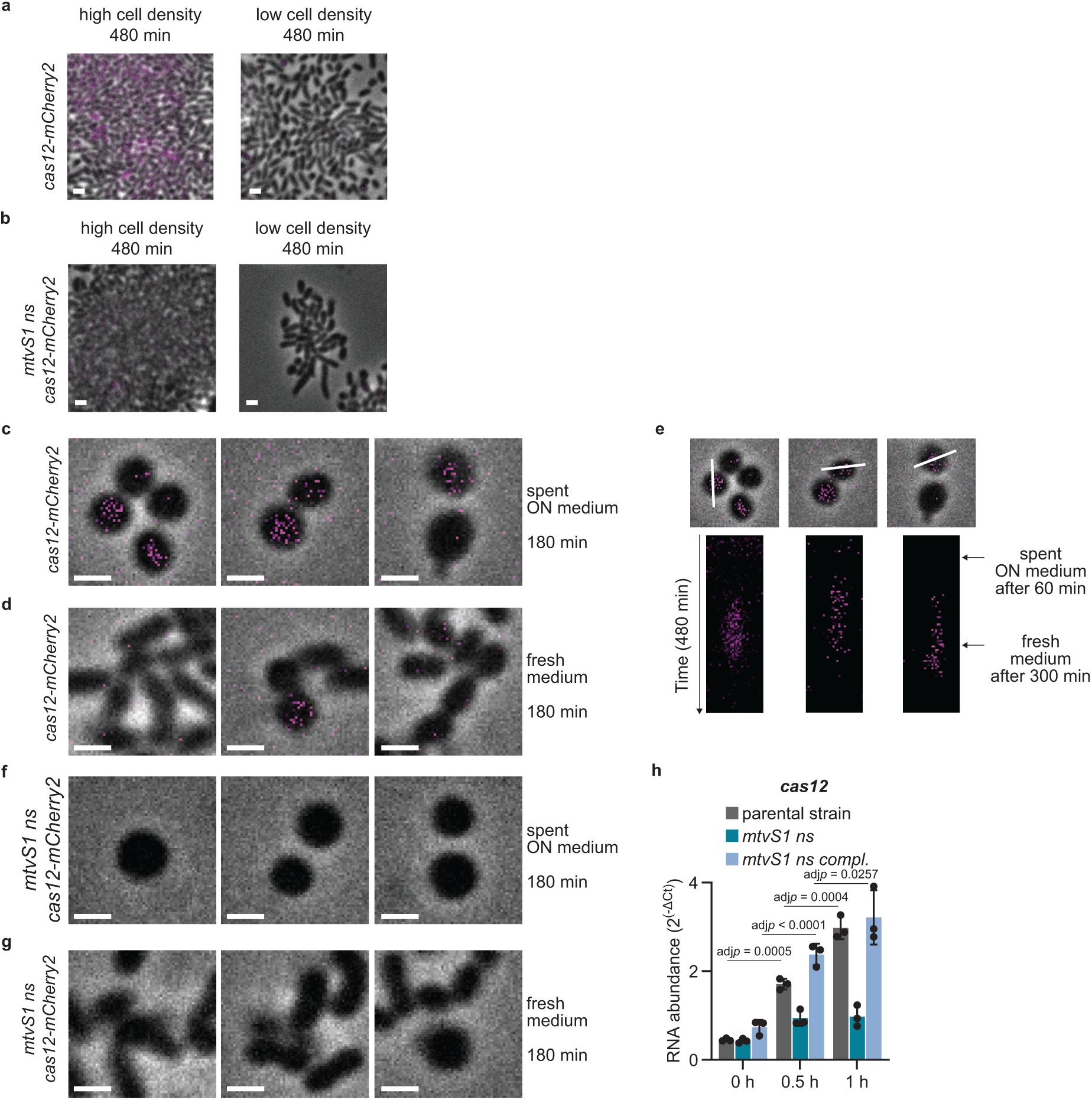
MtvS1 modulates *cas12* expression in a cell-density dependent manner. **a** Fluorescence microscopy of Cas12-mCherry2 in the parental strain in a microfluidic plate. BHI was flown through for 8 h. Merge of phase contrast and RFP channel in 13 x 13 μm fields of view are shown. Same contrast settings as in **b** and **Supplementary Movies 1-4**. Scale bars represent 1 μm. Representative images are shown (n = 3 biological replicates). Source data are provided as a Source Data file. **b** Fluorescence microscopy of Cas12-mCherry2 in the *mtvS1 nonsense* mutant in a microfluidic plate. BHI was flown through for 8 h. Merge of phase contrast and RFP channel in 13 x 13 μm fields of view are shown. Same contrast settings as in **a** and **Supplementary Movies 1-4**. Scale bars represent 1 μm. Representative images are shown (n = 3 biological replicates). Source data are provided as a Source Data file. **c-d** Fluorescence microscopy of Cas12-mCherry2 in the parental strain in a microfluidic plate. BHI was flown through for 1 h before switching to spent overnight medium of parental strain for 4h. Afterwards, fresh BHI was flown through for 3 h. **(c)** Three examples of 240 min time point (180 min after switching to spent ON medium) and **(d)** three examples of 480 min (180 min after switching back to fresh BHI) are shown. Merge of phase contrast and RFP channel in 4 x 4 μm fields of view are shown. Same contrast settings as in **e-g**, **Supplementary Fig. 4d** and **Supplementary Movies 5, 7-8**. Scale bars represent 1 μm. Representative images are shown (n = 3 biological replicates). Source data are provided as a Source Data file. **e** Examples of **c** and **d** were used to create time-resolved kymographs of RFP channel. Merge of phase contrast and RFP channel is shown in the upper images. White lines show the axis along which the kymographs of RFP channel were made (lower images). Same contrast settings as in **c-d** and **f-g**, **Supplementary Fig. 4d** and **Supplementary Movies 5, 7-8**. Representative images are shown (n = 3 biological replicates). Source data are provided as a Source Data file. **f-g** Fluorescence microscopy of Cas12-mCherry2 in the *mtvS1 nonsense* mutant in a microfluidic plate. BHI was flown through for 1 h before switching to spent overnight medium of parental strain for 4h. Afterwards, fresh BHI was flown through for 3 h. **(f)** Three examples of 240 min time point (180 min after switching to spent ON medium) and **(g)** three examples of 480 min (180 min after switching back to fresh BHI) are shown. Merge of phase contrast and RFP channel in 4 x 4 μm fields of view are shown. Same contrast settings as in **c-e**, **Supplementary Fig. 4d** and **Supplementary Movies 5, 7-8**. Scale bars represent 1 μm. Representative images are shown (n = 2 biological replicates). Source data are provided as a Source Data file. **h** Determination of mRNA levels for *cas12* after switching to spent overnight medium by RT-qPCR. Samples were taken 0 h, 0.5 h and 1 h after medium change. Grey bars: parental strain. Turquoise bars: *mtvS1 nonsense* mutant. Light blue bars: *mtvS1 nonsense* mutant with wild-type *mtvS1* at Tn7 insertion site. Means with standard deviations are displayed (n = 3 biological replicates). Significant differences were determined with a two-way ANOVA and Tukey’s multiple comparison test (95 % confidence interval). Source data are provided as a Source Data file.

Fresh medium reversed these changes, with cocci converting into rods and resuming growth, and a decrease in Cas12-mCherry2 fluorescence (Fig. 4d). Time-resolved kymographs showed that the mCherry2 signal increased and decreased within 1-2 hours after each media change (Fig. 4e). Since the mCherry2 signal is rather low from the endogenously expressed Cas12-mCherry2, we repeated the experiment expressing mCherry2 under the control of the *cas12* promoter from a plasmid to obtain better visualization of the fluorescence signal than from a single chromosomal locus. Indeed, we detected higher levels of mCherry2 fluorescence, which followed similar dynamics when challenged with spent medium as the endogenously expressed Cas12-mCherry2 (Supplementary Fig. 4a-c, Supplementary Movie 6, Supplementary Data 2), a result that validated the use of the chromosomally encoded *cas12-mCherry2* strain. When exposed to overnight spent medium, *mtvS1 nonsense* mutant cultures also suffered rod-to-coccus shape changes, without detectable Cas12-mCherry fluorescence (Fig. 4f-g, Supplementary Movie 7, Supplementary Data 2). Heat-treatment of the spent overnight medium eliminated its activity on exponentially growing bacteria (Supplementary Fig. 4d, Supplementary Movie 8, Supplementary Data 2). These results were confirmed by RT-qPCR, using RNA extracted from parental strain and *mtvS1 nonsense* mutant cells before, 0.5 h and 1 h after switching to spent overnight medium and probed for *cas12* and *cas9* RNA levels. While *cas12* RNA levels increased in a MtvS1-dependent manner over 1 h after the medium change, *cas9* RNA levels did not significantly change (Fig. 4h and Supplementary Fig. 4e). Together these results suggest that *F. novicida* U112 may integrate quorum sensing signals into the MtvS1 dependent regulation of Cas12 expression.

### MtvS1 and MtvS2 interact with RNA polymerase

To investigate the mechanism by which MtvS1 and MtvS2 modulate *cas12* transcription, we heterologously expressed and purified His_6_-tagged versions of the proteins in *Escherichia coli* (Supplementary Fig. 5a-b) and tested whether they bind directly to the *cas12* promoter. To control that the addition of the His_6_-tag did not alter their functions, we integrated the *His_6_*-tag into the chromosome of *F. novicida* U112 for both genes, determined *cas12* RNA levels by RT-qPCR, and found that these strains displayed a similar *cas12* transcription regulation as the wild-type strain (Supplementary Fig. 5c-d). Electrophoretic mobility shift assays (EMSA) showed binding of MtvS2-His_6_ to both *cas12* promoter DNA as well as an intragenic *cas12* sequence which we used as a non-specific control sequence (Fig. 5a). On the other hand, we were unable to detect DNA binding for MtvS1-His_6_ in the tested conditions. Addition of MtvS1-His_6_ did not alter the DNA shift pattern of MtvS2-His_6_ (Fig. 5b). We performed Western blot analysis of MtvS1-His_6_ expressed in *Francisella* to look for changes in electrophoretic mobility in stationary phase. Such changes could indicate amino acid modifications not present in the heterologously expressed protein that may affect its DNA binding activity. The mobility of MtvS1-His_6_, however, did not change in the different growth phases (Supplementary Fig. 5e), a result that suggests lack of major protein modifications.

**Figure 5.**
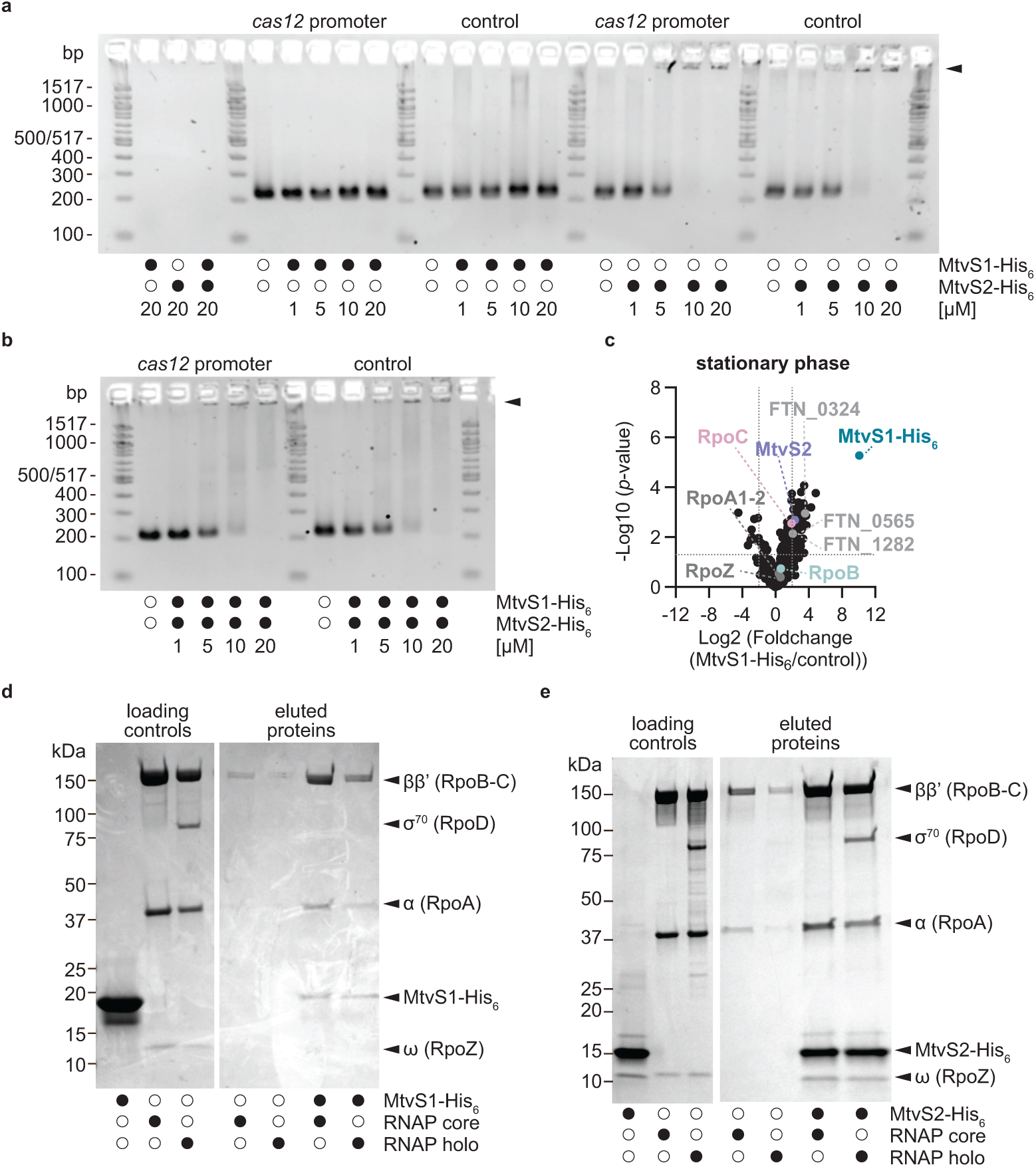
MtvS1 and MtvS2 interact with β’-subunit of RNA polymerase. **a-b** Electrophoretic mobility shift assay of *cas12* promoter and intragenic control sequences in the presence of increasing concentrations of MtvS1-His_6_ and MtvS2-His_6_ proteins individually **(a)** or combined **(b)**. Arrow indicates shifted DNA. Representative images are shown (n = 3 for MtvS1-His_6_ and n = 1 for MtvS2-His_6_). Source data are provided as a Source Data file. **c** Pull-down of proteins from stationary phase lysate of the *mtvS1 nonsense* mutant binding specifically to MtvS1-His_6_. Turquoise circle: MtvS1-His_6_. Grey circles: DNA-binding proteins (FTN_0565, and FTN_1282) and putative anti-anti sigma factor (FTN_0324). Purple cirlce: MtvS2. Pale cyan circle: RpoB. Pale pink circle: RpoC. Dark grey circles: RpoA1, RpoA2 and RpoZ. Dotted horizontal line: *p*-value of 0.05 (two-sample *t*-test). Dotted vertical line: 4-fold difference. n = 3 biological replicates. **d-e** SDS-PAGE gel of *E. coli* RNA polymerase core enzyme and holoenzyme pull-downs with MtvS1-His_6_ **(d)** or MtvS2-His_6_ **(e)**. 20 µM of MtvS1-His_6_ or MtvS2-His_6_ was incubated with 400 units/ml E. coli RNA polymerase core enzyme or holoenzyme and then pulled down with magnetic beads. σ^70^ was not pulled down when *E. coli* RNA polymerase holoenzyme was incubated with MtvS1-His_6_. Representative images are shown (n = 3). Source data are provided as a Source Data file.

Therefore, our findings suggest that MtvS1 associates with the *cas12* promoter DNA through interactions with other proteins. We then used purified MtvS1-His_6_ protein as bait to pull down potential partners from lysates of *mtvS1 nonsense* mutant cells grown to stationary phase. LC-MS/MS analysis of the proteins pulled-down with recombinant MtvS1-His_6_ revealed that two DNA-binding proteins (FTN_0565 and FTN_1282), a putative anti-anti sigma factor (FTN_0324) as well as MtvS2 were among the significantly enriched proteins suggesting that MtvS1 and MtvS2 interact (Fig. 5c, Supplementary Data 1, tab K). However, while deletion of *FTN_0565* (containing a Lambda repressor-like DNA binding domain) was not possible, deletion of *FTN_ 0324* (putative anti-anti sigma factor with a STAS domain) and *FTN_1282* (LysR family transcriptional regulator) did not affect Cas12-mCherry2 expression (Supplementary Fig. 5f).

Another protein that was significantly enriched in the pull-down was RpoC (FTN_1567), the β’ subunit of RNA polymerase (Fig. 5c, Supplementary Data 1, tab K). To probe for an interaction between MtvS1 and RNA polymerase (RNAP), a pull-down with purified MtvS1-His_6_ and commercially available *E. coli* RNAP core enzyme and holoenzyme (RNAP core enzyme in complex with σ^70^) was performed. We found that *E. coli* RNAP core enzyme was pulled down with MtvS1-His_6_ regardless of whether σ^70^ was present or not (Fig. 5d). Since MtvS2 was pulled down with MtvS1-His_6_, we tested whether MtvS2-His_6_ also interacts with RNAP. Interestingly, we found that MtvS_2_-His_6_ bound not only core, but also the RNAP:σ^70^holoenzyme (Fig. 5e).

To investigate how MtvS1 and MtvS2 interact with the RNA polymerase, we used AlphaFold3 to model *Francisella* RNAP core enzyme together with MtvS1 and MtvS2 (Fig. 6a, Supplementary Fig. 5g)^66^. This analysis rendered a high-confidence structural prediction showing the binding of MtvS1 to the coiled-coil domain of the β’ subunit, the main σ^70^ binding interface^67^. A list of these predicted interactions between RpoC and MtvS1 can be found in Supplementary Table 1. Structural predictions using *E. coli* RNAP showed that the β’ coiled-coil domain is predicted to dock into a pocket of MtvS1 similar to what was found for the 2.2 region of σ^70^ (Fig. 6b, Supplementary Fig. 5g-h)^67^. Therefore, our results suggest that MtvS1 binds to the β’ subunit of RNAP likely at the β’ coiled-coil domain. For MtvS2, the AlphaFold3 model is less certain (Supplementary Fig. 6b), but nevertheless predicts interactions with both the *Francisella* RNA polymerase β’ subunit and MtvS1, corroborating our pull-down results with MtvS1-His_6_ (Fig. 5c). Detailed analysis of MtvS2 interactions suggests specific interactions with the β-flap of the β-subunit (Fig. 6c).

**Figure 6.**
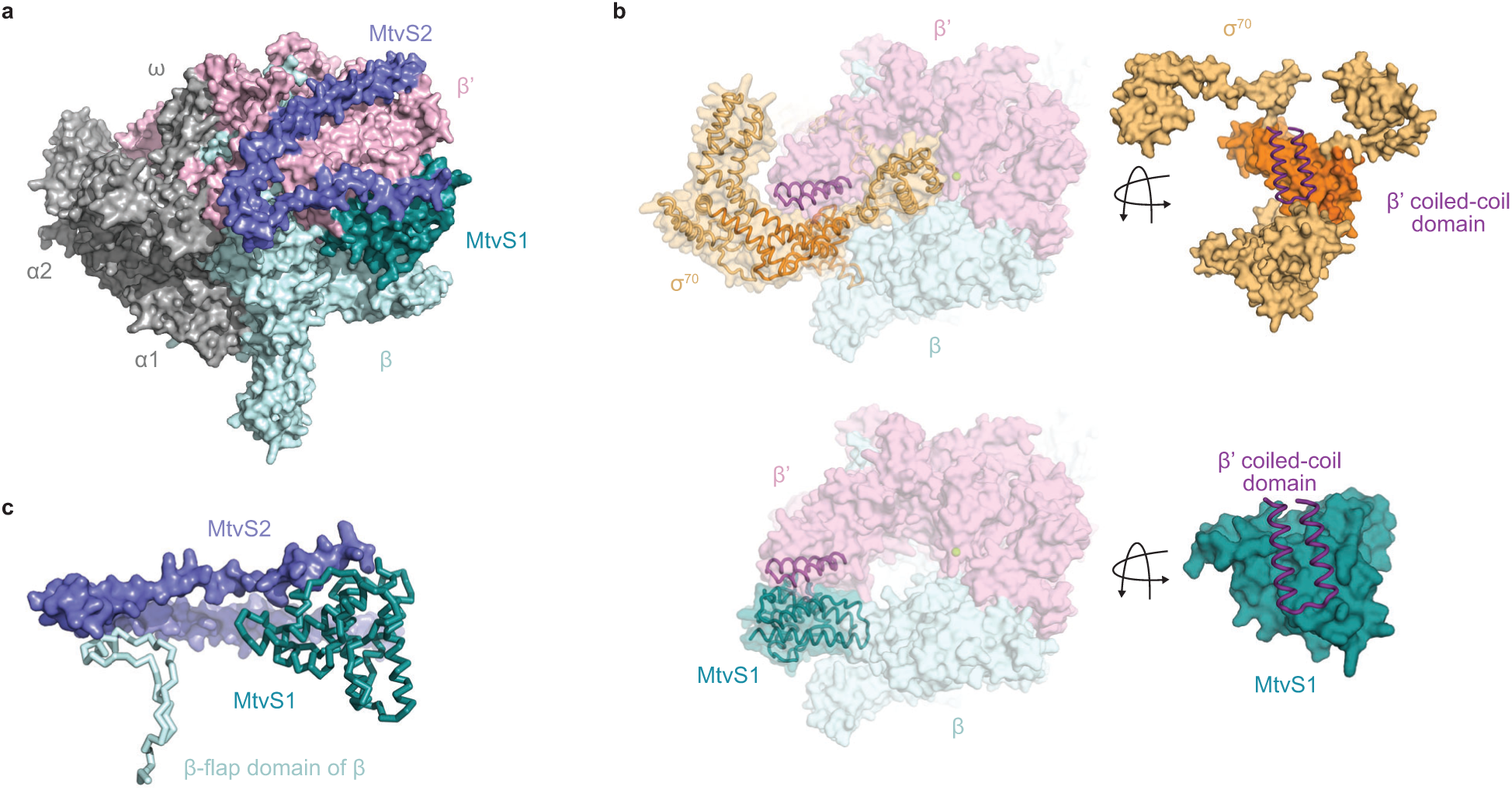
Structural predictions of MtvS1 and MtvS2 interacting with RNA polymerase. **a** AlphaFold 3 prediction of MtvS1 (A0A6I4RTR9, turquoise) and MtvS2 (A0Q819, purple) with *F. novicida* U112 RNA polymerase (RpoA1: A0Q4K8 (dark grey), RpoA2: A0Q7R6 (dark grey), RpoB: A0Q867 (pale cyan), RpoC: A0Q866 (pale pink) and RpoZ: A0Q5J3 (dark grey)). **b** AlphaFold 3 predictions of *E. coli* RNA polymerase (RpoB: P0A8V2 (pale cyan) and RpoC: P0A8T7 (pale pink)) with σ^70^ (P00579, gold) or MtvS1 (A0A6I4RTR9, turquoise). Coiled-coil domain of β’ RNAP subunit is shown in purple and catalytic magnesium ion is shown in light green. Close-ups show interface between coiled-coil domain of β’ RNAP subunit (purple) and domain 2 and 3 of σ^70^ (orange) or MtvS1 (turquoise). **c** Close-up of **(a)** shows predicted interface between MtvS2 (purple) and β-flap domain of β RNAP subunit (pale cyan) and MtvS1 (turquoise).

Given the above results, we hypothesized that MtvS1 and MtvS2 may be a noncanonical split alternative sigma factor similar to the smaller extracytoplasmic function (ECF) and phage sigma factors^68–70^. Since domain 2 of σ^70^ is also involved in recognition and melting of the -10 promoter element^71^, it is possible that MtvS1 has a similar role. However, the protein is too small to fulfill the function of domain 4 of σ^70^, which interacts with the β-flap and recognizes the -35 promoter element^72,73^. Therefore, we propose that MtvS2 may be a co-activator of MtvS1, similar to Gp55 and Gp33, responsible for the transcription of phage T4 late phage genes^74–76^. Yet, transcription from the *cas12* promoter, was not stimulated by neither MtvS1-His_6_ or MtvS2-His_6_ individually, nor together, nor with σ^70^ (Supplementary Fig. 5i-j).

### YgfB is a MtvS1 homolog that affects expression of the type I-E CRISPR-Cas system in E. coli

We searched for MtvS1 homologs in 99,999 bacterial genomes and found them present in ∼ 40 % of the organisms (Supplementary File, tab L-O). We also analyzed the distribution of CRISPR-Cas systems within genomes containing MtvS1 homologs and observed that they co-occur more often with type I and type IV CRISPR-Cas systems (Fig. 7a, Supplementary Data 1, tab L). *E. coli* harbors the MtvS1 homolog YgfB (b2909), which is 27.2 % identical at the amino acid sequence level (Fig. 7b) as well as a type I-E CRISPR-Cas system^6^ (Supplementary Fig. 6b). Structural alignment of the two protein models suggests a highly conserved structure (RSMD value: 1.396 Å over 788 atoms, Fig. 7c, Supplementary Fig. 6a), with many of the MtvS1 residues that are predicted to interact with β’ coiled-coil domain conserved in YgfB (Fig. 7b). To investigate whether YgfB affects the transcription of the CRISPR locus, we performed RT-qPCR of the *cas6e*, *cas7* and *cas8e* transcripts, which encode proteins of the Cascade complex involved in crRNA processing and target recognition^77^. We measured RNA levels in both the parental strain and a *ygfB::ampR* mutant that were grown to exponential and stationary phase. Similarly to the results we obtained for the type V-A CRISPR-*cas* locus of *F. novicida* U112, all transcripts tested were increased in stationary phase cells, compared to exponential growth, for the parental strain (Fig. 7d-f). This increase, however, significantly less pronounced in the *ygfB::ampR* mutant, but was restored in the presence of plasmids expressing YgfB or MtvS1 (using their native promoters) (Fig. 7d-f), a result that suggests that YgfB and MtvS1 have similar functions.

**Figure 7.**
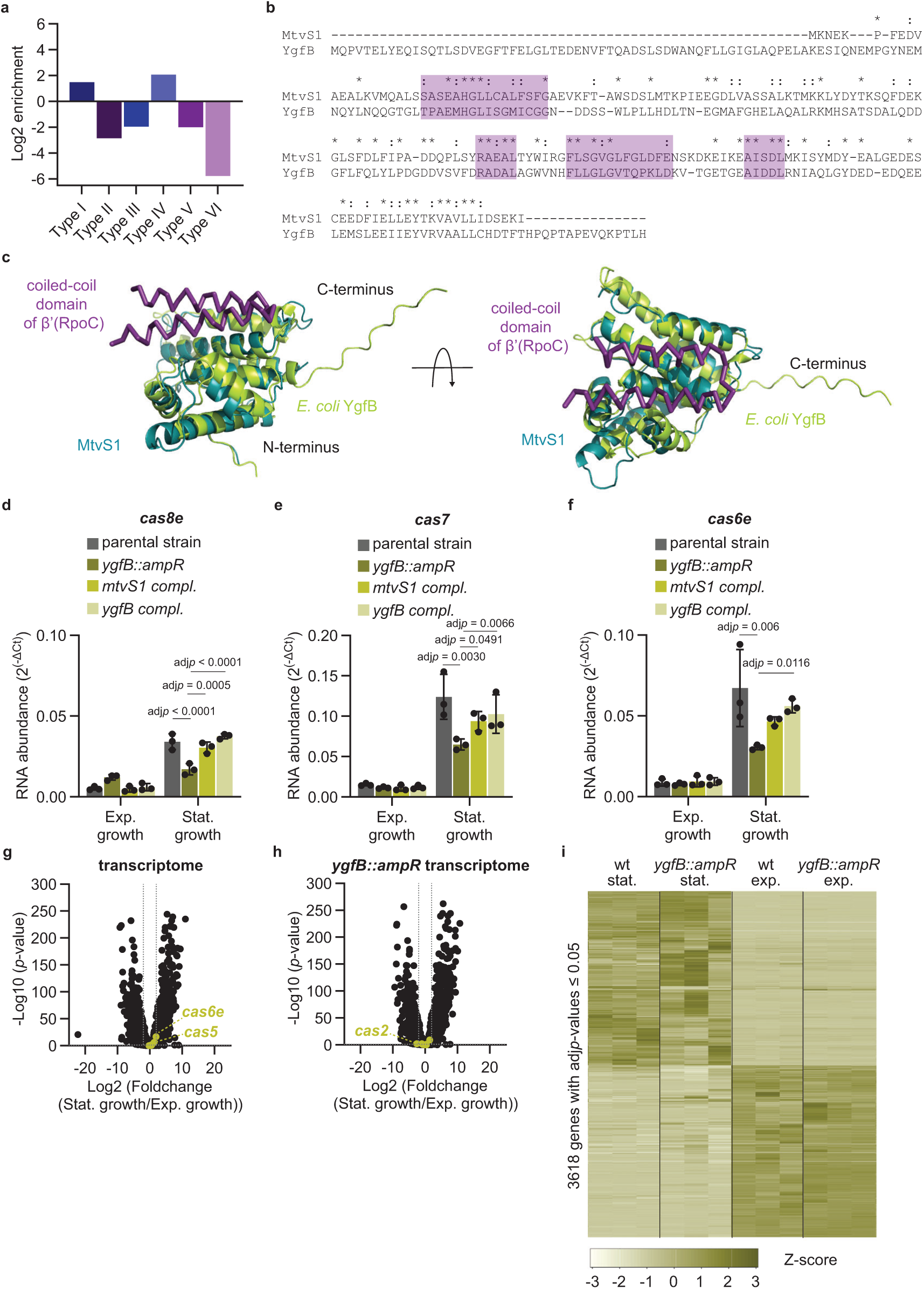
MtvS1 homolog YgfB affects transcription of the *type I-E CRISPR-Cas* system in *E. coli*. **a** Calculated enrichment score for the prevalence of different types of CRISPR-Cas systems in genomes containing MtvS1 homologs. Shades of blue: class I CRISPR-Cas systems. Shades of purple: class II CRISPR-Cas systems. **b** Protein sequences alignment of MtvS1 and *E. coli* YgfB. Identical and similar residues are indicated by (*) and (:), respectively. Purple boxes indicate the sequences involved in the predicted interaction with the coiled-coil domain of β’. **c** AlphaFold3 model of *E. coli* YgfB (light green, A8A450) is aligned to MtvS1 (turquoise, A0A6I4RTR9, RSMD value: 1.396 Å over 788 atoms) together with coiled-coil domain of β’ subunit (purple, A0Q867). **d-f** Determination of mRNA levels for *cas8e* **(d)**, *cas7* **(e)** and *cas6e* **(f)** by RT-qPCR. Grey bars: parental strain with empty plasmid. Olive green bars: *yfgB::ampR* mutant with empty plasmid. Light green bars: *ygfB::ampR* mutant with *F. novicida U112 mtvS1* expressed from plasmid. Pale green bars: *ygfB::ampR* mutant with *E. coli ygfB* expressed from plasmid. Means with standard deviations are displayed (n = 3 biological replicates). Significant differences were determined with a two-way ANOVA and Tukey’s multiple comparison test (95 % confidence interval). Source data are provided as a Source Data file. **g-h** RNA-seq data of differentially expressed genes between stationary and exponential phase in *E. coli* parental strain **(g)** and *ygfB::ampR* mutant **(h)**. Light green: *type I-E CRISPR-Cas* genes. Dotted horizontal line: *p*-value of 0.05 (Wald test). Dotted vertical line: 4-fold difference. n = 3 biological replicates. **i** Clustered heatmaps of genes with an adj*p*-value < 0.05 (Wald test with Benjamini-Hochberg correction) in RNA-seq data set showing differential expression of several genes in exponential (exp) and stationary (stat) phase between parental strain (wt) and *ygfB::ampR* mutant. Z-scores represent the standard deviation of the actual value from the mean value calculated for each gene across samples. n = 3 biological replicates.

To investigate whether YgfB, like MtvS1, affects the expression of additional genes beyond the *cas* operon, we compared RNA-seq data for the parental strain and *ygfB::ampR* mutant in both growth conditions (Supplementary Fig. 6c, Supplementary Data 1, tabs P-R). We found that two genes of the type I-E CRISPR-*cas* operon, *cas6e* and *cas5*, were upregulated in stationary phase (Fig. 7g, Supplementary Data 1, tab P). Compared to *cas* transcripts levels in the parental strain in stationary phase, RNA reads for all the genes in the operon, except *cas6e*, were decreased (Fig. 7h, Supplementary Data 1, tab P). Beyond the CRISPR-*cas* locus, other genes were differently expressed in the *ygfB::ampR* mutant (Fig. 7i, Supplementary Data 1, tab P). GSE analysis revealed stationary phase downregulation of a group of 15 poorly characterized genes (5 of which are encoded in cryptic prophage regions; *b1137*, *b1138*, *b2623*, *b4306*, *b4307*) as well as genes involved in ubiquinone biosynthesis (Supplementary Data 1, tab R). On the other hand, expression of carbohydrate transport and metabolism genes was increased in stationary phase in the *ygfB::ampR* mutant. We did not detect significantly up- or down-regulated pathways in exponential growth phase compared to the parental strain. Altogether our data showed that MtvS1 and YgfB not only share sequence and structural homology, but also participate in the regulation of *cas* genes and other genes in *F. novicida* U112 and *E. coli*, respectively, most likely through their interaction with RNA polymerase as an alternative sigma factor.

## Discussion

Here we compared the expression of the two native type II-B and type V-A CRISPR-Cas loci in *F. novicida* U112 in the absence of phage infection. We found that while the type II-B CRISPR-Cas system is continuously expressed throughout growth in liquid culture, the type V-A CRISPR-Cas system is significantly upregulated at high cell density and stationary growth phase. We believe that there are different benefits for *Francisella* behind this unique regulation. First, it is known that bacteria enhance their anti-phage defenses in high cell density environments when they are more vulnerable to phage predation^30,31,78–84^. Given that the type II-B system is constitutively expressed, the upregulation of the type V-A system will equip *F. novicida* U112 with additional defenses in stationary phase. It is also possible that Cas12 or other components of the type V-A CRISPR-Cas system have a role beyond immunity in stationary growth phase, analogous to the function of *F. novicida* U112 Cas9 in the regulation of lipoprotein expression^48–51^. Our proteomics and RNA-seq experiments showed drastic changes in transcription and translation in stationary phase, indicating the need to adapt to this growth condition, an adaptation in which Cas12 could play a yet unknown function.

Regarding the mechanisms behind *cas12* upregulation in stationary phase, we identified MtvS1 and MtvS2, proteins with previously unknown function in *F. novicida* U112, as factors required for the increase in the transcription of the type V-A CRISPR-*cas* locus. While these factors substantially contribute to this increase, anti-plasmid interference assays, fluorescence microscopy visualization of Cas12, and transcriptomics and proteomics data indicated presence of Cas expression in stationary phase even in the absence of MtvS1 and MtvS2. Therefore, we believe it is likely that *F. novicida* U112 encodes additional regulators that feed into the regulatory network for type V-A CRISPR-Cas system activation. An analysis of promoters of the downregulated operons in stationary phase in the *mtvS1* and *mtvS2 nonsense* mutants showed that more than half of these promoters encode the same -10 sequence as the *cas12* promoter (TATAAT). However, our study cannot distinguish whether these genes are directly regulated by MtvS1 and MtvS2 or whether they are up- or downregulated as part of a compensatory mechanism that takes place in the mutant strains.

We found that MtvS1 is predicted to interact with the β’ subunit of *F. novicida* U112 and *E. coli* RNA polymerase, through the binding of a coiled coil domain of the β’ subunit that has been shown to interact with σ^70^ (domain 2) and σ^32^ in *E. coli*^72,85–87^. Domain 2 of σ^70^ is also required for recognition and melting of the -10 promoter sequence^71^. Since *Francisella* only possesses two sigma factors^52^, σ^70^ and σ^32^, it is tempting to speculate that MtvS1 has a similar role in recognizing the -10 promoter sequence and initiating transcription. The broad transcriptional change, especially in stationary phase, caused by the disruption of MtvS1 supports a role for this protein as an alternative sigma factor. *In vitro*, however, MtvS1 neither alone nor together with σ^70^, was sufficient to stimulate transcription of the *cas12* promoter. In addition, structural predictions indicated that MtvS1 lacks the equivalent of domain 4 of σ^70^, which interacts with the β-flap to recognize the -35 promoter sequence^72,73^. On the other hand, MtvS2 is predicted to interact with the β-flap of RNAP, similarly to domain 4 of σ^70^, in the presence of MtvS1. It is possible that MtvS1-2 constitute a pair of split sigma factors, similarly to Gp55 and Gp33 in T4 phage which, together with the DNA clamp Gp45, initiate transcription of late viral genes by taking over the roles of domain 2 and domain 4 of σ^70^, respectively^74–76^. Given the non-specific DNA binding of MtvS2 *in vitro*, we propose that promoter selectivity and binding are possibly determined in the complex formed with RNA polymerase and additional factors such as MtvS1. However, we were not able to reconstitute transcription of the *cas12* promoter using MtvS2 alone or together with MtvS1, a result that suggests the existence of yet additional factors required for regulation. Alternatively, *in vitro* transcription assays could have failed due to the need of *Francisella*, and not *E. coli*, RNA polymerase. *Francisella* encodes two different α-subunits that form an α^1^α^2^ββ’ core enzyme^54,55^ and are evolutionarily distinct from other γ-proteobacteria, including *E. coli*, that harbor a single α-subunit gene and form an α_2_ββ’ core enzyme. Since α C-terminal domains can interact with promoter DNA elements, it is possible that *E. coli* RNAP cannot appropriately recognize the *cas12* promoter. In spite of these caveats, our results highlight that both MtvS1 and MtvS2 are novel factors that interact with RNA polymerase and modulate gene expression in *F. novicida* U112. According to structural predictions, while MtvS1 excludes the binding of the canonical sigma factor σ^70^, MtvS2 seems to occupy a more flexible space. We speculate that the advantage of having two separate proteins either replacing MtvS1-2 or working together with canonical sigma factors could enable a mix and match approach that would allow the fine tuning of gene expression to differentially change the transcription profile of *Francisella* in response to diverse conditions and environments.

While MtvS2 is a *Francisella* specific regulator, MtvS1 belongs to the YecA/YgfB-like protein family^60^. We found that MtvS1 homologs are particularly present in bacteria encoding type I and type IV CRISPR-Cas systems despite MtvS1 being important for the regulation of the type V-A CRISPR-Cas system in *Francisella*. This discrepancy may be explained by the fact that type V CRISPR-Cas systems are more prominent in *Spirochaetota*, *Cyanobacteriota*, *Planctomycetota* and *Verrucomicrobiota*, while Mtvs1 homologs are confined to γ-proteobacteria, which belong to the *Pseudomonadota* phylum^60,88^. Besides the structure of *H. influenzae* YgfB, little is known about the function of this family of proteins^89^. In *Pseudomonas aeruginosa*, YgfB increases β-lactam resistance through interactions with the antitermination factor AlpA that lead to the upregulation of AmpC expression^90,91^. We believe that it is unlikely that MtvS1 interacts with an anti-termination factor in *F. novicida* U112 since *cas12* transcription decreases in the *mtvS1 nonsense* mutant. *E. coli* YgfB is functionally redundant to MtvS1 and is important for transcription in the stationary phase, including transcription of the type I-E CRISPR-Cas system present in this organism. This result highlights the broader role that MtvS1/YgfB proteins play in modulating bacterial transcription in stationary phase, particularly of bacterial defense systems. Under laboratory conditions, the *E. coli* type I-E CRISPR-Cas system is inactive due to transcriptional repression by H-NS^19,20^. This repression can be relieved through LeuO or CRP^20,92^. It is currently unclear whether YgfB directly acts on its targets or through one of these regulators; although our RNA-seq data does not suggest changes in expression of *h-ns* (*b1237*), *leuO* (*b0076*) or *crp* (*b3357*) in the *ygfB::ampR* mutant. Since MtvS2 has no homologs in *E. coli*, it will be of interest to investigate whether additional factors interact with YgfB to modulate gene expression or whether the C-terminal tail of YgfB (not present in MtvS1) may play a similar role as MtvS2.

Finally, we found that *F. novicida* U112 cells reacted strongly to spent overnight medium, displaying a growth arrest, morphological changes and an increase in Cas12 expression, the latter also dependent on the presence of MtvS1. This behavior is a hallmark of quorum sensing^93^. Interestingly, *F. novicida* U112 lacks canonical quorum sensing genes^94^ and therefore our results suggest that the type V-A CRISPR-Cas system and MtvS1 could be part of a yet to be described quorum sensing circuit that monitors and responds to changes in cell density in this organism. Future studies will investigate the quorum sensing network and the exact mechanisms of how MtvS1 and MtvS2 promote transcription of *cas12* and other genes. Together with the characterization of the affected genes, elucidating these mechanisms in detail will not only be relevant for the understanding of phage defense strategies but also for the pathogenesis of *Francisella*.

## Methods

### Bacterial strains and growth conditions

*Francisella novicida* U112 (*F. novicida* U112) and derivative mutants were grown aerobically in brain heart infusion (BHI, BD Biosciences) broth or on BHI agar plates supplemented with 0.1 % L-cysteine (Sigma-Aldrich) and 50 µg/ml ampicillin (GoldBio) at 37 °C. F*. novicida* U112 strains carrying plasmids were grown with 15 µg/ml kanamycin (GoldBio).

To receive exponentially growing cultures, stationary phase cells were diluted 1:20 in fresh media and grown for 2-3 h until they reached an OD_600_ of 0.5-0.8 and had a rod-shaped morphology. Stationary phase cells were grown over night for at least 16 hours until they reached an OD_600_ of 3.5-4.5 and displayed a more coccus-like cell shape. Cell morphology was checked by microscopy.

*Escherichia coli* OneShotpir (*E. coli,* Thermo Fisher Scientific) and derivative strains were aerobically grown in Luria broth (LB, BD Biosciences) or on LB agar plates supplemented with 50 µg/ml kanamycin at 37 °C.

For conjugation *E. coli* JKE201 strain, 1.2 mM 2,6-Diaminopimelic acid (DAP, Sigma-Aldrich) was added to all growth media since this strain is auxotrophic for mDAP.

*E. coli* K-12 *substr*. BW25113 (Coli Genetic Stock Center at Yale University), the parental strain of the Keio collection^95^, and derivative strains were aerobically grown in LB or LB agar plates supplemented with 30 µg/ml chloramphenicol (GoldBio), 100 µg/ml ampicillin and/or 50 µg/ml kanamycin if required.

To receive exponentially growing cultures, stationary phase cells were diluted 1:500 in fresh media and grown for 2-3 h until they reached an OD_600_ of 0.5-0.8. Stationary phase cells were grown over night for at least 16 hours until they reached an OD_600_ of 4-4.5.

*E. coli* Rosetta 2 (DE3) (Novagen) containing the pET21 overexpression plasmid was grown aerobically in LB with 100 µg/ml ampicillin. The strain carrying the recombinant *mtvS2-His_6_* construct was grown at 30 °C.

*E. coli* BL21 (DE3) (Novagen) containing the pVS11 and pACYC-Duet1_*EcoRpoZ* expression plasmid was grown aerobically in LB with 100 µg/ml ampicillin and 34 µg/ml chloramphenicol to overexpress *E. coli* RNAP core enzyme.

*E. coli* BL21 (DE3) containing the pSAD1403 expression plasmid was grown aerobically in LB with 50 µg/ml kanamycin to overexpress *E. coli* σ^70^ factor.

Strains used for this study are listed in Supplementary Table 2.

### Bacterial mutagenesis

To generate chromosomal in-frame *F. novicida* U112 mutants, suicide vector pDMK3 was used ^96^. For fluorescent tags, an Ala-Ala-Ala-Gly-Gly-Gly linker was added and after the stop codon the 30 bp sequence of the gene was repeated to avoid downstream polar effects. For the His-tags only a Gly linker was added. In-frame deletions yielded in peptide scars containing the first and last 10 amino acids. Nonsense mutations were generated by introducing 3 stop codons after amino acid 10. Mutagenesis and conjugation was carried out as reported previously ^64,97^. For conjugation, donor strain *E. coli* JKE201 from A. Harms and C. Dehio^98^ and recipient *F. novicida* U112 strains were grown until a OD_600_ of 1 was reached. 1 ml of each culture was mixed in 20 µl of LB and spotted on a LB agar plate supplemented with 1.2 mM DAP and incubated over night at 25 °C. The mixture was plated on Mueller Hinton (BD Biosciences) agar plates supplemented with 0.1 % L-cysteine, 0.1 % D-glucose (Sigma-Aldrich), 0.1 % heat inactivated fetal calf serum (Gibco), 50 µg/ml ampicillin and 15 µg/ml kanamycin and incubated at 37 °C for 2 days. *F. novicida* U112 colonies containing the suicide vector were restreaked on BHI agar plates supplemented with 15 µg/ml kanamycin. To select against pDMK3, individual colonies were restreaked on LB agar plates supplemented with 5 % sucrose (Sigma-Aldrich), 0.1 % L-cysteine and 50 µg/ml ampicillin and incubated at 25 °C until colonies lost the suicide vector. Each mutant was verified by colony PCR using primers binding outside of the recombined region and subsequent sequencing. In addition, they were restreaked on kanamycin again to check for loss of pDMK3.

To generate the *ygfB::ampR* mutant in *E. coli* K-12 *substr*. BW25113, λ Red recombineering was used^99^. The parental strain carrying the λ Red genes on plasmid was grown until an OD_600_ of 0.3 before expression of the λ Red genes was induced with 0.2 % L-arabinose (Sigma - ldrich). At OD_600_ of 0.8, cells were made electrocompetent^100^. First, the cells were cooled on ice for 30 min before washed twice with cold 10 % glycerol (Sigma-Aldrich). The washed cells were resuspended in cold GYT buffer (10 % glycerol, 0.125 % yeast extract (Thermo Fisher Scientific) and 0.25 % tryptone (Difco)) and electroporated with 1 µg of a dialyzed PCR product encoding an *ampR* gene including promoter with 50 bp long homology arms matching the DNA sequence of *ygfB* at amino acid 30 (2 mm Bio-Rad Gene Pulser cuvette at 2.5 kV). Cells were resuspended in SOC medium (Invitrogen) and incubated at 37 °C shaking for recovery for 2 h. Dilutions of cells were plated on LB agar plates containing ampicillin to select for the *ygfB::ampR* mutant. The next day, mutants were confirmed by PCR. Loss of the recombineering plasmid in the *ygfB::ampR* was facilitated by inducing expression of λ Red gene with 0.2 % L-arabinose overnight in the *ygfB::ampR* without selection for plasmid retention. Plasmid loss was confirmed by replica plating on LB agarose plates either containing ampicillin or chloramphenicol. The final mutant strain was confirmed by Sanger sequencing of the *ygfB::ampR* region.

To generate the pFNMB2 *multiple cloning site* plasmid (empty vector control), pFNMB2 *msfGFP*^97^ was cut with Mlu1 (NEB) and Xma1 (NEB) restriction enzymes and the amplified multiple cloning site was cloned into pFNMB2. The two protospacers 3 and 5 of the native type V-A array were cloned into pFNMB *msfGFP* in two steps, by using primers with overhangs containing first the sequence for protospacer 3 and then subsequently the sequence for protospacer 5. For supplementation of a wild-type copy of *mtvS1*, *mtvS2*, *E. coli* K-12 *substr*. BW25113 *ygfB* and the *cas12* promoter reporter, pFNMB2 *msfGFP* was cut with Spe1 (NEB) and Xma1 restriction enzymes and amplified genes including their endogenous promoters or *cas12* promoter linked to *mCherry2* were cloned into the backbone. Plasmids were moved into *F. novicida* U112 by conjugation or into *E. coli* K-12 *substr*. BW25113 by electroporation.

All primers and plasmids are listed in Supplementary Table 3.

### Fluorescence live cell imaging

A Nikon Ti-E inverted motorized microscope equipped with Perfect Focus System and a CFI Plan Apochromat DM Lambda 100X Oil objective lens (NA 1.4) was used for live-cell imaging. A SOLA Light Engine (Lumencor) set to 5 % power and C-FL GFP HC HISN Zero Shift filter (excitation: 470/40 nm (450–490 nm), emission: 525/50 nm (500–550 nm), dichroic mirror: 495 nm) and C-FL DS RED HC HISN Zero Shift filters (excitation: 545/30nm (530-560nm), emission: 620/60nm (590-630nm), dichroic mirror: 570nm), respectively, were used to excite and filter fluorescence from msfGFP and mCherry2. The exposure time for each channel was set to 500 ms and images were collected with a Zyla 4.2 sCMOS (Andor, pixel size 65 nm) or an ORCA-FusionBT sCMOS camera (Hamamatsu, pixel size 65 nm). Phase contrast was imaged with the SOLA light Engine set to 2% power and 10 ms excitation. The microscope was controlled by the software NIS-Elements AR version 5.21.03.

Exponentially grown (rod-shaped) and stationary phase (roundish) *F. novicida* U112 were spun down and resuspended in phosphate saline buffer (PBS, Gibco) and spotted on pads consisting of 1% agarose in PBS. Snapshots of phase contrast, RFP and GFP channels were taken immediately after.

Long-term imaging experiments which include switching of media, were carried out in CellASIC ONIX microfluidic plates (Millipore). Exponentially growing bacteria were resuspended in BHI supplemented with 0.2 % L-cysteine and loaded into the microfluidic plate along with fresh BHI supplemented with 0.2 % L-cysteine for initial growth and the tested medium as specified in the figure legends. For generating spent overnight medium, cells in stationary phase were pelleted for 1.5 minutes at 16000 g and the supernatant was filtered with 0.22 µm filters (Millipore). For heat-treated spent overnight medium, filtered spent overnight medium was incubated at 95 °C for 10-20 minutes. After connecting the plate to the CellASIC ONIX2 Microfluidic System, the plates were incubated at 37 °C for 1 h using a Tokai HIT thermal box (Zeiss). Images of phase contrast, RFP and GFP channel were taken every 5 minutes after loading cells into the imaging chamber. The medium and the flow rate was controlled by the software ONIX2 version 1.0.4. First, fresh BHI was flown over for 1 h before switching to the tested medium for 4h. Afterwards, fresh BHI was flown over again for 3 h. For all media, the flow rate was set to 13.8 kPa.

### Image analysis

Image analysis was carried out with Fiji software version 1.54p^101^. For comparison of fluorescent signal intensities, the contrast was set to same values for sets of compared images in same sub panel. If compared across sub panels, it is stated in the figure legends.

Kymographs were created with the reslice function.

For long-term imaging, images were aligned to the first image using NIS-Elements AR version 5.21.03.

### RNA-seq

For *F. novicda* U112, RNA was extracted from exponentially growing and stationary phase cells in three biological replicates for each condition. Cells yielding in a total OD_600_ of 3 were pelleted and resuspended in 80 µl of PBS and 20 µl of 10 % Sarcosyl (Sigma-Aldrich). Then, RNA was extracted with TRI Reagent according to the manufacturer’s protocol (Zymo Research). An in-column DNA digest during RNA extraction was performed. A second DNA digestion was carried out with 10 µg of RNA using Turbo DNase (Invitrogen) for 2x 30 minutes. The DNA-free RNA was cleaned up with the RNA Clean & Concentrator-5 kit (Zymo Research) and eluted in 15 µl of water. RNA quality was checked by running a bleach agarose gel^102^. RNA concentrations were measured with a Qubit 4 fluorometer (Invitrogen) using the Qubit RNA Broad Range Assay Kit.

Ribosomal RNA was depleted with the NEBNext rRNA depletion (bacteria) kit (NEB). An input of 1 µg of RNA was used. Then, the RNA was prepared for sequencing using the TruSeq Stranded mRNA kit (Illumina) according to manufacturer’s instruction. For the first strand synthesis, Superscript IV reverse transcriptase (Invitrogen) was used with an annealing temperature of 50 °C. Final DNA concentrations and lengths were measured with a Qubit 4 fluorometer (Invitrogen) using the Qubit dsDNA High Sensitivity Assay Kit and the Tapestation 4200 bioanalyzer (Agilent) with the Agilent D500 ScreenTape Assay (Agilent).

The DNA libraries for the *mtvS1 nonsense* mutant were diluted to 12 pM, mixed with 2.5% 20 pM PhiX and subjected to single read sequencing with a 150-cycle MiSeq Reagent Kit v3 (Illumina) on the Illumina MiSeq platform (firmware v4.0). DNA libraries for the *mtvS2 nonsense* mutant were diluted to 0.8 nM and mixed with 2% 0.6 nM PhiX. 100 pM of these libraries were subjected to single read sequencing with a 25 M 300 cycles kit on a MiSeq i100 platform(firmware v1.0.109).

The reads were mapped to the *F. novicida* U112 genome (RefSeq NC_008601^103^) and RPKM values for each gene was calculated using coordinates of coding genes (ptt files) and of rRNAs and tRNAs (rtt files) with EDGE-Pro v1.3.1^104^. Significant changes in gene expression between used conditions were calculated with R (version 4.3.0) and DESeq2 (version 1.40.2)^105^. rRNAs (*FTN_0472, FTN_0475, FTN_0476, FTN_0501, FTN_1304, FTN_1305, FTN_1308, FTN_1720, FTN_1721, FTN_1724*) are marked in italic in tables and excluded in graphs and analysis.

For *E.coli*, pellets of stationary phase and exponentially grown cells at total OD_600_ of 3 were submitted to SeqCenter in Pittsburgh, PA, USA, for RNA-Seq using the Stranded Total RNA Prep Ligation (Illumina) with the Ribo-Zero Plus kit (Illumina). Sequencing was done on a NovaSeq X Plus, producing paired end 150bp reads.

The mapping of reads to the *E. coli* K-12 subsp. MG1655 genome (RefSeq NC_000913.3^106^) was performed with EDGE-Pro v1.3.1 and gene expression changes between conditions were calculated with R (version 4.3.0) and DESeq2 (version 1.40.2). rRNAs (*b0201*, *b0204*, *b0205*, *b2588*, *b2589*, *b2591*, *b3272*, *b3274*, *b3275*, *b3278*, *b3756*, *b3758*, *b3759*, *b3851*, *b3854*, *b3855*, *b3968*, *b3970*, *b3971*, *b4007*, *b4009*, *b4010*) are marked in italic in the tables and excluded in graphs and analysis.

EDGE-Pro statistics for each sample can be found in Supplementary Table 4-6. Results and normalized counts are listed in the Supplementary Data 1, tab C-H and P-R). *p*-values of 0.05 (Wald test) or adjusted *p*-value (adj*p*, Benjamini-Hochberg) of 0.05 and log2 foldchanges of 2 were defined as significant.

### RT-qPCR

RNA was extracted as reported for RNA-seq. The RNA was reverse transcribed to cDNA with Superscript IV reverse transcriptase (Invitrogen) using 1 µg of RNA, random hexamer primers, and a reaction temperature of 50 °C.

qPCR was performed in the QuantStudio 3 Real-Time PCR System (Applied Biosystems) using the PowerUp SYBR Green Master Mix (Applied Biosystems). Reactions were set up in a total volume of 20 µl containing 50 ng (*F. novicida* U112) or 250 ng (*E. coli* K-12 *substr*. BW25113) of cDNA template and 500 nM primers. Each reaction was set up in three technical replicates. All primers used for RT-qPCR are found in Supplementary Table 7. The annealing and amplification temperature was set to 60 °C.

The geometrical mean generated with Diomni™Design and Analysis (RUO) version 3.1.0 of the three technical replicates for each biological replicate was used for further analysis. RNA levels of probed genes were normalized to the DNA 3’-5’ helicase gene *uvrD* (*FTN_1594*) for *F. novicida* U112 and DNA gyrase subunit B gene *gyrB* (*b3699*) *for E. coli* K-12 *substr*. BW25113, which both have been used previously for normalization and do not change its expression in the used conditions as determined by RNA-seq^107,108^. Relative RNA abundance of probed genes was calculated as 2^-ΔCt^.

Statistical analysis for RT-qPCR data was performed with Prism10.2.3 (403) (GraphPad Software). To test if relative RNA abundances determined by RT-qPCR are significantly different, two-way ANOVA with correction for multiple comparison (Tukey’s multiple comparisons test with 95 % confidence intervals) or a Welch’s *t*-test was performed*. p*-values and adjusted *p*-values (adj*p*) are given in the figures.

### 5’ RACE

Rapid amplification of 5’ cDNA ends (5’ RACE) was carried out with RNA extracted as reported for RNA-seq. Parental strain RNA from both growth phases in three biological replicates was used. Sequences of template switch oligo (TSO) and primers are listed in Supplementary Table 8.

The manufacturer’s protocol was followed. In short, 1 µg of RNA was annealed with the RT primer for the gene of interest (10 µM) and 10 mM dNTPs (Thermo Fisher Scientific) in a 6 µl reaction at 70 °C for 5 minutes. Afterwards, the reaction was put on ice. Then the Template Switching RT Enzyme Mix (NEB) and the template switch oligo (3.75 µM) were added (final volume: 10 µl) and reverse transcription and template switching was performed in a thermocycler (BioRad) for 90 minutes at 42 °C, followed by 5 minutes at 85 °C and hold at 4°C. The reaction was diluted 1:2 with H2O. The 5’ region of transcripts was amplified with a touch down PCR using Q5 Hot Start High-Fidelity polymerase (NEB), 4 µl of the diluted template and primers (10 µM) binding to the template switch oligo and to the gene of interest. The PCR products were sent for Sanger sequencing and align to the *F. novicida* U112 genome (RefSeq NC_008601^103^) with SnapGene v8.2.2.

### Whole cell lysate proteomics

Three biological replicates of exponentially growing and stationary phase parental strain and *mtvS1 nonsense* mutant cells yielding a total OD_600_ of 5 were washed once with PBS and resuspended in 500 µl lysis byffer (20 mM Tris-HCl pH 7.5 (VWR, Thermo Fisher Scientific), 100 ml KCl, 2 % NP-40 (Thermo Fisher Scientific), mini cOmplete EDTA-free protease inhibitor cocktail (Roche)). Samples were lysed by 4 rounds of sonication on ice (40% amplitude, 5 s pulses, 30 s breaks, Q500 sonicator). Cell debris was removed by centrifugation at 21000 g, 4 °C for 10 minutes. Protein concentrations were measured with a BCA protein assay (Pierce). Samples were stored at -80 °C until 50 µg of protein per sample was subjected to LC-MS/MS analysis.

### Promoter sequence pull-down

Pull-down of proteins binding the *cas12* promoter and the intragenic *cas12* control sequence was carried out in three biological replicates as reported previously^109,110^.

In short, primers with a 5’ Biotin-TEG label were used to generate bait DNA sequences from *F. novicida* U112 gDNA. Primers are listed in Supplementary Table 9. PCR products were checked by agarose gel and cleaned up with QIAquick PCR Purification Kit (QIAGEN) and concentrated with Amicon Ultra-0.5 mL Centrifugal Filters (Millipore Sigma) to 200 ng/µl in H_2_O. Washed Dynabeads MyOne Streptavidin C1 (Invitrogen) were loaded with 40 µg of DNA for 20 minutes. DNA concentrations in the removed supernatants were measured to ensure that most DNA bound to the Dynabeads. Exponentially growing and stationary phase parental strain cells yielding a total OD_600_ of 800 were resuspended in 5 ml 1x FastBreak cell lysis buffer (Promega) in BS/THES buffer (10 mM HEPES (Thermo Fisher Scientific) pH 7.5, 5 mM CaCl_2_ (Sigma_Aldrich), 50 mM KCl (Sigma-Aldrich), 12 % glycerol, 22 mM Tris-HCl pH 7.5, 4 mM EDTA (Thermo Fisher Scientific), 9 % sucrose (Sigma-Aldrich), 62 mM NaCl (Sigma-Aldrich), mini cOmplete EDTA-free protease inhibitor cocktail) for 30 minutes on ice and then sonicated afterwards. 200 µl of precleared lysates was added to the resuspended beads supplemented with 0.03 µg/µl salmon sperm DNA (Thermo Fisher Scientific) and incubated for 1 h with agitation at 4 °C. The beads were washed 5x with BS/THES buffer and 2x with wash buffer (100 mM NaCl, 25 mM Tris-HCl pH 7.5) and then switched to new tubes before a last wash. The dried beads were stored at 4 °C until they were subjected to LC-MS/MS analysis.

### MtvS1-His pull-down

Pull-down of proteins binding MtvS1-His_6_ was carried out by following the same protocol as reported previously and used for the promoter sequence pull-down with changes indicated below^109,110^.

The *mtvS1 ns* mutant was grown to exponential and stationary phase in three biological replicates yielding a total OD_600_ of 450 each and resuspended in 5 ml 1x FastBreak cell lysis buffer in BS/THES supplemented with 0.5 mg/ml lysozyme (Sigma-Aldrich) and 0.1 mg/ml DNase I (Roche). Lysis buffer did not contain EDTA as it is not compatible with the Dynabeads His-Tag Isolation and Pull-down beads (Invitrogen) according to the manual. For cell lysis, the samples were incubated shaking at 4 °C for 1 h and subsequently sonicated according to promoter sequence pull-down protocol. Cell debris was removed by centrifugation and filtration of lysates through 0.45 µm filters (Cytiva). 50 µl of prewashed Dynabeads His-Tag Isolation and Pull-down beads per sample were loaded with 80 µg purified MtvS1-His_6_ according to instructions and incubated on a roller at 4 °C for 30 minutes. Empty beads subjected to the same treatment as beads loaded with MtvS1-His_6_ served as controls for non-specific binders. After three washes, 350 µl lysates and 350 µl BS/THES buffer were added to the beads and incubated on a roller at 4 °C for 1 h. Beads were washed similarly to promoter pull-down protocol and the dried beads were stored at -20 °C until subjected to LC-MS/MS analysis.

### LC-MS/MS analysis

For whole cell lysates, proteins were reduced and alkylated followed by acetone precipitation. Precipitates were dissolved in 50 µl 1 µg of trypsin (Promega)/60 mM ammonium bicarbonate and digested overnight. An additional 1 µg of trypsin was added following heating to 45 °C. Digestions were halted by adding 10% trifluoroacetic acid, volumes were spun (16,000 g) and two thirds of the samples were subjected to solid phase extraction (Empore). Data was acquired on an Ascend orbitrap mass spectrometer using either DDA or DIA, and peptides were separated using a 25 cm EasySprayer column. A 120 minutes and 90 minutes analytical gradient increasing from 1% B to 30% B (B: 80% ACN,0.1% formic acid) was used.

For the promoter sequence and MtvS1-His_6_ pull-downs, beads were resuspended in 8 M urea and proteins were reduced and alkylated on beads. The supernatants were extracted, and proteins were digested with LysC (Wako/Fuji) with urea concentration lowered to less than 4 M, followed by trypsination (Promega) with urea concentration lowered to less than 2 M. Digestion was stopped with 10 % trifluoroacetic acid. Generated peptides were desalted and concentrated with solid-phase extraction prior to analysis by LC-MS/MS (promoter sequence pull-down: 12 cm packed-in-emitter column, 70 minutes analytical gradient, high resolution/high mass accuracy mode (Dionex 3000 and Q-Exactive HF, Thermo Fisher Scientific), MtvS1-His_6_ pull-down: 25 cm EASYSprayer column, 70 minutes analytical gradient, high resolution/ high mass accuracy mode (Ascend)).

For analysis, generated data was searched and quantified using ProteomeDiscoverer (v 1.4.1.14 and 3.2)/Mascot (v 2.8 and 3.1), MaxQuant (v 2.0.3.0 and 2.6.6.0) and Spectronault (v 19.7) and queried against Uniprot database for *F. tularensis* subspecies *novicida* U112 (taxon ID: 401614) concatenated with common contaminants to generate iBAQ values. Statistical analysis was carried out with Perseus software (v 1.6.15.0). The values were log2 transformed and contaminants and proteins that have been identified in less than two biological replicates of a sample were removed. After filtering, the data was median normalized and normalized iBAQ values were produced.

For whole lysate proteomes, the values were log2 transformed and contaminants and proteins that have not been detected in all three replicates for a least one condition were removed. Missing values were imputed. Log2 foldchanges were calculated for each strain and condition. An FDR-based (FDR = 0.05) two-sample *t*-test was used to determine which changes were significant. Results are listed in Supplementary Data 1, tab A. *p*-values of 0.05 or adj*p*-values of 0.05 and log2 foldchanges of 1 were defined as significant.

For the pull-downs, the values were log2 transformed and contaminants and proteins that have been identified in less than two biological replicates of a sample were removed. After filtering, the data was median normalized and normalized iBAQ values were produced. The log2 foldchange was calculated between proteins pulled-down with the *cas12* promoter sequence and proteins pulled-down with the intragenic *cas12* control sequence for both exponential and stationary conditions. An FDR-based (FDR = 0.05) two-sample *t*-test was used to determine which changes were significant. Results are listed in Supplementary Data 1, tabs I-K. *p*-values of 0.05 or adj*p*-values of 0.05 and log2 foldchanges of 2 were defined as significant.

### Plasmid interference assay

Strains were generated with the native type II-B or type V-A CRISPR array replaced by a targeting or non-targeting spacer matching expression plasmid pFNMB2 *msfGFP*. These strains were grown exponentially together with the donor *E. coli* JKE201 strain harboring pFNMB2 *msfGFP*. To assess the Cas12-dependent targeting capability of the *mtvS1-2 nonsense* mutant, parental and mutant strains containing native type V-A CRISPR arrays and two additional deletions of restriction endonucleases (Δ*FTN_0710* and Δ*FTN_1487*) as well as a *E. coli* JKE201 donor strain carrying pFNMB2 *sp3_typeV_seedG3T_targ sp5_typeV_seedA3C_targ* were used. The two protospacers 3 and 5 encoded on the plasmid contained seed mutations at position 3 to make targeting less efficient. Strains and plasmids used are listed in Table 2 and 3.

Cells yielding a total OD_600_ of 1 were pelleted and washed once with LB. Then, recipient *F. novicida* U112 strains were mixed with donor *E. coli* JKE201 cells in a final volume of 20 µl LB and spotted on nitrocellulose filter membranes with a 0.45 µm pore size (Millipore) located on LB agar plates supplemented with 1.2 mM DAP. The plates were incubated for 2 h at 37 °C before the bacteria were resuspended in a final volume of 100 µl MH broth (BD Biosciences). Serial dilutions in MH broth were carried out and 5 µl of each dilution was spotted in technical duplicates on MH agar plates supplemented with 50 µg/ml ampicillin for calculation of total number of colony forming units (CFUs) and on MH agar plates supplemented with 50 µg/ml ampicillin and 15 µg/ml kanamycin for calculation of number of transconjugants. The average number of CFUs of the two technical replicates was used for calculating CFU/ml. Three biological replicates for each strain were analyzed.

To calculate the conjugation efficiency, the calculated concentration of transconjugants was divided by the concentration of recipient cells. Significant differences were determined with a one-way ANOVA and Tukey’s multiple comparison test (95 % confidence interval).

### Growth curves

Strains were grown overnight in liquid cultures, diluted 1:20 the next day and grown until OD_600_ of ∼1. For OD_600_ measurements with a plate reader (Agilent Synergy H1, Gen5 version 3.15.15 software), OD_600_ was adjusted to 0.2 and cells were seeded in 150 μl per well in a 96-well plate (Greiner Cellstar, flat bottom with lid) in technical triplicates. Absorbance at 600 nm was measured every 10 minutes for 24 h while plate was incubated at 37 °C and shaking with 567 cpm (3 mm). Growth curves for three biological replicates per strain were measured. Means with standard deviation are shown.

### SDS-PAGE

For protein analysis by SDS-PAGE, 4-20 % polyacrylamide gels (BioRad) were used and proteins were resuspended in 1x Laemmli buffer (BioRad). 15 µl of sample was loaded together with 5 µl of Precision Plus Protein™ WesternC standard (BioRad) and gel was run at 140 V for 50 min. Staining was performed with Coomassie Brilliant Blue R-250 (Biorad).

### Western blot

*F. novicida* U112 strains encoding *mtvS1-His_6_* on the chromosome was were grown exponentially and to stationary phase yielding a total OD_600_ of 200 and resuspended in 30 ml lysis buffer (50 mM HEPES pH 7.5, 250 mM NaCl, 10 % glycerol, 0.5 mg/ml lysozyme, 1 mM Tris (2-Carboxyethyl) phosphine hydrochloride (TCEP, GoldBio), mini cOmplete EDTA-free protease inhibitor cocktail). The cells were lysed for 1 h at 4 °C shaking and sonicated for 15 rounds on ice (70 % amplitude, 10 s pulses, 30 s breaks). 15 µl of each sample and 5 µl of 5 µl of Precision Plus Protein™ WesternC standard were run on a SDS-PAGE gel and then transferred to nitrocellulose membrane (0.2 µm, BioRad) using a Trans-Blot Turbo transfer system (BioRad). After blocking the membrane in 5 % milk in TBST (PBS, 0.05 % Tween 20 (Sigma-Aldrich)) for 1 h, MtvS1-His_6_ was probed with primary mouse THE His Tag anti-His_6_ antibody (GenScript, A00186) diluted 1:4000 in PBST at 4 °C overnight. The membrane was incubated with the secondary goat anti-mouse IgG (H+L) HRP antibody (Thermo Fisher Scientific, 31430) diluted 1:10000 in PBST for 1 h. HRP signal was detected with Clarity Western ECL substrate (BioRad) and ImageQuant 800 (Amersham) in SNOW imaging mode.

### Expression and purification of *E. coli* RNA polymerase

Plasmid pVS11 (also called pEcrpoABC(-XH)Z) was used to overexpress each subunit of RNA polymerase (full-length α, β, ω) as well as β’-PPX-His10 (PPX; PreScission protease site, LEVLFQGP, GE Healthcare), and co-transformed with a pACYCDuet-1 plasmid containing *E. col rpoZ* into *E. coli* BL21(DE3) to ensure saturation of all RNA polymerases with *E. coli* RpoZ. The cells were grown in the presence of 100µg/ml ampicillin and 34μg/ml chloramphenicol to an OD600 of 0.6 in a 37 °C shaker. Protein expression was induced with 1 mM isopropyl β-d-1-thiogalactopyranoside (IPTG, final concentration) for 4 h at 30 °C. Cells were harvested by centrifugation and resuspended in 50mM Tris-HCl pH 8 (VWR, Thermo Fisher Scientific), 5% w/v glycerol, 10 mM Dithiothreitol (DTT), 1 mM PMSF, and 1x protease inhibitor cocktail (PIC - 0.174 mg/ml PMSF, 312 μg/ml benzamidine, 5 μg/ml chymostatin, 5 μg/ml leupeptin, 1 μg/ml pepstatin A, 10 μg/ml aprotinin dissolved in ethanol). After French Press (Avestin) lysis at 4 °C, the lysate was centrifuged twice for 30 min each. Polyethyleneimine (PEI, 10% (w/v) pH 8, Acros Organics – ThermoFisher Scientific) was slowly added to the supernatant to a final concentration of ∼0.6 % PEI with continuous stirring. The mixture was stirred at 4 °C for an additional 25 min, then centrifuged for 1.5 h at 4 °C. The pellets were washed three times with 50 mM Tris-HCl pH 8, 500 mM NaCl, 10 mM DTT, 5 % w/v glycerol, 1 mM PMSF, 1x PIC. For each wash, the pellets were homogenized, then centrifuged again. RNA polymerase was eluted by washing the pellets three times with 50 mM Tris-HCl pH 8, 1 M NaCl, 10 mM DTT, 5% w/v glycerol, 1x PIC, 1 mM PMSF. The PEI elutions were combined and precipitated with ammonium sulfate, at a final [(NH_4_)_2_SO_4_] of 35% w/v, overnight. The mixture was centrifuged, and the pellets were resuspended in 20 mM Tris-HCl pH 8, 1 M NaCl, 5% w/v glycerol, 1 mM 2-mercaptoethanol (BME). The mixture was loaded onto two 5 ml HiTrap IMAC HP columns (Cytiva) for a total column volume of 10 ml. RNA polymerase (β’-PPX-His10) was eluted at 250 mM imidazole (Sigma-Aldrich) in column buffer (20 mM Tris-HCl pH 8, 1 M NaCl, 5% w/v glycerol, 1 mM BME). The eluted RNA polymerase fractions were combined and dialyzed to a final 10 mM Tris-HCl pH 8, 0.1 mM EDTA pH 8, 100 mM NaCl, 5% w/v glycerol, 5 mM DTT. The sample was then loaded onto a 40 ml Bio-Rex-70 column (Bio-Rad), washed with 10 mM Tris-HCl pH 8, 0.1 mM EDTA, 5% w/v glycerol, 5 mM DTT in isocratic steps of increasing NaCl concentration (elutes at 0.5M NaCl). The eluted fractions were combined, concentrated by centrifugal filtration, then loaded onto a 320 ml HiLoad 26/600 Superdex 200 column (Cytiva) equilibrated in gel filtration buffer (10 mM Tris-HCl pH 8, 0.1 mM EDTA pH 8, 0.5 M NaCl, 5 % w/v glycerol, 5 mM DTT). The eluted RNA polymerase was concentrated to 8-10 mg/ml by centrifugal concentration and supplemented with glycerol to 20% w/v, flash frozen in liquid N2, and stored at −80°C.

### Expression and purification of *E. coli* σ^70^

Plasmid pSAD1403 was transformed into *E. coli* BL21(DE3). The cells were grown in the presence of 50 μg/ml kanamycin to an OD_600_ of 0.6 at 37 °C. Protein expression was induced with 1 mM IPTG for 1-1.5 h at 30 °C. Cells were harvested by centrifugation and resuspended in 20 mM Tris-HCl pH 8, 500 mM NaCl, 0.1 mM EDTA pH 8, 5 mM imidazole, 5% w/v glycerol, 0.5 mM BME, 1 mM PMSF, 1x PIC (the same as used for the *E. coli* RNA polymerase). After French Press, lysis at 4 °C, cell debris was removed by centrifugation twice. The lysate was loaded onto two 5 ml HiTrap IMAC HP columns for a total column volume of 10 ml. His_10_-SUMO-σ^70^ was eluted at 250 mM imidazole in 20 mM Tris-HCl pH 8, 500 mM NaCl, 0.1 mM EDTA pH 8, 5% w/v glycerol, 0.5 mM BME. Peak fractions were combined, cleaved with Ulp1, and dialyzed against 20 mM Tris-HCl pH 8, 500 mM NaCl, 0.1 mM EDTA pH 8., 5% w/v glycerol, 0.5 mM BME, resulting in a final imidazole concentration of 25 mM. The cleaved sample was loaded onto one 5 ml HiTrap IMAC HP to remove His10-SUMO-tag along with any remaining uncleaved σ^70^. Untagged σ^70^ fractions were pooled and diluted with 10mM Tris-HCl pH 8, 0.1 mM EDTA pH 8, 5% w/v glycerol, 1 mM DTT until the conductivity corresponds to a NaCl concentration slightly below 200mM. The diluted sample was injected on to three 5 ml HiTrap Heparin HP columns (total column volume of 15 ml; Cytiva) equilibrated at the same diluent buffer but with 200 mM NaCl, and run a gradient to 1 M NaCl, with the first major peak as the target peak. The target peak sample was pooled and concentrated by centrifugal filtration before being loaded onto a HiLoad 16/60 Superdex 200 (Cytiva) in 20 mM Tris-HCl pH 8, 500 mM NaCl, 5% w/v glycerol, 1 mM DTT. Peak fractions of σ^70^ were pooled, supplemented with glycerol to a final concentration of 20% w/v, flash-frozen in liquid N_2_, and stored at -80°C.

### Expression and purification of MtvS1-His_6_

MtvS1 (FTN_1238) was cloned into pET21 with a C-terminal His_6_-tag separated by a glycine linker. MtvS1-His_6_ overexpression was performed in *E. coli* Rosetta 2 (DE3). 2 l of bacteria were grown to OD_600_ of 0.7 at 37 °C before the temperature was shifted to 16 °C and MtvS1 expression was induced with 500 µM IPTG (GoldBio) over night. Cells were harvested in 50 ml lysis buffer (50 mM HEPES pH 7.5, 250 mM NaCl, 10 % glycerol, 0.5 mg/ml lysozyme, 10 mM imidazole, 1 mM TCEP-HCl, mini cOmplete EDTA-free protease inhibitor cocktail) and lysed for 1 h at 4 °C shaking before 15 rounds of sonication on ice (70 % amplitude, 10 s pulses, 30 s breaks). Cell lysate was centrifuged at 18000 g, 4 °C for 1 h and the supernatant was filtrated with 0.45 µm filters to remove remaining cell debris. After loading the column with Nickel resin (3 ml HisPur Ni-NTA, Thermo Fisher Scientific) and equilibrating the resin with lysis buffer, the cell lysate was loaded, followed by two washes with 50 ml wash buffer (50 mM HEPES pH 7.5, 1 M NaCl, 10 % glycerol, 10 mM imidazole, 1 mM TCEP-HCl). Then, MtvS1-His_6_ was eluted with 30 ml elution buffer (50 mM HEPES pH 7.5, 250 mM NaCl, 10 % glycerol, 300 mM imidazole, 1 mM TCEP-HCl). Elute was concentrated with 10 kDA cut-off filter (Amicon) and loaded on a Superdex 200 10/300-increase column (Cytiva) pre-equilibrated in size-exclusion buffer (50 mM HEPES pH 7.5, 150 mM NaCl, 10 % glycerol, 1 mM TCEP-HCl) for further purification by size-exclusion chromatography.

Purified MtvS1-His_6_ was stored at -80 °C.

### Expression and purification of MtvS2-His_6_

MtvS2 (FTN_1519) was cloned into pET21 with a C-terminal His_6_-tag separated by a glycine linker and three additional stop codons after the His_6_-tag. MtvS2-His_6_ overexpression was performed in *E. coli* Rosetta 2 (DE3). 6 l of bacteria were grown to OD_600_ of 0.4 at 30 °C before MtvS2 expression was induced with 500 µM IPTG for 2 h at 30 °C. Cells were harvested in 50 ml denaturing lysis buffer (50 mM HEPES pH 8, 250 mM NaCl, 10 % glycerol, 8 M urea (Fisher Scientific), 0.5 mg/ml lysozyme, 10 mM imidazole, 1 mM TCEP-HCl, mini cOmplete EDTA-free protease inhibitor cocktail) and lysed for 1 h at 4 °C shaking before 15 rounds of sonication on ice (70 % amplitude, 10 s pulses, 30 s breaks). Cell lysate was centrifuged at 18000 g, 4 °C for 1 h and the supernatant was filtrated with 0.45 µm filters to remove remaining cell debris. Purification was carried out at 4 °C. After loading the column with Nickel resin and equilibrating the resin with lysis buffer, the cell lysate was loaded, followed by two washes with 50 ml wash buffer (50 mM HEPES pH 8, 1 M NaCl, 10% glycerol, 6 M urea10 mM imidazole, 1 mM TCEP-HCl). Then MtvS2-His_6_ was eluted with 30 ml elution buffer (50 mM HEPES pH 8, 250 mM NaCl, 10 % glycerol, 6 M urea, 300 mM imidazole, 1 mM TCEP-HCl). Elute was concentrated with 10 kDA cut-off filter for dialysis. Dialysis was carried out at 4 °C with Pur-A-Lyzer Maxi 6000 tubes (Sigma-Aldrich) in 1 l dialysis buffer (50 mM HEPES, pH 8,150 mM NaCl, 30 % glycerol, 0.5 mM EDTA, 1 mM TCEP-HCl) containing first 4 M urea for 4 h, then 2 M urea overnight and last, 0 M urea for 4 h. Purified and dialyzed MtvS2-His_6_ was stored at -80 °C.

### Electrophoretic mobility shift assay

Electrophoretic mobility shift assays were carried out as described previously^111,112^. Primers for *cas12* promoter and *cas12* intragenic control sequence are listed in Supplementary Table 10. PCR products amplified from genomic DNA, cleaned up and eluted in H_2_O.

50 nM of PCR products were incubated with 0, 1, 5, 10 or 20 µM of MtvS1-His_6_ and MtvS2-His_6_ individually or combined in 20 µl EMSA buffer (20 mM Tris-HCl pH 8, 0.1 mM EDTA-Na_2_ (Sigma-Aldrich), 50 mM KCl, 5 % glycerol and 0.1 mM DTT (Goldbio)) for 30 min on ice. Samples were then loaded with SDS-free loading dye (NEB) and a 100 bp DNA ladder (NEB) on a 3 % 1x Tris-borate (TB) agarose gel (Sigma-Aldrich) and separated by electrophoresis (2 h, 80 V) at 4 °C using 1x TB buffer as running buffer. The gel was stained with 2 µg/ml ethidium bromide (Sigma-Aldrich) for 20 min and washed 3x 10 min with H_2_O. The DNA was then visualized with the ImageQuant 800 (Amersham).

### Pull-down of *E. coli* RNA polymerase

Interaction of MtvS1-His_6_ and MtvS2-His_6_ with *E. coli* RNA polymerase was probed as described previously^113^. In short, purified MtvS1-His_6_ (20 µM) was incubated with *E. coli* RNA polymerase core enzyme or holoenzyme (400 units/ml, NEB) in 40 µl reaction buffer (10 mM Tris-HCl pH 7.5, 50 mM KCl, 1 mM DTT) at 37 °C for 40 minutes. MtvS2-His_6_ (20 µM), after a buffer exchange into reaction buffer (pH8) with desalting columns (Zeba, 7K MWCO, 0.5 ml), was incubated with *E. coli* RNA polymerase core enzyme in 60 µl reaction buffer (pH 8). Afterwards for MtvS1-His_6_, 260 µl of reaction buffer H (20 mM HEPES pH 7.5, 50 mM KCl, 10 mM imidazole, 0.001 % Triton-X (Sigma-Aldrich)) supplemented with 0.1 mg/ml bovum serum albumin (BSA, Sigma-Aldrich) was added to the reactions. For MtvS2-His_6_, reactions were resuspended in 260 µl reaction buffer H (pH 8). 50 µl of Dynabeads His-Tag Isolation and Pull-down beads per sample were washed twice with 300 µl of reaction buffer H supplemented with BSA before the samples were transferred to the prewashed beads and incubated on a roller at room temperature for 5 minutes (MtvS1-His_6_) or for 30 minutes (MtvS2-His_6_). The beads were washed once with 300 µl reaction buffer H supplemented with BSA and subsequently washed three times with reaction buffer H. Afterwards, the proteins were eluted from the beads in 30 µl reaction buffer H containing 300 mM imidazole by incubating on a roller at room temperature for 5 minutes. Beads for MtvS2-His_6_ pull-down were boiled at 95 °C for 10 minutes in 1x Laemmli buffer for better elution. 15 µl of eluted samples and individual controls (MtvS1-His_6_ and MtvS2-His_6_: 20 µM, *E. coli* RNA polymerase core enzyme and holoenzyme: 400 units/ml) were analyzed on a SDS-PAGE gel with subsequent Coomassie staining.

### *In vitro* transcription assays

*In vitro* transcription assays were based on a protocol reported previously^114^. Aptamer sequences and primers are listed in Supplementary Table 11.

The spinach aptamer sequence flanked by first 32 base pairs of *cas12* and 32 base pairs of the middle of *cas12* as well as the *E. coli rrnB T1* terminator was ordered as gblock (Azenta). The gblock was cloned into pFNMB flanked upstream by the amplified promoter sequence (400 bp upstream of gene of interest). Linearized aptamers were amplified from plasmid, cleaned up and eluted in H_2_O.

The storage buffer of recombinant *E. coli* RNA polymerase and σ^70^ as well as *F. novicida* U112 MtvS1-His_6_ and MtvS2-His_6_ was exchanged to transcription buffer (50 mM Tris-HCl pH8, 150 mM KCl, 10 mM MgCl_2_, 1 mM DTT, 0.1 mg/ml BSA) with desalting columns (7K MWCO, 0.5 ml). In short, the columns were centrifuged at 1500 g and 4 °C for 1 minute to expel the storage buffer. Then, the columns were washed twice with 300 μl transcription buffer supplemented with 0.01 % Tween-20 (Millipore). The last wash was performed with 300 μl transcription buffer only. The proteins resuspended in 50-100 μl transcription buffer were added to the columns and recovered in fresh tubes.

Complexes of *E. coli* RNA polymerase (500 nM) with *E. coli* σ^70^ (2500 nM), MtvS1 (2500 nM) or MtvS2 (2500 nM) were reconstituted by mixing them in a 1:5:5 ratio in transcription buffer followed by an incubation at 37 °C for 40 minutes.

The *in vitro* transcription assays were carried out in black 384-well plates (Corning, low volume black round bottom polystyrene NBS) in technical triplicates and no NTP controls. First, 10 nM aptamer DNA, 0.4 U/μl RiboLock RNAse inhibitor (Thermo Fisher Scientific), 0.5 mM NTPs (Thermo Fisher Scientific) and 20 μM 3,5-difluoro-4-hydroxybenzylidene imidazolinone (DFHBI, Lucerna) in 15 μl transcription buffer (final volume 20 µl) was added to each wells. Then the plate was incubated at 37 °C in the dark for 10 minutes before 50 nM RNA polymerase complexes (5 μl of the reconstituted complexes) were added. The no NTP controls contained everything except for the NTPs. Measurement of fluorescence (excitation: 470 nm, emission: 528 nm) in a plate reader (Agilent Synergy H1, Gen5 version 3.15.15 software) started immediately after adding the RNA polymerase complexes. Measurements were taken every 30 seconds for 1 h.

### Homology prediction with Foldseek

Foldseek was used to find structural homologs for the predicted structure of MtvS1 (FTN_1238, Uniprot ID: A0A6I4RTR9)^115^. PyMol (version 2.5.7) was used to generate the figure^116^.

### Heatmaps

The normalized counts (RNA-seq) and normalized log2 iBAQ values (proteomics) were used to create heatmaps for genes and proteins, which were deemed significant (adj*p*-value < 0.05) with heatmap2 on usegalaxy.org^117,118^. Z-scores were computed on rows before clustering and describe how many standard deviations the actual value is from the mean value calculated across samples for each gene.

### Gene set enrichment analysis (GSEA)

Gene set enrichment analysis was carried out with the FUNAGE-Pro^119^ (v2) online tool using the genbank file ASM1464v1 for *F. nov*icida U112 and ASM584v2 for *E. coli* K-12 *substr.* MG1655. Only genes or proteins with an adj*p* value ≤ 0.05 were considered and upregulated and downregulated genes were analyzed separately. For RNA-seq data sets, ribosomal RNAs were excluded. For comparison of protein levels between the two growth phases for the parental strain, the threshold values were set to -1> log2FC > 1. For comparison of protein levels between parental strain and *mtvS1 nonsense* mutant, the threshold values were set to - 0.5 > log2FC > 0.5. Enriched pathways for each strain and condition are listed in Supplementary Data 1, tab B. For comparison of RNA levels between the two growth conditions for the *F. novicida* U112 and *E. coli* K-12 *substr*. BW25113 parental strains and between the *F. novicida* U112 parental strain and *mtvS2 nonsense* mutant, the threshold values were set to -1.5 > log2FC > 1.5. For comparison of RNA levels between the *F. novicida* U112 parental strain and the *mtvS1 nonsense* mutant and between the *E. coli* K-12 *substr*. BW25113 parental strain and the *ygfB::ampR* mutant, the threshold values were set to -1 > log2FC > 1. Enriched pathways for each strain and condition are listed in Supplementary Data 1, tab E, H and R.

### Sequence alignments

Promoter sequences of operons significantly down regulated in stationary phase for both the *mtvS1 nonsense* and *mtvS2 nonsense mutants* according to FUNAGE-Pro were aligned with MEGA12 (version 12.0.10) using the ClustalW algorithm^120^. The sequence logo plot of the putative -35 consensus sequence was created using WebLogo^121^.

Protein sequences of MtvS1 (A0A6I4RTR9) and *E. coli* K12 YgfB (P0A8C4) were aligned with MEGA12 using the ClustalW alogrithm.

### Structural predictions with AlphaFold 3

Interactions between MtvS1 (AlphaFold ID: A0A6I4RTR9), MtvS2 (AlphaFold ID: A0Q819) and *F. novicida* U112 RNA polymerase (AlphaFold IDs: RpoA1 - A0Q4K8, RpoA2 - A0Q7R6, RpoB - A0Q867, RpoC - A0Q866, RpoZ - A0Q5J3) were predicted with the AlphaFold 3 web server (https://alphafoldserver.com, 27.02.2025)^66^.

Interactions between the coiled-coil domain of β’ subunit (RpoC) and σ^70^ or MtvS1 were compared using AlphaFold 3 predictions of *E. coli* K12 RNA polymerase (AlphaFold ID: RpoA - P0A7Z4, RpoB - P0A8V2, RpoC - P0A8T7, RpoZ - P0A800) with *E. coli* K12 σ^70^ (AlphaFold ID: RpoD - P00579) or MtvS1 (AlphaFold ID: A0A6I4RTR9) (https://alphafoldserver.com, 04.05.2025). The magnesium ion was superimposed from the solved *E. coli* RNA polymerase structure (6omf)^122^.

For assessment of prediction accuracy, the PAE values, ipTM and pTM scores were taken into account according to the guidelines.

PyMol (version 2.5.7)^116^ was used to generate figures and to determine interactions under 3 Å between MtvS1 and RpoC. These interactions are listed in Supplementary Table 1. RSMD value between MtvS1(AlphaFold ID: A0A6I4RTR9) and *E. coli* K12 YgfB (AlphaFold ID : A8A450) was also calculated with PyMol.

### Bioinformatic analysis of MtvS1 homologs

To determine the prevalence of MtvS1 across bacterial genomes and their co-occurrence with CRISPR systems, 99,999 bacterial genomes were collected from NCBI (Supplementary Data 1, tab M). All translations were predicted in these genomes using Prodigal version 2.6.3 with default settings^123^. An HMM model was created with the *F. novicida* U122 MtvS1 (FTN_1238, A0A6I4RTR9) sequence using hmmbuild with default settings (Supplementary Data 1, tab N) and then all genome translations files were scanned for MtvS1 homologs using hmmscan with default settings and a ‘--domtblout’ output all within HMMER version 3.4^124^. MtvS1 hmmscan hits with *e*-values were less than 0.01 and whose ‘hmm coord’ range was greater than 130 residues (query MtvS1 sequence is 178 residues) were classified as true hits (Supplementary Data 1, tab O). To determine the CRISPR systems that co-occur with MtvS1 homologs, we predicted the bacterial immune systems over all the bacterial genomes analyzed with DefenseFinder version 1.3.0^125^.

## Data availability

Source data are provided as a Source Data file. RNA-seq data generated in this study have been deposited at NCBI under accession code BioProject ID PRJNA1308637 (https://www.ncbi.nlm.nih.gov/bioproject/PRJNA1308637/). Proteomics data generated in this study have been deposited to the ProteomeXchange Consortium via PRIDE^126^ under accession code PXD068088 (https://www.ebi.ac.uk/pride/archive/projects/PXD068088).

## Acknowledgments

We thank D.M. Monack (Stanford University) for the conjugation plasmid pDMK3, A. Harms (ETH Zurich) and C. Dehio (Biozentrum, University of Basel) for the *E. coli* JK201 conjugation strain, M. Basler (Biozentrum, University of Basel) for plasmid pFNMB2 *msfGFP*, H. Molina and C. Peralta (Proteomics Resource Center, The Rockefeller University) for performing and analyzing the LC-MS/MS experiment. Proteomics data was generated by the Proteomics Resource Center at The Rockefeller University (RRID:SCR_017797) using instrumentation funded by the Sohn Conferences Foundation and the Leona M. and Harry B. Helmsley Charitable Trust. We thank M. Lilic from the Campbell lab (The Rockefeller University) for advice on the *in vitro* transcription assays. Thanks also go all members of the Marraffini lab (The Rockefeller University) for scientific discussions and feedback. M.B. was supported by the SNSF Early Postdoc.Mobility (P2BSP3_195673) and SNSF Postdoc.Mobility (P500PB_210998) fellowships. E.A.C. is supported by the Stavros Niarchos Foundation (SNF) as part of its grant to the SNF Institute for Global Infectious Disease Research at The Rockefeller University and NIH R35 GM151879. L.A.M. is an investigator of the Howard Hughes Medical Institute.

## Author Contributions Statement

M.B. and L.A.M. conceived the study. C.F.B. carried out the MtvS1-His_6_ purification and performed the bioinformatics analysis of MtvS1 homologs. J.C. expressed and purified the *E. coli* RNAP core enzyme and σ^70^ factor. E.A.C. analyzed data and contributed to figures. M.B. performed all the other experiments. M.B. and L.A.M. wrote the manuscript with the help of E.A.C. All authors read and approved the manuscript.

