## Supplementary Data 3 for "MtvS1 and MtvS2 Interact with RNA Polymerase to Regulate the *Francisella* Type V-A CRISPR-Cas System"

**Supplementary Movie Legends:**

**Supplementary Movie 1: Cas12-mCherry2 is expressed at high cell density.**

Fluorescence microscopy of Cas12-mCherry2 in the parental strain in a microfluidic plate. BHI was flown through for 8 h. Three representative examples are shown (n = 3 biological replicates). Pictures were taken every 5 min. Same contrast settings as in **Fig. 4a-b** and **Supplementary Movies 2-4**. Merge of phase contrast and RFP channel is shown. Fields of view are 13 x 13 µm. Scalebars represent 1 µm. Movies play at a frame rate of 7 frames per second.

**Supplementary Movie 2: Cas12-mCherry2 is not expressed at low cell density.**

Fluorescence microscopy of Cas12-mCherry2 in the parental strain in a microfluidic plate. BHI was flown through for 8 h. Three representative examples are shown (n = 3 biological replicates). Pictures were taken every 5 min. Same contrast settings as in **Fig. 4a-b** and **Supplementary Movies 1, 3-4**. Merge of phase contrast and RFP channel is shown. Fields of view are 13 x 13 µm. Scalebars represent 1 µm. Movies play at a frame rate of 7 frames per second.

**Supplementary Movie 3: Cas12-mCherry2 is not expressed at high cell density in *mtvS1 nonsense* mutant.**

Fluorescence microscopy of Cas12-mCherry2 in the *mtvS1 nonsense* mutant in a microfluidic plate. BHI was flown through for 8 h. Three representative examples are shown (n = 3 biological replicates). Pictures were taken every 5 min. Same contrast settings as in **Fig. 4a-b** and **Supplementary Movies 1-2, 4**. Merge of phase contrast and RFP channel is shown. Fields of view are 13 x 13 µm. Scalebars represent 1 µm. Movies play at a frame rate of 7 frames per second.

**Supplementary Movie 4: Cas12-mCherry2 is not expressed at low cell density in *mtvS1 nonsense* mutant.**

Fluorescence microscopy of Cas12-mCherry2 in the *mtvS1 nonsense* mutant in a microfluidic plate. BHI was flown through for 8 h. Three representative examples are shown (n = 3 biological replicates). Pictures were taken every 5 min. Same contrast settings as in **Fig. 4a-b** and

**Supplementary Movies 1-3**. Merge of phase contrast and RFP channel is shown. Fields of view are 13 x 13 µm. Scalebars represent 1 µm. Movies play at a frame rate of 7 frames per second.

**Supplementary Movie 5: *F. novicida* U112 reacts to spent overnight medium with growth arrest, morphology change and Cas12-mCherry2 expression.**

Fluorescence microscopy of Cas12-mCherry2 in the parental strain in a microfluidic plate. BHI was flown through for 1 h before switching to spent overnight medium of parental strain for 4h. Afterwards, fresh BHI was flown through for 3 h. Three representative examples are shown (n = 3 biological replicates). Pictures were taken every 5 min. Same contrast settings as in **Figure 4c-g**, **Supplementary Fig. 4d** and **Supplementary Movies 7-8**. Merge of phase contrast and RFP channel is shown. Fields of view are 13 x 13 µm. Scalebars represent 1 µm. Movies play at a frame rate of 7 frames per second.

**Supplementary Movie 6: *F. novicida* U112 reacts to spent overnight medium with growth arrest, morphology change and mCherry2 expression from *cas12* promoter.**

Fluorescence microscopy of mCherry2 in a strain carrying pFNMB *prom-cas12 mCherry2* in a microfluidic plate. BHI was flown through for 1 h before switching to spent overnight medium of parental strain for 4h. Afterwards, fresh BHI was flown through for 3 h. Three representative examples are shown (n = 2 biological replicates). Pictures were taken every 5 min. Same contrast settings as in **Supplementary Fig. 4a-c**. Merge of phase contrast and RFP channel is shown. Fields of view are 13 x 13 µm. Scalebars represent 1 µm. Movies play at a frame rate of 7 frames per second.

**Supplementary Movie 7: Cas12 is not expressed in *mtsvS1 nonsense* mutant in spent overnight medium.**

Fluorescence microscopy of Cas12-mCherry2 in the *mtvS1 nonsense* mutant in a microfluidic plate. BHI was flown through for 1 h before switching to spent overnight medium of parental strain for 4h. Afterwards, fresh BHI was flown through for 3 h. Three representative examples are shown (n = 2 biological replicates). Pictures were taken every 5 min. Same contrast settings as in **Fig. 4c-g**, **Supplementary Fig. 4d and Supplementary Movies 5 and 8**. Merge of phase contrast and RFP channel is shown. Fields of view are 13 x 13 µm. Scalebars represent 1 µm. Movies play at a frame rate of 7 frames per second.

**Supplementary Movie 8: *F. novicida* U112 does not react to heat-treated spent overnight medium with morphology change and Cas12 expression.**

Fluorescence microscopy of Cas12-mCherry2 in the parental strain in a microfluidic plate. BHI was flown through for 1 h before switching to heat-treated spent overnight medium of parental strain for 4h. Afterwards, fresh BHI was flown through for 3 h. Three representative examples are shown (n = 2 biological replicates). Pictures were taken every 5 min. Same contrast settings as in **Fig. 4c-g**, **Supplementary Fig. 4d** and **Supplementary Movies 5** and **8**. Merge of phase contrast and RFP channel is shown. Fields of view are 13 x 13 µm. Scalebars represent 1 µm. Movies play at a frame rate of 7 frames per second.
