## Supplementary Figures and Tables for "MtvS1 and MtvS2 Interact with RNA Polymerase to Regulate the *Francisella* Type V-A CRISPR-Cas System"

### Supplementary Figures 1-6

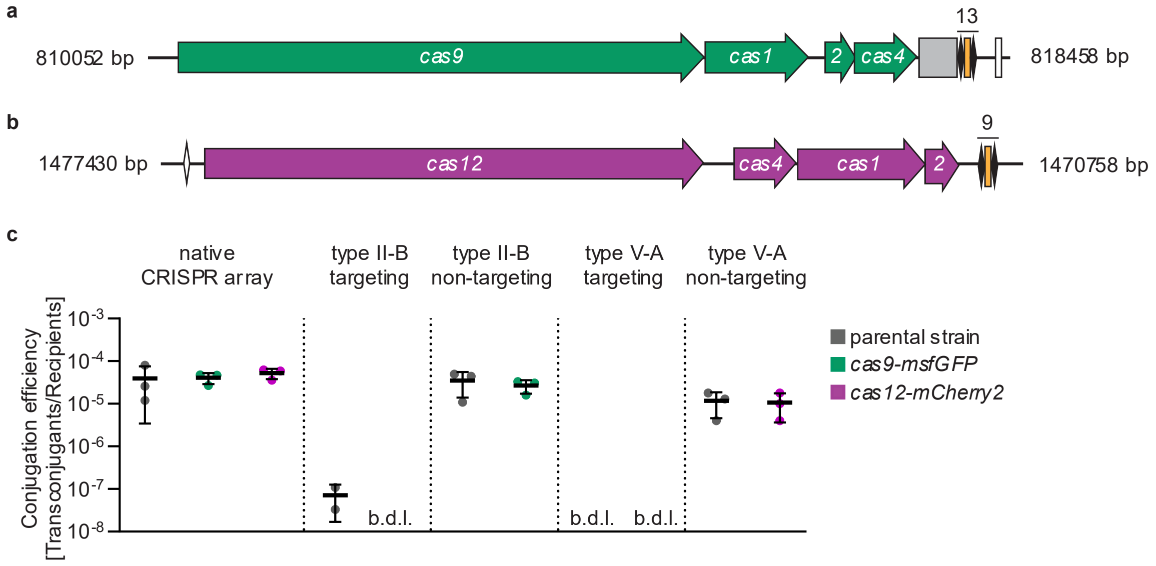


**Supplementary Figure 1**

**C-terminal fluorescent tags do not inhibit Cas9- and Cas12-mediated plasmid targeting.**

**a** Type II-B CRISPR-Cas system in *F. novicida U112*. Green arrows: *cas* genes. Grey box: *leader/tracr sequence*. Black diamonds: *repeats*. Orange box: *spacers*. White box: *scaRNA*. Drawn in scale.

**b** Type V-A CRISPR-Cas system in *F. novicida U112*. White diamond: *single repeat unit*. Magenta arrows: *cas* genes. Black diamonds: *repeats*. Orange box: *spacers*. Drawn in scale.

**c** Assessment of interference capability of wild-type and C-terminal tagged Cas9 and Cas12 in a conjugation-based plasmid inhibition assay. Strains have both native type II-B and type V-A CRISPR arrays or have the respective CRISPR array replaced by a spacer matching a protospacer on the conjugated plasmid (targeting spacer) or one that does not match (non-targeting spacer). Grey circles: parental strain. Green circles: *cas9-msfGFP*. Magenta circles: *cas12-mCherry2*. b.d.l.: below detection limit. Means with standard deviations are displayed (n = 3 biological replicates). Source data are provided as Source Data file.

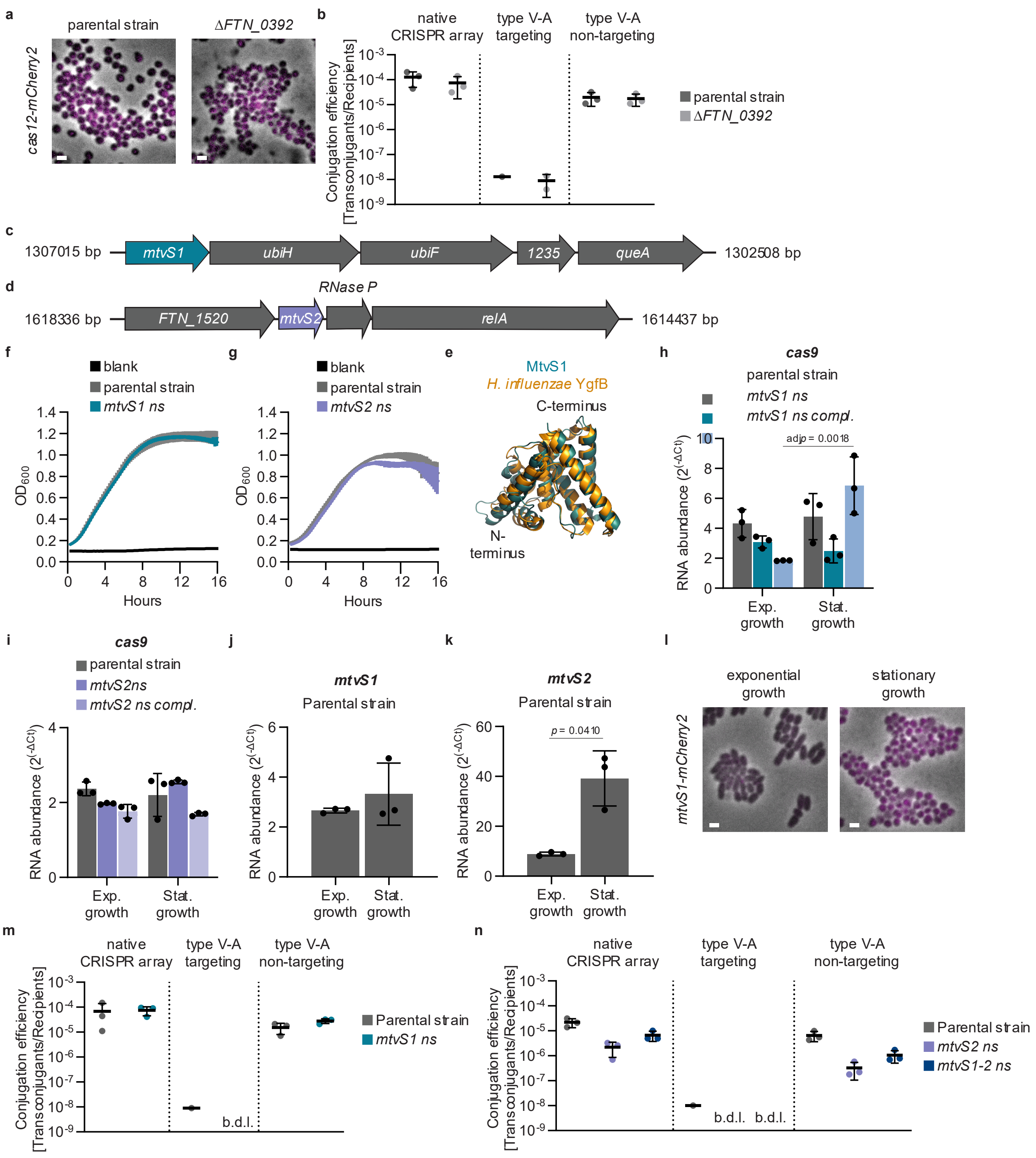


**Supplementary Figure 2**

**MtvS1 and MtvS2 have different expression patterns.**

**a** Fluorescence microscopy of Cas12-mCherry2 in parental strain and *ΔFTN_0392* mutant. Comparable fluorescent intensities in stationary growth were detected. Merge of phase contrast and RFP channel in 13 x 13 µm fields of view are shown. Scale bars represent 1 µm. Representative images are shown (n = 3 biological replicates). Source data are provided as Source Data file.

**b** Assessment of interference capability of parental strain and *ΔFTN_0392* mutant in a conjugation-based plasmid inhibition assay. Strains have a native type V-A CRISPR array or have a native type V-A CRISPR array replaced by a spacer matching a protospacer on the conjugated plasmid (targeting spacer) or one that does not match (non-targeting spacer). Grey circles: parental strain. Purple circles: *ΔFTN_0392* mutant. Means with standard deviations are displayed (n= 3 biological replicates). Source data are provided as Source Data file.
**c** *mtvS1* is the first gene of a conserved operon which also includes *ubiH* (2-octaprenyl-6-methoxyphenyl hydroxylase), *ubiF* (2-octaprenyl-3-methyl-6-methoxy-1,4-benzoquinol hydroxylase), *FTN_1235* (putative phage protein) and *queA* (S-adenosylmethionine:tRNA ribosyltransferase-isomerase).
**d** *mtvS2* is the second gene of a predicted operon which also includes *FTN_1520* (amino acid transporter), *RNAse P* RNA and *relA* (GTP pyrophosphokinase).
**e** Comparison of AlphaFold 3 MtvS1 structure (turquoise) with solved *H. influenzae* YgfB structure (yellow, 1IZM). RSMD value: 1.258 Å over 765 atoms.
**f-g** Growth curves of parental strain and *mtvS1 nonsense* **(f)** or *mtvS2 nonsense* **(g)** mutants. OD600 was measured every 10 min. Black: blank. Grey: parental strain. Turquoise: *mtvs1 nonsense* mutant. Purple: *mtvS2 nonsense* mutant. Means with standard deviations are displayed (n = 3 biological replicates with each 3 technical replicates). Source data are provided as Source Data file.
**h-i** Determination of mRNA levels for *cas9* in the *mtvS1 nonsense* **(h)** and *mtvS2 nonsense* **(i)** mutants by RT-qPCR. Grey bars: parental strain (with empty plasmid for **i**). Turquoise bars: *mtvS1 nonsense* mutant. Light blue bars: *mtvS1 nonsense* mutant with wild-type *mtvS1* at Tn7 insertion site. Purple bars: *mtvS2 nonsense* mutant with empty plasmid. Light purple bars: *mtvS2 nonsense* mutant with wild-type *mtvS2* expressed from plasmid. Means with standard deviations are displayed (n = 3 biological replicates). Significant differences were determined with a two-way ANOVA and Tukey's multiple comparison test (95 % confidence interval). Source data are provided as Source Data file.
**j-k** Determination of mRNA levels for *mtvS1* **(j)** and *mtvS2* **(k)** in the parental strain by RT-qPCR. Grey bars: parental strain. Means with standard deviations are displayed (n = 3 biological replicates). Significant differences were determined with Welch's *t*-test (95 % confidence interval). Source data are provided as Source Data file.
**l** Fluorescence microscopy of MtvS1-mCherry2 in parental strain. Fluorescence intensity is higher in stationary phase than in exponential growth phase. Merge of phase contrast and RFP channel in 13 x 13 µm fields of view are shown. Scale bars represent 1 µm. Representative images are shown (n = 3 biological replicates). Source data are provided as Source Data file.
**m-n** Assessment of interference capability of parental strain and *mtvS1 nonsense* **(m)**, *mtvS2 nonsense* and *mtvS1-2 nonsense* **(n)** mutants in a conjugation-based plasmid inhibition assay. Strains have a native type V-A CRISPR array or have the CRISPR array replaced by a spacer matching a protospacer on the conjugated plasmid (targeting spacer) or one that does not match (non-targeting spacer). Grey circles: parental strain. Turquoise circles: *mtvS1 nonsense* mutant. Purple circles: *mtvS2 nonsense* mutant. Blue circles: *mtvS1-2 nonsense* mutant. b.d.l.: below detection limit. Means with standard deviations are displayed (n = 3 biological replicates). Source data are provided as Source Data file.

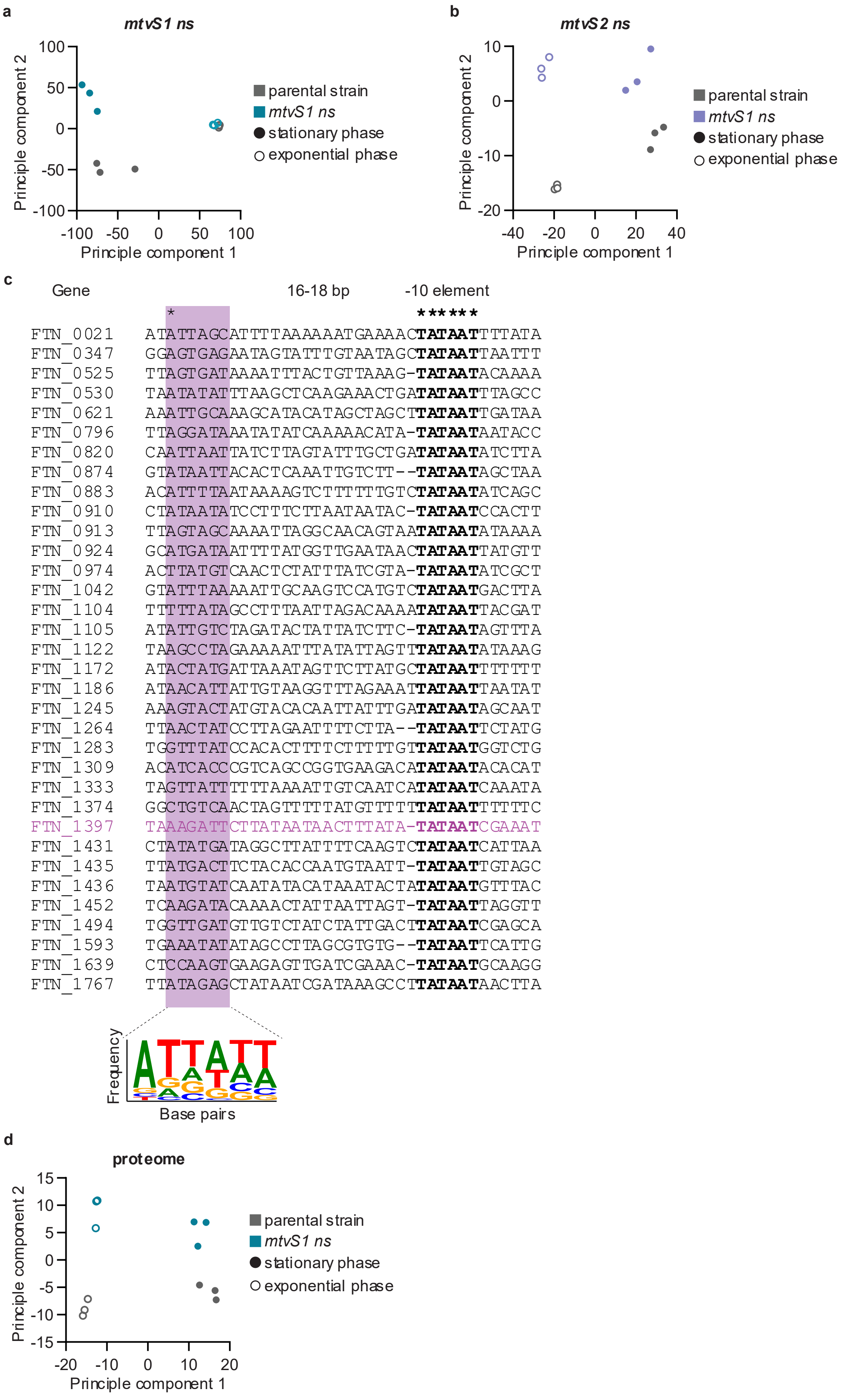
a
mtvS1 ns
Principle component 2
Principle component 1
100
50
0
-50
-100
-100 -50 0 50 100
parental strain
mtvS1 ns
stationary phase
exponential phase
b
mtvS2 ns
Principle component 2
Principle component 1
10
0
-10
-20
-40 -20 0 20 40
parental strain
mtvS1 ns
stationary phase
exponential phase
c
Gene
16-18 bp
-10 element
*
******
FTN_0021 ATATTAGCATTTTAAAAAATGAAAACTATAATTTTATA
FTN_0347 GGAGTGAGAATAGTATTTGTAATAGCTATAATTAATTT
FTN_0525 TTAGTGATAAAATTTACTGTTAAAG-TATAATACAAAA
FTN_0530 TAATATATTTAAGCTCAAGAAACTGATATAATTTAGCC
FTN_0621 AAATTGCAAAGCATACATAGCTAGCTTATAATTGATAA
FTN_0796 TTAGGATAAATATATCAAAAACATA-TATAATAATACC
FTN_0820 CAATTAATTATCTTAGTATTTGCTGATATAATATCTTA
FTN_0874 GTATAATTACACTCAAATTGTCTT--TATAATAGCTAA
FTN_0883 ACATTTTAATAAAAGTCTTTTTTGTCTATAATATCAGC
FTN_0910 CTATAATATCCTTTCTTAATAATAC-TATAATCCACTT
FTN_0913 TTAGTAGCAAAATTAGGCAACAGTAATATAATATAAAA
FTN_0924 GCATGATAATTTTATGGTTGAATAACTATAATTATGTT
FTN_0974 ACTTATGTCAACTCTATTTATCGTA-TATAATATCGCT
FTN_1042 GTATTTAAAAATTGCAAGTCCATGTCTATAATGACTTA
FTN_1104 TTTTTATAGCCTTTAATTAGACAAAATATAATTACGAT
FTN_1105 ATATTGTCTAGATACTATTATCTTC-TATAATAGTTTA
FTN_1122 TAAGCCTAGAAAAATTTATATTAGTTTATAATATAAAG
FTN_1172 ATACTATGATTAAATAGTTCTTATGCTATAATTTTTTT
FTN_1186 ATAACATTATTGTAAGGTTTAGAAATTATAATTAATAT
FTN_1245 AAAGTACTATGTACACAATTATTTGATATAATAGCAAT
FTN_1264 TTAACTATCCTTAGAATTTTCTTA--TATAATTCTATG
FTN_1283 TGGTTTATCCACACTTTTCTTTTTGTTATAATGGTCTG
FTN_1309 ACATCACCCGTCAGCCGGTGAAGACATATAATACACAT
FTN_1333 TAGTTATTTTTTAAAATTGTCAATCATATAATCAAATA
FTN_1374 GGCTGTCAACTAGTTTTTATGTTTTTATAATTTTTTC
FTN_1397 TAAAGATTCTTATAATAACTTTATA-TATAATCGAAAT
FTN_1431 CTATATGATAGGCTTATTTTCAAGTCTATAATCATTAA
FTN_1435 TTATGACTTCTACACCAATGTAATT-TATAATTGTAGC
FTN_1436 TAATGTATCAATATACATAAATACTATATAATGTTTAC
FTN_1452 TCAAGATACAAAACTATTAATTAGT-TATAATTAGGTT
FTN_1494 TGGTTGATGTTGTCTATCTATTGACTTATAATCGAGCA
FTN_1593 TGAAATATATAGCCTTAGCGTGTG--TATAATTCATTG
FTN_1639 CTCCAAGTGAAGAGTTGATCGAAAC-TATAATGCAAGG
FTN_1767 TTATAGAGCTATAATCGATAAAGCCTTATAATAACTTA
Frequency
Base pairs
d
proteome
Principle component 2
Principle component 1
15
10
5
0
-5
-10
-15
-20 -10 0 10 20
parental strain
mtvS1 ns
stationary phase
exponential phase

**Supplementary Figure 3**
**Analysis of *F. novicida* U112 transcriptomic and proteomics data.**
**a-b** Principal component analysis of parental strain and *mtvs1 nonsense* **(a)** and *mtvS2 nonsense* **(b)** mutants. Grey: parental strain. Turquoise: *mtvs1 nonsense* mutant. Purple: *mtvS2 nonsense* mutant. Filled circles: stationary phase samples. Empty circles: exponential phase samples. n = 3 biological replicates. Source data are provided as Source Data file.
**c** Promoter sequences of significantly downregulated operons for both the *mtvs1 nonsense* and *mtvS2 nonsense* mutants in stationary phase with TATAAT motifs were aligned to TATAAT motif (in bold). First genes of operons indicated. Bold stars: 100 % conserved sites. Star: 79 % conserved site. Magenta: promoter sequence of *type V-A CRISPR-Cas system*. Web logo plot: frequency of base pairs 16-18 bp upstream of TATAAT motif.
**d** Principal component analysis of parental strain and *mtvs1 nonsense* mutant. Grey: parental strain. Turquoise: *mtvs1 nonsense*. Filled circles: stationary phase samples. Empty circles: exponential phase samples. n = 3 biological replicates. Source data are provided as Source Data file.

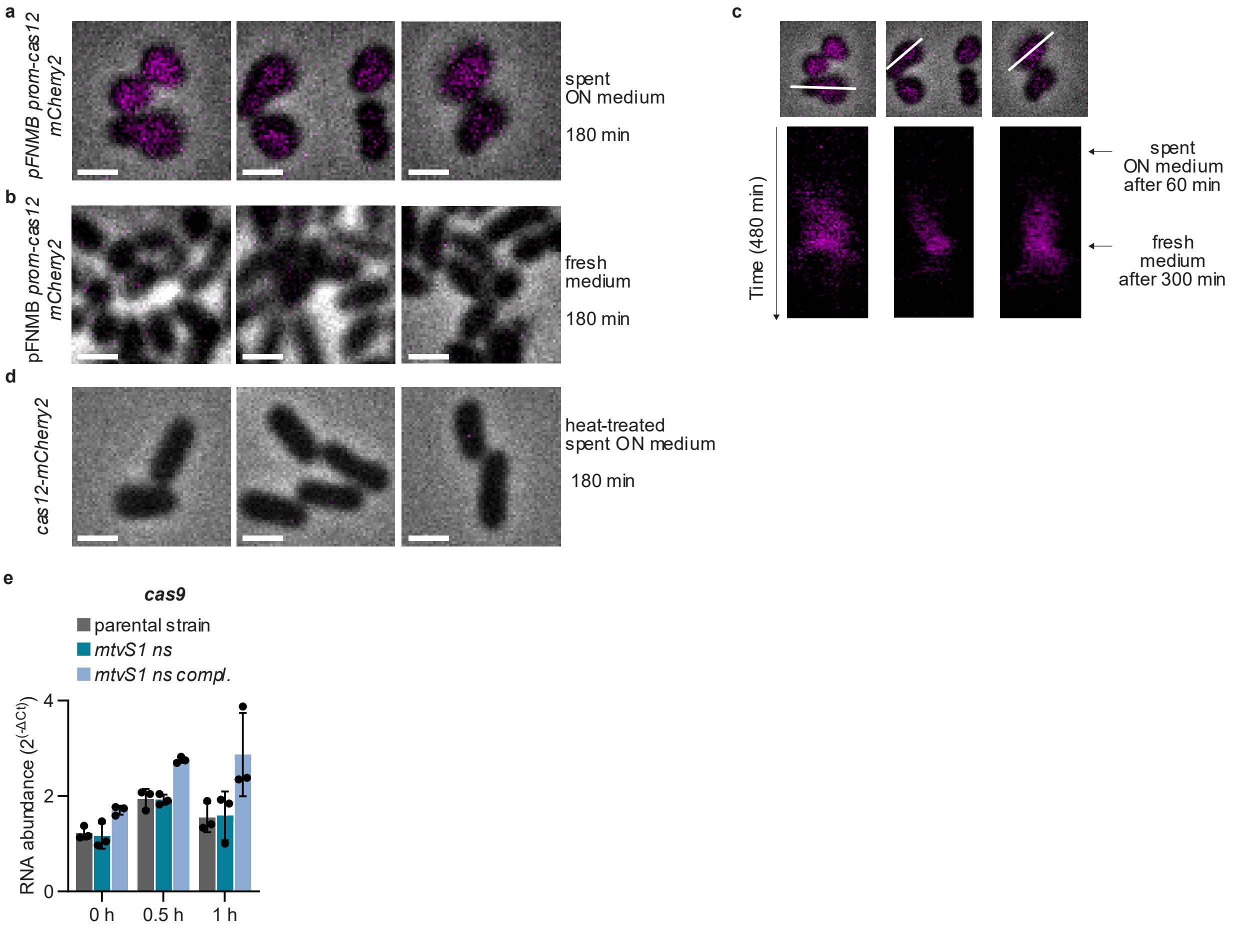


**Supplementary Figure 4**

**Spent overnight medium is heat sensitive and does not affect *cas9* expression.**

**a-b** Fluorescence microscopy of mCherry2 from *cas12* promoter on plasmid in a microfluidic plate. BHI was flown through for 1 h before switching to spent overnight medium of parental strain for 4h. Afterwards fresh BHI was flown through for 3 h. **(a)** Three examples of 240 min time point (180 min after switching to spent ON medium) and **(b)** three examples of 480 min (180 min after switching back to fresh BHI) are shown. Merge of phase contrast and RFP channel in 4 x 4 µm fields of view are shown. Same contrast settings as in **c** and **Supplementary Movie 6**. Scale bars represent 1 µ m. Representative images are shown (n = 2 biological replicates). Source data are provided as Source Data file.

**c** Examples of **a** and **b** were used to create time-resolved kymographs of RFP channel. Merge of phase contrast and RFP channel is shown (upper images). White lines show axis along which the kymographs of RFP channel were made (lower images). Same contrast settings as in **a-b** and **Supplementary Movie 6**. Representative images are shown (n = 2 biological replicates). Source data are provided as Source Data file.

**d** Fluorescence microscopy of Cas12-mCherry2 in the parental strain in a microfluidic plate. BHI was flown through for 1 h before switching to heat-treated spent overnight medium of parental strain for 4h. Afterwards fresh BHI was flown through for 3 h. Three examples of 240 min time point (180 min after switching to spent ON medium) are shown. Merge of phase contrast and RFP channel in 4 x 4 µm fields of view are shown. Same contrast settings as in **Fig. 4c-g** and **Supplementary Movies 5, 7-8**. Scale bars represent 1 µm. Representative images are shown (n = 2 biological replicates). Source data are provided as Source Data file.

**e** Determination of mRNA levels for *cas9* after switching to spent overnight medium by RT-qPCR. Samples were taken 0 h, 0.5 h and 1 h after medium change. Grey bars: parental strain. Turquoise bars: *mtvS1 nonsense* mutant. Light blue bars: *mtvS1 nonsense* mutant with wild-type *mtvS1* at Tn7 insertion site. Means with standard deviations are displayed (n = 3 biological replicates). Significant differences were determined with a two-way ANOVA and Tukey's multiple comparison test (95 % confidence interval). Source data are provided as Source Data file.

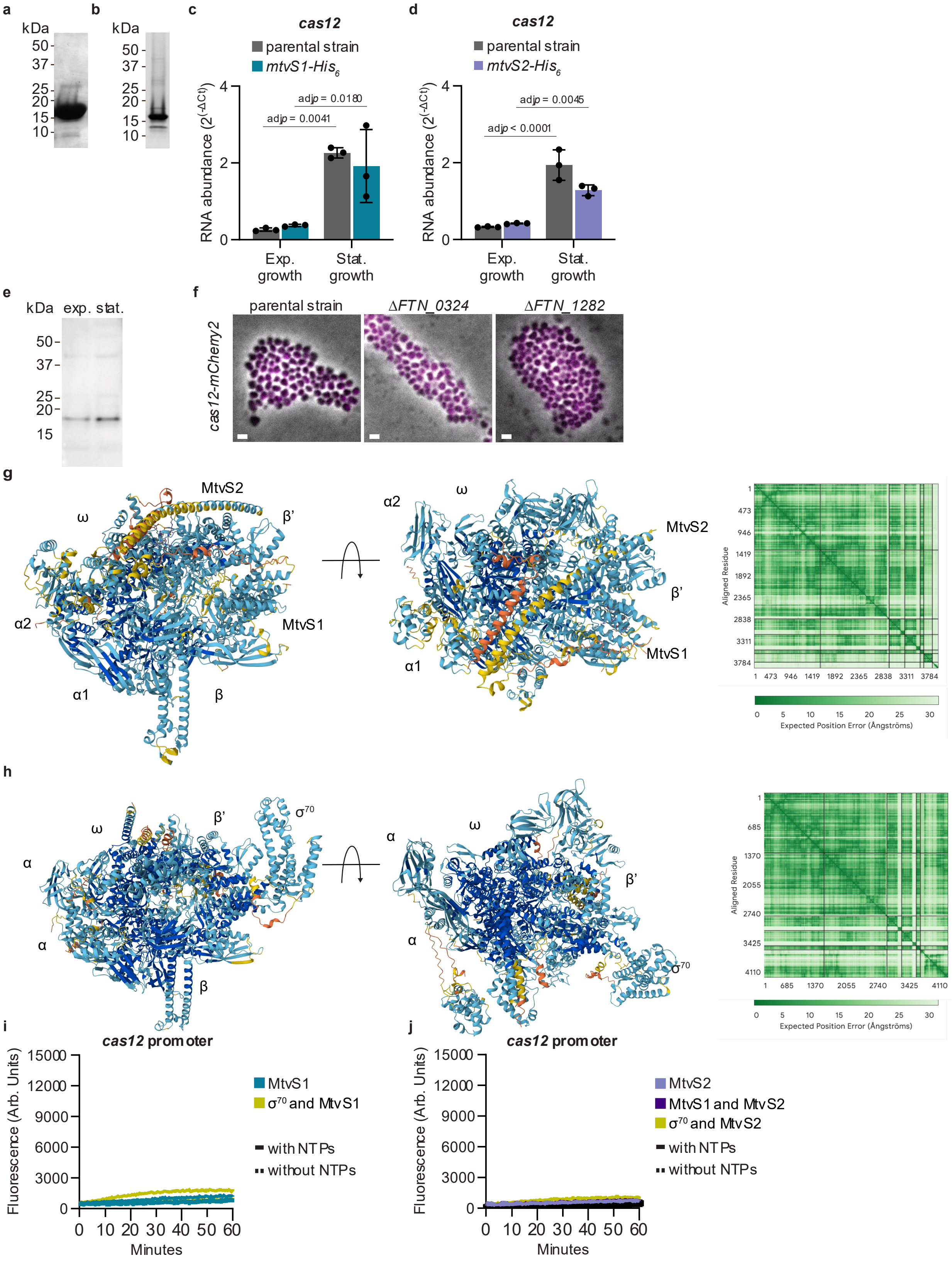
a
kDa
50
37
25
20
15
10
b
kDa
50
37
25
20
15
10
c
cas12
parental strain
mtvS1-His6
RNA abundance (2^(-ΔCt))
adjp = 0.0041
adjp = 0.0180
Exp. growth
Stat. growth
d
cas12
parental strain
mtvS2-His6
RNA abundance (2^(-ΔCt))
adjp < 0.0001
adjp = 0.0045
Exp. growth
Stat. growth
e
kDa
exp.
stat.
50
37
25
20
15
f
parental strain
ΔFTN_0324
ΔFTN_1282
cas12-mCherry2
g
MtvS2
ω
β'
α2
MtvS1
α1
β
Aligned Residue
Expected Position Error (Ångströms)
h
ω
β'
σ70
α
β
Aligned Residue
Expected Position Error (Ångströms)
i
cas12 promoter
Fluorescence (Arb. Units)
Minutes
MtvS1
σ70 and MtvS1
with NTPs
without NTPs
j
cas12 promoter
Fluorescence (Arb. Units)
Minutes
MtvS2
MtvS1 and MtvS2
σ70 and MtvS2
with NTPs
without NTPs

**Supplementary Figure 5**
**MtvS1-His$_6$ and MtvS2-His$_6$ purification and AlphaFold 3 predictions.**
**a** SDS-PAGE gel of MtvS1-His$_6$ purified with size-exclusion chromatography from *E. coli*. Representative image is shown (n = 2.) Source data are provided as Source Data file.
**b** SDS-PAGE gel of MtvS2-His$_6$ after dialysis after purification with 8 M urea from *E. coli*. Representative image is shown (n = 2). Source data are provided as Source Data file.
**c-d** Determination of mRNA levels for *cas12* by RT-qPCR. Grey bars: parental strain. Turquoise bars: *mtvS1-His$_6$* strain. Purple bars: *mtvS2-His$_6$* strain. Means with standard deviations are displayed (n = 3 biological replicates). Significant differences were determined with a two-way ANOVA and Tukey's multiple comparison test (95 % confidence interval). Source data are provided as Source Data file.
**e** Western blot probing for chromosomal MtvS1-His$_6$ in exponential growth and stationary phase lysates. n = 1. Source data are provided as Source Data file.
**f** Fluorescence microscopy of Cas12-mCherry2 in parental strain, *ΔFTN_0324* and *ΔFTN_1282* mutant. Comparable fluorescent intensities in stationary growth were detected. Merge of phase contrast and RFP channel in 13 x 13 µm fields of view are shown. Scale bars represent 1 µm. Representative images are shown (n = 2 biological replicates). Source data are provided as Source Data file.
**g-h** AlphaFold 3 stats for predictions of MtvS1 (A0A6I4RTR9) and MtvS2 (A0Q819) with *F. novicida* U112 RNA polymerase (RpoA1: A0Q4K8, RpoA2: A0Q7R6, RpoB: A0Q867, RpoC: A0Q866 and RpoZ: A0Q5J3) **(g)** and of *E. coli* $\sigma^{70}$ (RpoD, P00579) with *E. coli* RNA polymerase (RpoB: P0A8V2, RpoC: P0A8T7, RpoA: P0A7Z4 and RpoZ: P0A800) (**h**). Models with pLDDT score overlaid from two viewing angles. Color code: dark blue, very high confidence; blue, confident; yellow, low confidence; orange, very low confidence. The predicted aligned error (PAE) plot is shown on the right. Overall prediction scores: ipTM = 0.79 and pTM = 0.82 **(g)** and ipTM = 0.82 and pTM = 0.84 **(h)**.
**i-j** Fluorescence based *in vitro* transcription assay with *cas12* promoter aptamer and *E. coli* RNA polymerase core enzyme with MtvS1-His$_6$ (turquoise) and with both MtvS1-His$_6$ and $\sigma^{70}$ (light green) **(i)** or with MtvS2-His$_6$ (purple, both MtvS1-His$_6$ and MtvS2-His$_6$ (dark purple) or with $\sigma^{70}$ (light green) **(j)**.Representative assays (mean and standard deviation, n = 3 technical replicates, repeated twice) with (solid line) and without (dashed line) NTPs added are shown. Source data are provided as Source Data file.

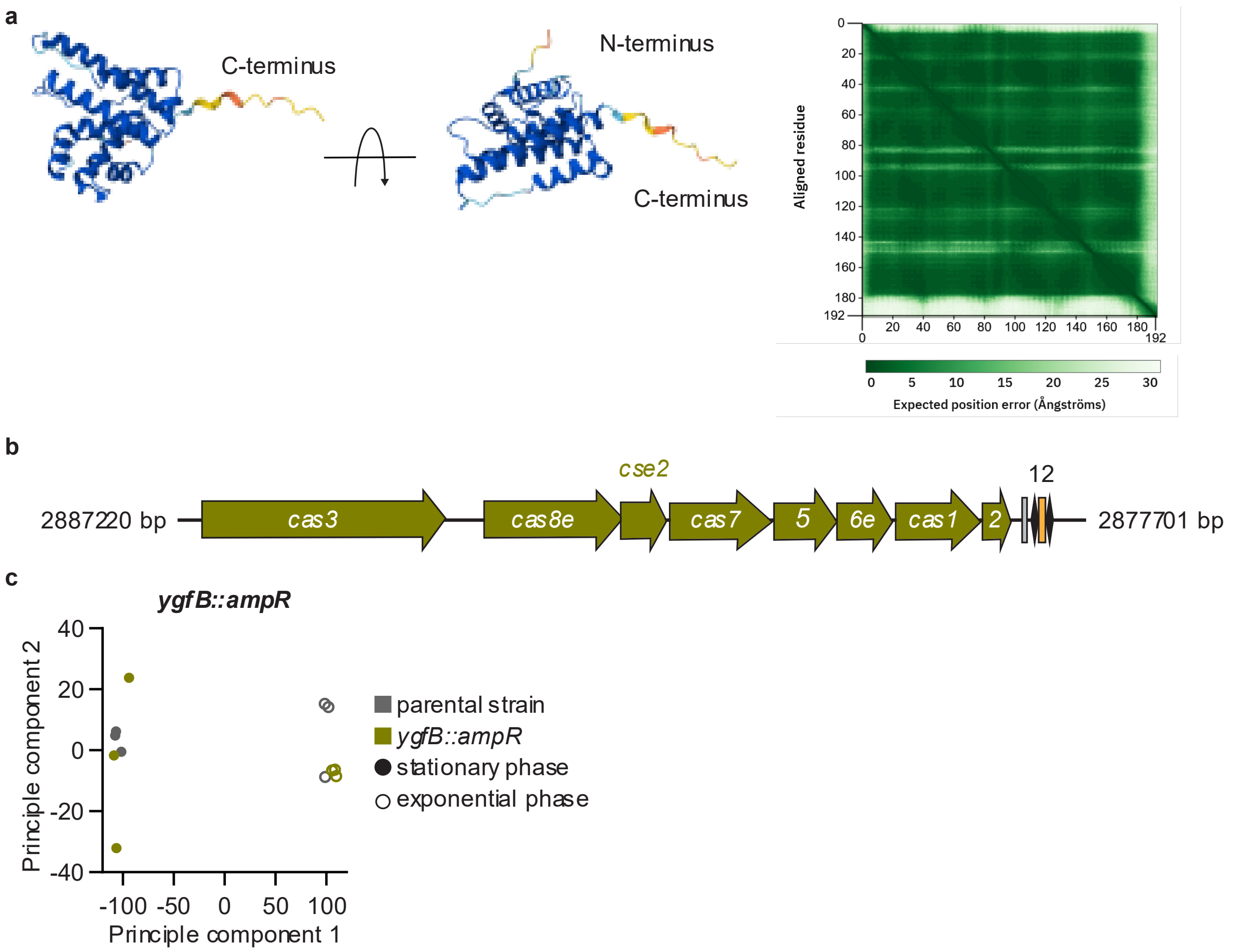


**Supplementary Figure 6**

**Principal component analysis for *E. coli* transcriptomic data.**

**a** AlphaFold 3 stats for predictions of *E. coli* YgfB (A8A450). Model with pLDDT score overlaid from two viewing angles. Color code: dark blue, very high confidence; blue, confident; yellow, low confidence; orange, very low confidence. The predicted aligned error (PAE) plot is shown on the right.

**b** Type I-E CRISPR-Cas system in *E.coli* K-12 substr. BW25113. Olive green: *cas* and *cse* genes. Grey box: *leader/tracr sequence*. Black diamonds: *repeats*. Orange box: *spacers*. Drawn in scale.

**c** Principal component analysis of parental strain and *ygfB::ampR* mutant. Grey: parental strain. Olive green: *ygfB::ampR* mutant. Filled circles: stationary phase samples. Empty circles: exponential phase samples. n = 3 biological replicates. Source data are provided as Source Data file.

### Supplementary Tables

#### Supplementary Table 1: Predicted interactions under 3 Å between RpoC and MtvS1.

| RpoC (amino acid) | Type of interaction | MtvS1 (amino acid) |
|---|---|---|
| E40 | side chain carboxyl hydrogen bond with backbone carbonyl | T52 |
| T41 | backbone carbonyl carbon interaction with lysine amine | K53 |
| I42 | backbone NH hydrogen bond with lysine amine | K53 |
| N43 | side chain amide nitrogen hydrogen bond with glutamic acid side chain carboxyl | E56 |
| Y44 | side chain phenol hydrogen bond with backbone carbonyl | E56 |
| Y44 | side chain phenol hydrogen bond with backbone carbonyl | E57 |
| R45 | side chain guanidinium hydrogen bond with carbonyl | E57 |
| R268 | side chain guanidinium hydrogen bond with side chain amide | Q19 |
| R276 | side chain guanidinium hydrogen bond with side chain carboxyl | E26 |
| P286 | proline ring carbon interaction with backbone carbonyl | L121 |
| I288 | backbone carbonyl interaction with side chain carbon | M139 |
| E293 | side chain carboxyl hydrogen bond to serine hydroxyl | S114 |
| R295 | side chain guanidinium hydrogen bond with backbone carbonyl | S142 |
| R295 | side chain guanidinium interactions with side chain phenol carbons | Y143 |
| E299 | side chain carboxyl hydrogen bond with side chain guanidinium | R110 |
| R312 | side chain carbons and backbone NH interactions with side chain carboxyl | E147 |
| S317 | serine hydroxyl hydrogen bond to lysine amine | K16 |

### Supplementary Table 2: Strains used in this study.

| Organism | Genotype | Plasmid | Relevant features | Source |
|---|---|---|---|---|
| *Francisella novicida U112* | | | Parental strain | Bei resources (NR-13) |
| | *typeII_tet_spacer_targ* | | Type II-B wild-type CRISPR array was replaced with targeting spacer matching *tetR* on plasmid pFNMB2 | This study |
| | *typeII_tet_spacer_nontarg* | | Type II-B wild-type CRISPR array was replaced with non-targeting spacer matching *tetR* on plasmid pFNMB2 | This study |
| | *typeV_tet_spacer_targ* | | Type V-A wild-type CRISPR array was replaced with targeting spacer matching *tetR* on plasmid pFNMB2 | This study |
| | *typeV_tet_spacer_nontarg* | | Type V-A wild-type CRISPR array was replaced with non-targeting spacer matching *tetR* on plasmid pFNMB2 | This study |
| | Δ*FTN_0710*, Δ*FTN_1487* | | Deletion of *FTN_0710* and *FTN_1487* | This study |
| | | pFNMB2 *mcs* | Empty vector control | This study |
| | | pFNMB *prom-cas12 mCherry2* | mCherry2 expressed from *cas12* promoter | This study |
| | *mtvS1 nonsense* | | Three stop codons introduced at position 11 of *mtvs1* | This study |
| | *mtvS1 nonsense*, *Tn7_mtvS1* | | Wild-type *mtvS1* with endogenous promoter was introduced at the canonical Tn7 insertion site downstream of *glmS* | This study |
| | *mtvS1 nonsense*, *typeV_tet_spacer_targ* | | Type V-A wild-type CRISPR array was replaced with targeting spacer matching *tetR* on plasmid pFNMB2 | This study |
| | *mtvs1 nonsense*, *typeV_tet_spacer_nontarg* | | Type V-A wild-type CRISPR array was replaced with non-targeting spacer matching *tetR* on plasmid pFNMB2 | This study |
| | *mtvS1-mCherry2* | | C-terminal chromosomal fusion of *mCherry2* to *mtvS1* | This study |
| | *mtvS1-His*$_6$ | | C-terminal chromosomal His$_6$-tag with a glycine linker | This study |
| | Δ*FTN_0392*, *typeV_tet_spacer_targ* | | Type V-A wild-type CRISPR array was replaced with targeting spacer matching *tetR* on plasmid pFNMB2 | This study |
| | Δ*FTN_0392*, *typeV_tet_spacer_nontarg* | | Type V-A wild-type CRISPR array was replaced with non-targeting spacer matching *tetR* on plasmid pFNMB2 | This study |
| | *mtvS2 nonsense* | | Three stop codons introduced at position 11 of *mtvs2* | This study |
| | *mtvS2 nonsense* | pFNMB2 *mcs* | Empty vector control | This study |
| | *mtvS2 nonsense* | pFNMB *prom-upstream mtvS2* | Wild-type *mtvS2* with endogenous promoter upstream of gene on plasmid | This study |
| | *mtvS1 nonsense, mtvS2 nonsense* | | Three stop codons introduced at position 11 of *mtvs1* and *mtvS2* | This study |
| | *mtvS1 nonsense, mtvS2 nonsense,* Δ*FTN_0710,* Δ*FTN_1487* | | Three stop codons introduced at position 11 of *mtvs1* and *mtvS2* and deletion of *FTN_0710* and *FTN_1487* | This study |
| | *mtvS2-His*$_6$ | | C-terminal chromosomal His$_6$-tag with a glycine linker | This study |
| | *cas9-msfGFP*, *cas12-mCherry2* | | C-terminal chromosomal fusion of *msfgfp* to *cas9* and c-terminal chromosomal fusion of *mCherry2* to *cas12* | This study |
| | *cas9-msfGFP*, *cas12-mCherry2*, *typeV_tet_spacer_targ* | | Type V-A wild-type CRISPR array replaced with targeting spacer matching *tetR* on plasmid pFNMB2 | This study |
| | *cas9-msfGFP*, *cas12-mCherry2*, *typeV_tet_spacer_nontarg* | | Type V-A wild-type CRISPR array replaced with non-targeting spacer matching *tetR* on plasmid pFNMB2 | This study |
| | *cas9-msfGFP*, *cas12-mCherry2*, *typeII_tet_spacer_targ* | | Type II-B wild-type CRISPR array replaced with targeting spacer matching *tetR* on plasmid pFNMB2 | This study |
| | *cas9-msfGFP*, *cas12-mCherry2*, *typeII_tet_spacer_nontarg* | | Type II-B wild-type CRISPR array replaced with non-targeting spacer matching *tetR* on plasmid pFNMB2 | This study |

| | | | | |
|---|---|---|---|---|
| | *cas9-msfGFP*, *cas12-mCherry2*, *mtvs1 nonsense* | | Three stop codons introduced at position 11 of *mtvS1* | This study |
| | *cas9-msfGFP*, *cas12-mCherry2*, Δ*FTN_0324* | | Deletion of *FTN_0324* | This study |
| | *cas9-msfGFP*, *cas12-mCherry2*, Δ*FTN_0392* | | Deletion of *FTN_0392* | This study |
| | *cas9-msfGFP*, *cas12-mCherry2*, Δ*FTN_1282* | | Deletion of *FTN_1282* | This study |
| | *cas12::mCherry2* | | *cas12* was replaced with *mCherry2* | This study |
| *Escherichia coli* K-12 *substr.* BW25113 | | | | Coli Genetic Stock Center at Yale University and [1] |
| | | pAM38 (*red*) | λ Red recombineering plasmid | [2] |
| | | pFNMB2 *mcs* | Empty vector control | This study |
| | *ygfB::ampR* | | *ygfB* disrupted by ampicillin resistance cassette | This study |
| | *ygfB::ampR* | pFNMB2 *mcs* | Empty vector control | This study |
| | *ygfB::ampR* | pFNMB *prom mtvS1* | Wild-type *mtvS1* with endogenous promoter on plasmid | This study |
| | *ygfB::ampR* | pFNMB *prom ygfB* | Wild-type *ygfB* with endogenous promoter on plasmid | This study |
| *Escherichia coli OneShotpir* | | | Cloning strain | ThermoFisher |
| *Escherichia coli JKE 201* | | | Donor *E. coli* used for conjugation into *F. novicida* | [3] |
| | | pFNMB2 *msfGFP* | Donor *E. coli* containing mobilizable *Francisella* plasmid pFNMB2 *msfGFP* | [4] |
| | | pFNMB2 *msfGFP* *sp3_typeV_seedG3T_targ* *sp5_typeV_seedA3C_targ* | Plasmid containing protospacers 3 and 5 of native Type V CRISPR array carrying seed mutations G to T and A to C at position 3, respectively, and TTTG PAM | This study |
| *Escherichia coli Rosetta (DE3)* | | | Overexpression strain | Novagen |
| | | pET21 *mtvS1-His*$_6$ | Expression of MtvS1-$His_6$ | This study |
| | | pET21 *mtvS2-His*$_6$ | Expression of MtvS2-$His_6$ | This study |
| *Escherichia coli BL21 (DE3)* | | pVS11/ p*Ec*rpo*ABC(-XH) Z)*, pACYC-Duet1_*EcoRpoZ* | Overexpression of *E. coli* RNA polymerase | [5,6] |
| | | pSAD1403 | Overexpression of *E. coli* $\sigma^{70}$ | [7] |

### Supplementary Table 3: Plasmids, antibiotic resistance casettes and primers used to generate mutants.

| Plasmid Name | Spacer/Peptide sequence | Primers | Sequence 5'-3' [base pairs] |
|---|---|---|---|
| pDMK3 *cas9-msfGFP* | | FTN_0757-fusion_1.FOR | ATTGTCGACCCTCGAGCAAGCAGGTTTTGGATAAG |
| | | FTN_0757-fusion_1.REV | GCGGCCGCATTATTAGATGTTTCATTATAAATACCT |
| | | FTN_0757-fusion_2.FOR | TCTAATAATGCGGCCGCAGGA |
| | | FTN_0757-msfGFP_2.REV | TACCAAGCATTTTGTAGAGCTCATCCATG |
| | | FTN_0757-msfGFP_3.FOR | GCTCTACAAAATGCTTGGTATGAAATTAG |
| | | FTN_0757-fusion_3.REV | ACAATTTGTGGAATTCCCGGGCGATACTCCTCCATTAGA |
| | | FTN_0757-fusion_Det.FOR | TCTAAAAACCCAAAAATTACATGAG |
| | | FTN_0757-fusion_Det.REV | TGTTCACCAGCAAATTCTCCAG |
| pDMK3 *cas12-mCherry2* | | FTN_1397-fusion_1.FOR | CTGAATTGTCGACCCTCGAGCAAAAATATTAACAAAAACTATATTCTTT |
| | | FTN_1397-mCherry2_1.REV | TGTACAAGTTATTTGAGTTCGTGCAGA |
| | | FTN_1397-mCherry2_2.FOR | ACGAACTCAAATAACTTGTACAGCTCGT |
| | | FTN_1397-fusion_2.REV | GAATAACGCGGCCGCAGGA |
| | | FTN_1397-fusion_3.FOR | CGGCCGCGTTATTCCTATTCTGCACG |
| | | FTN_1397-fusion_3.REV | CAATTTGTGGAATTCCCGGGGTGCTTAGAGCTTATCAG |
| | | FTN_1397-fusion_Det.FOR | TCTCTTGCTGCTCTGGAGTT |
| | | FTN_1397-fusion_Det.REV | AAAAGAGGGCGTTTCAAGGT |
| pDMK3 *cas12::mCherry2* | | FTN_1397::mCherry2_1.FOR | CTTCTGAATTGTCGACCCTCGAGAGTCCGTTTTATCACATAAG |
| | | FTN_1397::mCherry2_1.REV | TGTACAAGTAAGGTAGGATCAAAAATAATCA |
| | | FTN_1397::mCherry2_2.FOR | TGATCCTACCTTACTTGTACAGCTCGTC |
| | | FTN_1397::mCherry2_2.REV | AGTCTTTATCATGGTGAGCAAGGGC |
| | | FTN_1397::mCherry2_3.FOR | GCTCACCATGATAAAGACTCCTTATAAAATT |
| | | FTN_1397::mCherry2_3.REV | TCAGTAGCGGCCGCATTTTGATAAAGGTAAAAGAGC |
| | | FTN_1397::mCherry2_Det.FOR | TCTCTTGCTGCTCTGGAGTT |
| | | FTN_1397::mCherry2_Det.REV | ACACTTGCTGATGTTGGTGA |
| pDMK3 *typeII_tet_spacer_targ* | TCGCGATGACTTAGTAAAGCACATC TAAAACTTT | TypeII_spacer_1.FOR | CTGAATTGTCGACCCTCGAGATAGTTGACTGGGTTATAG |
| | | TypeII_spacer_tet_targ_1.REV | AAAGTTTTAGATGTGCTTTACTAAGTCATCGCGAGTTTCAGTTGCTGAATT ATTTGGTAAACTACTGTTAGAGCCAAAACAAACCTAGTGT |
| | | TypeII_spacer_tet_targ_2.FOR | TCGCGATGACTTAGTAAAGCACATCTAAAACTTTCTAACAGTAGTTTACCA AATAATTCAGCAACTGAAACTTAATTAATTTTGTAAATTA |
| | | TypeII_spacer_2.REV | ATCTATCTAGAAGGGCCCCGTTTCACTAAATTCTTA |
| | | TypeII_spacer_Det.FOR | ATGGGTTGCTCAAGGTGAAG |
| | | TypeII_spacer_Det.REV | TCTCATGCTGGTTACGCTTG |
| pDMK3 *typeII_tet_spacer_nontarg* | AAAGTTTTAGATGTGCTTTACTAAG TCATCGCGA | TypeII_spacer_1.FOR | CTGAATTGTCGACCCTCGAGATAGTTGACTGGGTTATAG |
| | | TypeII_spacer_tet_nontarg_1.REV | TCGCGATGACTTAGTAAAGCACATCTAAAACTTTGTTTCAGTTGCTGAATT ATTTGGTAAACTACTGTTAGAGCCAAAACAAACCTAGTGT |
| | | TypeII_spacer_tet_nontarg_2.FOR | AAAGTTTTAGATGTGCTTTACTAAGTCATCGCGACTAACAGTAGTTTACCA AATAATTCAGCAACTGAAACTTAATTAATTTTGTAAATTA |
| | | TypeII_spacer_2.REV | ATCTATCTAGAAGGGCCCCGTTTCACTAAATTCTTA |
| | | TypeII_spacer_Det.FOR | ATGGGTTGCTCAAGGTGAAG |
| | | TypeII_spacer_Det.REV | TCTCATGCTGGTTACGCTTG |

| | | | |
|---|---|---|---|
| pDMK3 *typeV_tet_spacer_targ* | TCTAACATCTCAATGGCTAAGGCGTCGAG | TypeV_spacer_1.FOR | TCTATCGCCTTCTTGACGAGTTCTTCTGAATTGTCGACCCTCGAGATGCTGTTTCCAAGC |
| | | TypeV_spacer_tet_targ_1.REV | CTCGACGCCTTAGCCATTGAGATGTTAGAGTCTAAGAACTTTAAATAATTTGTCTGTATATTATTGATTTCTAAATTAGAATTTT |
| | | TypeV_spacer_tet_targ_2.FOR | TCTAACATCTCAATGGCTAAGGCGTCGAGATCTACAACAGTAGAAATTATTTAAAGTTCTTAGACCCGTTTTTGCCTAAATCAGC |
| | | TypeV_spacer_2.REV | TTGTGAGCGGATAACAATTTGTGGAATTCCCGGGAGAGCTCCCTAGTTTGTGACTTTGTT |
| | | TypeV_spacer_Det.FOR | GCCAGCGAATAAAAGTCCTG |
| | | TypeV_spacer_Det.REV | CAAGCCTAGCGGAGCTAATG |
| pDMK3 *typeV_tet_spacer_nontarg* | CTCGACGCCTTAGCCATTGAGATGTTAGA | TypeV_spacer_1.FOR | TCTATCGCCTTCTTGACGAGTTCTTCTGAATTGTCGACCCTCGAGATGCTGTTTCCAAGC |
| | | TypeV_spacer_tet_nontarg_1.REV | TCTAACATCTCAATGGCTAAGGCGTCGAGGTCTAAGAACTTTAAATAATTTGTCTGTATATTATTGATTTCTAAATTAGAATTTT |
| | | TypeV_spacer_tet_nontarg_2.FOR | CTCGACGCCTTAGCCATTGAGATGTTAGAATCTACAACAGTAGAAATTATTTAAAGTTCTTAGACCCGTTTTTGCCTAAATCAGC |
| | | TypeV_spacer_2.REV | TTGTGAGCGGATAACAATTTGTGGAATTCCCGGGAGAGCTCCCTAGTTTGTGACTTTGTT |
| | | TypeV_spacer_Det.FOR | GCCAGCGAATAAAAGTCCTG |
| | | TypeV_spacer_Det.REV | CAAGCCTAGCGGAGCTAATG |
| pDMK3 *mtvS1 nonsense* | MKNEKPNFED***ALK | FTN_1238_ns_1.FOR | CTTCTGAATTGTCGACCCTCGAGCCAAATAAGCCTACTCCT |
| | | FTN_1238_ ns_1.REV | AGATTAGTAGTAGGCATTAAAAGTAATGCAA |
| | | FTN_1238_ns_2.FOR | TACTTTTAATGCCTACTACTAATCTTCAAAATTAGGTTTTT |
| | | FTN_1238_ns_2.REV | ATAACAATTTGTGGAATTCCCGGGAGAGCTCAGGTTACGCGGCCGCCCACCCATGCCCATT |
| | | FTN_1238_ns_Det.FOR | CCAACTATCCCACCCCCAAC |
| | | FTN_1238_ns_Det.REV | TTTGCTGCTCATCAAAGGCT |
| | | FTN_1238_ns_Det2.FOR | CCTACTACTAATCTTCAAAATTAG |
| pDMK3 *Tn7_mtvS1* | | FTN_0485_Tn7_1.FOR | CTTCTGAATTGTCGACCCTCGAGAATGTGCCAAATAGTTCG |
| | | FTN_1238_Tn7_1.REV | TTTAGGTATTAACTTGGAAAATAAATGATAAGAAAGTAAAA |
| | | FTN_1238_Tn7_2.FOR | TTTATTTTCCAAGTTAATACCTAAATTTTTTCACTAT |
| | | FTN_1238_Tn7_2.REV | ATTTAGCTAAGTTTAAACTCTTGATTACATAGCC |
| | | FTN_1238_Tn7_3.FOR | ATCAAGAGTTTAAACTTAGCTAAATCTGTAACC |
| | | FTN_0485_Tn7_3.REV | AGAGCTCAGGTTACGCGGCCGCGTATCAAGCAAGAGCTTAG |
| | | FTN_0485_Tn7_Det.FOR | TCGTTGCATGTGGAACTAGCT |
| | | FTN_0485_Tn7_Det.REV | TGGTGAAACACTAGCTGAAGGT |
| pDMK3 *mtvS1-mCherry2* | | FTN_1238-tag_1.FOR | TCTATCGCCTTCTTGACGAGTTCTTCTGAATTGTCGACCCTCGAGTAGGTAACATCGCTAGTACAC |
| | | FTN_1238-mCherry2_1.REV | CTGTACAAGTTATAGTTATCAGGAGTAGGCTTATT |
| | | FTN_1238-mCherry2_2.FOR | ACTCCTGATAACTATAACTTGTACAGCTCG |
| | | FTN_1238-mCherry2_2.REV | AAAAAATTGCGGCCGCAGGA |
| | | FTN_1238-mCherry2_3.FOR | CGGCCGCAATTTTTTCACTATCAATAAGTAAAAC |
| | | FTN_1238-mCherry2_3.REV | AGAGCTCAGGTTACGCGGCCGCGATTACATAGCCATTATAAATCAC |
| | | FTN_1238-tag_Det.FOR | GCCGAAGCGATCTTGGACTA |
| | | FTN_1238_ns_Det.REV | TTTGCTGCTCATCAAAGGCT |

| | | | |
|---|---|---|---|
| pDMK3 *mtvS1-His*$_6$ | Added: GGTCACCACCACCACCACCAC | FTN_1238-tag_1.FOR | TCTATCGCCTTCTTGACGAGTTCTTCTGAATTGTCGACCCTCGAGTAGGTAACATCGCTAGTACAC |
| | | FTN_1238-His_1.REV | GGTCACCACCACCACCACCACTAGGTATTAACATGAATACTAAATATGATGTG |
| | | FTN_1238-His_2.FOR | CTAGTGGTGGTGGTGGTGGTGACCAATTTTTTCACTATCAATAAGTAAAAC |
| | | FTN_1238-His_2.REV | ATAACAATTTGTGGAATTCCCGGGAGAGCTCAGGTTACGCGGCCGCATGGGCGATGTATATTTCTTTTATGATGTGAAT |
| | | FTN_1238-tag_Det.FOR | GCCGAAGCGATCTTGGACTA |
| | | FTN_1238_ns_Det.REV | TTTGCTGCTCATCAAAGGCT |
| | | His_Det.REV | CACCACCACCACCACCAC |
| pDMK3 *mtvS2 nonsense* | MKKPNFDINK***PND | FTN_1519_ns_1.FOR | CTTCTGAATTGTCGACCCTCGAGCAGACATTTTTCGC |
| | | FTN_1519_ns_1.REV | ATTAATAAATAATAATAACCTAATGATGATTTAGTAGGC |
| | | FTN_1519_ns_2.FOR | ATTAGGTTATTATTATTTATTAATATCAAAATTAGGTTTCTTCA |
| | | FTN_1519_ns_2.REV | AGAGCTCAGGTTACGCGGCCGCGCACCACAAATTCG |
| | | dFTN_1519_Det.FOR | CCCTAGCAATAACTCGTTGGGT |
| | | dFTN_1519_Det.REV | AGAGAATGCAAGCTTAGCAAAGT |
| pDMK3 *mtvS2-His*$_6$ | Added: GGTCACCACCACCACCACCAC | FTN_1519_ns_1.FOR | CTTCTGAATTGTCGACCCTCGAGCAGACATTTTTCGC |
| | | FTN_1519-His_1.REV | AACGGTCACCACCACCACCACCACTAATAAATGCAAAGTCAGTCG |
| | | FTN_1519-His_2.For | ATTAGTGGTGGTGGTGGTGGTGACCGTTTTCTCTAATTTTAGCTAACTC |
| | | FTN_1519_ns_2.REV | ATAACAATTTGTGGAATTCCCGGGAGAGCTCAGGTTACGCGGCCGCATGGGCGATGTATATTTCTTTTATGATGTGAAT |
| pDMK3 Δ*FTN_0324* | MITVTNDKWTIETELTLKTVLSLAKVHGAKTILEEYIN* | dFTN_0324_1.FOR | CTTCTGAATTGTCGACCCTCGAGAATATTAGTACTGTTAACAAAGC |
| | | dFTN_0324_1.REV | CTAAAAACTGTTCTTTCTTTAGCAAAGGTG |
| | | dFTN_0324_2.FOR | TGCTAAAGAAAGAACAGTTTTTAGCGTTAATT |
| | | dFTN_0324_2.REV | AGAGCTCAGGTTACGCGGCCGCCATAGATTAACTTTAAGAGCTTT |
| | | dFTN_0324_Det.FOR | AGTGTTGGTACGACAGGTGC |
| | | dFTN_0324_Det.REV | ACACTTTTGTTGCAGGCAAAGA |
| pDMK3 Δ*FTN_0392* | MRITLKQLQVFIEKDINGCDIRLENFSI* | dFTN_0392_1.FOR | TCTATCGCCTTCTTGACGAGTTCTTCTGAATTGTCGACCCTCGAGAAGTGTTCTTACGTACA |
| | | dFTN_0392_1.REV | CTCTAAAACAGCTTCAAGTGTTTATAGAGAAAGATATAAATGGCTG |
| | | dFTN_0392_2.FOR | TTTATATCTTTCTCTATAAACACTTGAAGCTGTTT |
| | | dFTN_0392_2.REV | ATAACAATTTGTGGAATTCCCGGGAGAGCTCAGGTTACGCGGCCGCGTGAGACGAACCAAAAAC |
| | | dFTN_0392_Det.FOR | TGCAAATTCACTATAGCGCCA |
| | | dFTN_0392_Det.REV | TTGCAACATCCTCAACCCCA |
| pDMK3 Δ*FTN_0710* | MKKNNSQFSTQKAYELGVNNYD* | dFTN_0710_1.FOR | TCAGTACTCGAGTTATACATTATAAAGAAGTTTTTATCCC |
| | | dFTN_0710_1.REV | GTAAGCTTTTTGAGTTGAAAATTGAGAATTGTT |
| | | dFTN_0710_2.FOR | AATTTTCAACTCAAAAAGCTTACGAGCT |
| | | dFTN_0710_2.REV | TCAGTAGCGGCCGCGCTTGTAATGAAAATTCATGC |
| | | dFTN_0710_Det.FOR | AAGAAGCTATGCTCTCCTTTTGA |
| | | dFTN_0710_Det.REV | AAGCTATCGAAGGTGGCAAA |

| | | | |
|---|---|---|---|
| pDMK3 Δ*FTN_1282* | MGHLERLKIFIKAAHLSAYIEKNLPKIEL* | dFTN_1282_1.FOR | CTTCTGAATTGTCGACCCTCGAGGTTTTAGCTGTGTTTAAACC |
| | | dFTN_1282_1.REV | GCGCTTAGATGTGCTGCTTTTATAAATATC |
| | | dFTN_1282_2.FOR | CAGCACATCTAAGCGCTTATATAGAGAAA |
| | | dFTN_1282_2.REV | AGAGCTCAGGTTACGCGGCCGCGTTGTGATGGTACAGG |
| | | dFTN_1282_Det.FOR | ATCCTGCAGCAACAGAACCC |
| | | dFTN_1282_Det.REV | AGAGCAGCTCGAGGTAGTGA |
| pDMK3 Δ*FTN_1487* | MLFEKQQYQEDCVNN ELSNILQECQK* | dFTN_1487_1.FOR | TCAGTACTCGAGTCCTTGGAATATAGTTAAGTAATTAATT |
| | | dFTN_1487_1.REV | AGATTGTGTAAATAATGAATTATCTAATATTTTACAGGAGTG |
| | | dFTN_1487_2.FOR | AAATATTAGATAATTCATTATTTACACAATCTTCTTGATAT |
| | | dFTN_1487_2.REV | TCAGTAGCGGCCGCCGTTAGATATAGTTTTAGCAG |
| | | dFTN_1487_Det.FOR | CTGATGCAGCTTGGGAAAAT |
| | | dFTN_1487_Det.REV | TGAAACAGTATGGGCTAGTCAGA |
| pFNMB2 *mcs* | Multiple cloning site | pFNMB2_mcs.FOR | CTGCAGATTTAAGAAGGAGATACGCGTACTAGTGGGGCC |
| | | pFNMB2_mcs.REV | CGGGTTATTTGTAGAGCTCAGGTTACGCGGC |
| | | pFNMB_Det.FOR | TCATAGAAGCTTGCATGCCTG |
| | | pFNMB_Det.REV | AGACCCCACACTACCATCG |
| pFNMB *prom-cas12 mCherry2* | | pFNMB_prom-FTN_1397_1.FOR | TTAATCAGATAAAATATTTCTAGAACTAGTCTAGCTAGTTTAAAAGGACAA |
| | | FTN_1397_Prom_mCherry2_1.REV | TGCTCACCATGATAAAGACTCCTTATAAAATT |
| | | FTN_1397_Prom_mCherry2_2.FOR | GTCTTTATCATGGTGAGCAAGGGC |
| | | pFNMB_mCherry2.REV | TGTTTTATCAGACCGCCCCGGGTTACTTGTACAGCTCGTCCA |
| | | pFNMB_Det.FOR | TCATAGAAGCTTGCATGCCTG |
| | | pFNMB_Det.REV | AGACCCCACACTACCATCG |
| pFNMB *prom-upstream mtvS2* | | pFNMB_upstream-Prom-FTN_1519.FOR | TTAATCAGATAAAATATTTCTAGAACTAGTTATATGGAAACAATGGCTTC |
| | | pFNMB_FTN_1519.REV | TGTTTTATCAGACCGCCCCGGGTTAGTTTTCTCTAATTTTAGCTAACT |
| | | pFNMB_Det.FOR | TCATAGAAGCTTGCATGCCTG |
| | | pFNMB_Det.REV | AGACCCCACACTACCATCG |
| pFNMB *prom mtvS1* | | pFNMB_Prom-FTN_1238.FOR | TTAATCAGATAAAATATTTCTAGAACTAGTAAACTCTTGATTACATAGCCATTA |
| | | pFNMB_FTN_1238.REV | TGTTTTATCAGACCGCCCCGGGCTAAATTTTTTCACTATCAATAAGT |
| | | pFNMB_Det.FOR | TCATAGAAGCTTGCATGCCTG |
| | | pFNMB_Det.REV | AGACCCCACACTACCATCG |
| pFNMB *prom ygfB* | | pFNMB_Prom-b2909.FOR | TTAATCAGATAAAATATTTCTAGAACTAGTTGATGAGAAGAGACAAGC |
| | | pFNMB_b2909.REV | TGTTTTATCAGACCGCCCCGGGTTAGTGTAGAGTCGGTTTTT |
| | | pFNMB_Det.FOR | TCATAGAAGCTTGCATGCCTG |
| | | pFNMB_Det.REV | AGACCCCACACTACCATCG |
| pFNMB2 *msfGFP sp3_typeV_seedG3T_targ sp5_typeV_seedA3C_targ* | Protopacer 3 with seed mutation G3C and TTTG PAM: TTTGGTTTAGAGCCTTTTGTATTAGTAGCCG | pFNMB_TypeV_sp3_seedC3T.FOR | CGGCTACTAATACAAAAGGCTCTAAACCAAATTCTAGAAATATTTTATCTGATTAA |
| | | pFNMB_TypeV_sp3_seedC3T.REV | GTTTAGAGCCTTTTGTATTAGTAGCCGCTAGTTTAAGACCCACTTTCACATT |
| | Protopacer 5 with seed mutation A3C and TTTG PAM: TTTGAGCTTAAAAGGTAATTCTATCTTGTTGAG | pFNMB_TypeV_sp5_seedA3C.FOR | CTCAACAAGATAGAATTACCTTTTAATCTCAAAGAGCTCATCCATGCCGTGCG |
| | | pFNMB_TypeV_sp5_seedA3C.REV | AGCTTAAAAGGTAATTCTATCTTGTTGAGCCCGGGGCGGTCTGA |

| | | | |
|---|---|---|---|
| pFNMB *promoterless aptamer* | See Supplementary Table 11 | pFNMB_Promoterless-aptamer.FOR | TTAATCAGATAAAATATTTCTAGAACTAGTATGTCAATTTATCAAGAATTTG T |
| | | pFNMB_aptamer.REV | TGTTTTATCAGACCGCCCCGGGGAGAGCGTTCACCG |
| | | pFNMB_Det.FOR | TCATAGAAGCTTGCATGCCTG |
| | | pFNMB_Det.REV | AGACCCCACACTACCATCG |
| pFNMB *cas9 promoter aptamer* | See Supplementary Table 11 | pFNMB_Prom-cas9-aptamer.FOR | TTAATCAGATAAAATATTTCTAGAACTAGTTTAAATTTATAACCTATAAAGA AATTTCAG |
| | | Prom-cas9-aptamer_1.REV | GATAAATTGACATAAAATGACCTCTTTT |
| | | Prom-cas9-aptamer_2.FOR | AGAGGTCATTTTATGTCAATTTATCAAG |
| | | pFNMB_aptamer.REV | TGTTTTATCAGACCGCCCCGGGGAGAGCGTTCACCG |
| | | pFNMB_Det.FOR | TCATAGAAGCTTGCATGCCTG |
| | | pFNMB_Det.REV | AGACCCCACACTACCATCG |
| pFNMB *cas12 promoter aptamer* | See Supplementary Table 11 | pFNMB_Prom-cas12-aptamer.FOR | TTAATCAGATAAAATATTTCTAGAACTAGTAGATGAAACGCTAAAAATATAT G |
| | | Prom-cas12-aptamer_1.REV | ATAAATTGACATGATAAAGACTCCTTA |
| | | Prom-cas12-aptamer_2.FOR | GGAGTCTTTATCATGTCAATTTATCAAG |
| | | pFNMB_aptamer.REV | TGTTTTATCAGACCGCCCCGGGGAGAGCGTTCACCG |
| | | pFNMB_Det.FOR | TCATAGAAGCTTGCATGCCTG |
| | | pFNMB_Det.REV | AGACCCCACACTACCATCG |
| pET21 *mtvS1-His*$_6$ | | pET21.FOR | GGTCACCACCACCACCACCACTG |
| | | pET21.REV | ATGTATATCTCCTTCTTAAAGTTAAACAAAATTATTTCTAGAGGG |
| | | pET21_FTN_1238.FOR | TTAACTTTAAGAAGGAGATATACATATGGGCGATGTATATTTC |
| | | pET21_FTN_1238-His.REV | GTGGTGGTGGTGACCAATTTTTTCACTATCAATAAGTAAAAC |
| pET21 *mtvS2-His*$_6$ | | pET21_stop.FOR | TAATAGGTAAGCTGAATCCGGCTGCTAACAAAGCC |
| | | pET21.REV | ATGTATATCTCCTTCTTAAAGTTAAACAAAATTATTTCTAGAGGG |
| | | pET21_FTN_1519-His.FOR | TTAACTTTAAGAAGGAGATATACATATGAAGAAACCTAATTTTGATATTAA |
| | | pET21_FTN_1519-His_stop.REV | TCAGCTTACCTATTATCAGTGGTGGTGGTGGTGGTGACCGTTTTCTCTAA TTTTAGCTAACTC |
| | Ampicillin resistance cassette with 50 bp homology arms to *ygfB* | ygfB::ampR.FOR | GCCATGAGCTGTCATCGTTACCGCCACATATCATCCCGCTGATTAAACCA TTACCAATGCTTAATCAGTGAG |
| | | ygfB::ampR.REV | AACCAGTATCTGAACCAACAAGGGACGGGTCTGACCCCAGCTGAGATGC ACGCGGAACCCCTATTT |
| | | ygfB::ampR_Det.FOR | GCATCAGCCCGATCGAAAAC |
| | | ygfB::ampR_Det.REV | AAGCCCTTTTCTGGTCCACC |

**Supplementary Table 4: Sequencing and EDGE-Pro statistics for comparison of gene expression between exponentially growing and stationary phase cells of parental strain and *mtvS1 nonsense* mutant. RPKM reads were calculated as “Mapped reads”- “rRNA reads” and were the basis to calculate RPKM values.**

| | **Parental strain Exponential growth Replicate 1** | **Parental strain Exponential growth Replicate 2** | **Parental strain Exponential growth Replicate 3** | **Parental strain Stationary phase Replicate 1** | **Parental strain Stationary phase Replicate 2** | **Parental strain Stationary phase Replicate 3** |
|---|---|---|---|---|---|---|
| Total reads | 2986954 | 2712522 | 2765874 | 6025619 | 4080967 | 4575747 |
| Mapped reads | 2913730 | 2612638 | 2669732 | 5818536 | 3806447 | 4336287 |
| RPKM reads | 2899145 | 2463275 | 2644707 | 5766824 | 2503016 | 3178846 |
| rRNA reads | 14585 | 149363 | 25025 | 51712 | 1303431 | 1157441 |
| Unique reads | 2784875 | 2363729 | 2528180 | 5747217 | 2500232 | 3176617 |
| Multiple reads | 128855 | 248909 | 141552 | 71319 | 1306215 | 1159670 |

| | ***mtvS1 nonsense* Exponential growth Replicate 1** | ***mtvS1 nonsense* Exponential growth Replicate 2** | ***mtvS1 nonsense* Exponential growth Replicate 3** | ***mtvS1 nonsense* Stationary phase Replicate 1** | ***mtvS1 nonsense* Stationary phase Replicate 2** | ***mtvS1 nonsense* Stationary phase Replicate 3** |
|---|---|---|---|---|---|---|
| Total reads | 2061676 | 2567719 | 2530234 | 5902835 | 5486068 | 6047529 |
| Mapped reads | 1967414 | 2482642 | 2466641 | 5669047 | 5222292 | 5744518 |
| RPKM reads | 1965250 | 2477719 | 2447727 | 5446356 | 3641674 | 3529729 |
| rRNA reads | 2164 | 4923 | 18914 | 222691 | 1580618 | 2214789 |
| Unique reads | 1892514 | 2383803 | 2334436 | 5438902 | 3636613 | 3525217 |
| Multiple reads | 74900 | 98839 | 132205 | 230145 | 1585679 | 2219301 |

**Supplementary Table 5: Sequencing and EDGE-Pro statistics for comparison of gene expression between exponentially growing and stationary phase cells of parental strain and *mtvS2 nonsense* mutant. RPKM reads were calculated as "Mapped reads"- "rRNA reads" and were the basis to calculate RPKM values.**

| | Parental strain Exponential growth Replicate 1 | Parental strain Exponential growth Replicate 2 | Parental strain Exponential growth Replicate 3 | Parental strain Stationary phase Replicate 1 | Parental strain Stationary phase Replicate 2 | Parental strain Stationary phase Replicate 3 |
|---|---|---|---|---|---|---|
| Total reads | 8343386 | 3839674 | 3456171 | 3661319 | 3203981 | 2860169 |
| Mapped reads | 7896369 | 3615019 | 3233160 | 3347984 | 2996038 | 2697736 |
| RPKM reads | 7881916 | 3604581 | 3226269 | 2284452 | 2416302 | 2611472 |
| rRNA reads | 14453 | 10438 | 6891 | 1063532 | 579736 | 86264 |
| Unique reads | 7654326 | 3511513 | 3125523 | 2263161 | 2404676 | 2595038 |
| Multiple reads | 242043 | 103506 | 107637 | 1084823 | 591362 | 102698 |

| | *mtvS2 nonsense* Exponential growth Replicate 1 | *mtvS2 nonsense* Exponential growth Replicate 2 | *mtvS2 nonsense* Exponential growth Replicate 3 | *mtvS2 nonsense* Stationary phase Replicate 1 | *mtvS2 nonsense* Stationary phase Replicate 2 | *mtvS2 nonsense* Stationary phase Replicate 3 |
|---|---|---|---|---|---|---|
| Total reads | 3808430 | 4384771 | 3230924 | 3195029 | 3369067 | 2376528 |
| Mapped reads | 3639036 | 4199206 | 3072765 | 3028047 | 3201878 | 2282368 |
| RPKM reads | 3630774 | 4124349 | 3067361 | 1969882 | 2733793 | 2107863 |
| rRNA reads | 8262 | 74857 | 5404 | 1058165 | 468085 | 174505 |
| Unique reads | 3565246 | 4056135 | 3013881 | 1961332 | 2719408 | 2096063 |
| Multiple reads | 73790 | 143071 | 58884 | 1066715 | 482470 | 186305 |

**Supplementary Table 6: Sequencing and EDGE-Pro statistics for comparison of gene expression between exponentially growing and stationary phase cells of parental *E. coli* strain and *ygfB::ampR* mutant. RPKM reads were calculated as "Mapped reads"- "rRNA reads" and were the basis to calculate RPKM values.**

| | **Parental strain Exponential growth Replicate 1** | **Parental strain Exponential growth Replicate 2** | **Parental strain Exponential growth Replicate 3** | **Parental strain Stationary phase Replicate 1** | **Parental strain Stationary phase Replicate 2** | **Parental strain Stationary phase Replicate 3** |
|---|---|---|---|---|---|---|
| Total reads | 36054722 | 34304644 | 36224450 | 32786560 | 33084758 | 33454216 |
| Mapped reads | 35315609 | 33583473 | 35426675 | 30923012 | 30522365 | 30775577 |
| RPKM reads | 34848479 | 33557224 | 34954563 | 30835863 | 30367552 | 30524275 |
| rRNA reads | 467130 | 26249 | 472112 | 87149 | 154813 | 251302 |
| Unique reads | 34320171 | 33035446 | 34390746 | 30814664 | 30329727 | 30489307 |
| Multiple reads | 995438 | 548027 | 1035929 | 108348 | 192638 | 286270 |

| | ***ygfB::ampR*** **Exponential growth Replicate 1** | ***ygfB::ampR*** **Exponential growth Replicate 2** | ***ygfB::ampR*** **Exponential growth Replicate 3** | ***ygfB::ampR*** **Stationary phase Replicate 1** | ***ygfB::ampR*** **Stationary phase Replicate 2** | ***ygfB::ampR*** **Stationary phase Replicate 3** |
|---|---|---|---|---|---|---|
| Total reads | 35581174 | 35065810 | 34079434 | 35337534 | 36245370 | 36009826 |
| Mapped reads | 34997515 | 32587796 | 33452416 | 33606379 | 34180348 | 33920664 |
| RPKM reads | 34981316 | 32571859 | 33440952 | 31792241 | 34120083 | 33853745 |
| rRNA reads | 16199 | 15937 | 11464 | 1814138 | 60265 | 66919 |
| Unique reads | 34390494 | 32034671 | 32857912 | 31899435 | 34092439 | 33830137 |
| Multiple reads | 607021 | 553125 | 594504 | 1706944 | 87909 | 90527 |

### Supplementary Table 7: Primers used for RT-qPCR.

| Primers | Sequence 5'-3' [base pairs] |
|---|---|
| FTN_0757_qPCR. FOR | ATGCTAACACTTGTGCAGTTTG |
| FTN_0757_qPCR.REV | GTCTCTGAGCTTTGGCACTTA |
| FTN_1397_qPCR.FOR | AAGCTCTGCCTCACCATTTAG |
| FTN_1397_qPCR.REV | AAGGGCGACCAAATCTACATAC |
| FTN_1396_qPCR.FOR | GCCCATCTCTTGGAGACAAA |
| FTN_1396_qPCR.REV | GGGCTTACTTGTGGAGAGAAA |
| FTN_1395_qPCR.FOR | TATTCTAGGTTTGCGCCCTTG |
| FTN_1395_qPCR.REV | CAAGCCTAGCGGAGCTAATG |
| FTN_1394_qPCR.FOR | TCACATCTTTATCTGGAGCATT |
| FTN_1394_qPCR.REV | TGGTGTACGTTTACAATATTCGG |
| FTN_1238_qPCR.FOR | CACCCTCTTCAATCGGCTTAG |
| FTN_1238_qPCR.REV | GGACTGCTTTGTGCTTTGTTT |
| FTN_1519_qPCR.FOR | ATTCTTCCTGCAAAGCACATATC |
| FTN_1519_qPCR.REV | AGAATGGATCTTGAGTGTGAAGT |
| FTN_1594_qPCR.FOR | GAGTCTATCTCTGCGACGAAAC |
| FTN_1594_qPCR.REV | GTAACAAGAGCTATGAAAGCCTTAAC |
| b2756_qPCR.REV | CAGGTTGGTGTTCCACTCTATT |
| b2756_qPCR.REV | CAGGTTGGTGTTCCACTCTATT |
| b2758_qPCR.FOR | CCTGAAGTTGAGCGAGGTTAAT |
| b2758_qPCR.REV | GGTTCACCGCTGTAGATGATT |
| b2760_qPCR.FOR | ACGTTGTTACAGGTGTTCCTC |
| b2760_qPCR.REV | GCACTGCGATTGCGTTATTC |
| b3699_qPCR.FOR | TGTTCCTGCTTGCCTTTCT |
| b3699_qPCR.REV | GCTGCTGTTGACCTTCTTCTA |

### Supplementary Table 8: Template switch oligo and primers used for 5' RACE.

| Template switch oligo/ Primers | Sequence 5'-3' [base pairs] |
|---|---|
| Template switch oligo | GCTAATCATTGCAAGCAGTGGTATCAACGCAGAGTACATRGRGRG |
| FTN_1397-RT.REV | CTTTAGCTCTTTTCTCATCATCT |
| TSO.FOR | CATTGCAAGCAGTGGTATCAAC |
| FTN_1397-RT_PCR2.REV | ACCTCTTGCTTTTATGTTTTCAAGTGTTTTACCCT |

### Supplementary Table 9: Primers used for promoter sequence pull-down.

| Primers | Modification | Sequence 5'-3' [base pairs] |
|---|---|---|
| Promoter-FTN_1397.FOR | | CTAGCTAGTTTAAAAGGACAA |
| Promoter-FTN_1397_biotin.REV | 5' Biotin-TEG | TAACTCAAATCTTAGAGTTTTACTT |
| FTN_1397_pull-down.FOR | | GCTGTACCAATAACACTATAAT |
| FTN_1397_biotin.REV | 5' Biotin-TEG | AGAAAAATCTATTAAAGAAACACTATC |

### Supplementary Table 10: Primers used for EMSA.

| Sequences and Primers | Sequence 5'-3' [base pairs] |
|---|---|
| Promoter-FTN_1397.FOR | CTAGCTAGTTTAAAAGGACAA |
| Prom-FTN_1397_EMSA.REV | TAACTCAAATCTTAGAGTTTTACTT |
| FTN_1397_pull-down.FOR | GCTGTACCAATAACACTATAAT |
| FTN_1397_EMSA.REV | TGATGTAGTTACAACGATGC |

### Supplementary Table 11: Aptamer sequences used for *in vitro* transcription assays.

| Aptamers and Primers | Sequence 5’-3’ [base pairs] |
|---|---|
| Promoterless aptamer | ATGTCAATTTATCAAGAATTTGTTAATAAATAGACGCGACCGAAATGGTGAAGGACGGGTCCAGTGCTTCGGCACTGTTGAGTAGAGTGTGAGCTCCGTA<br>ACTGGTCGCGTCTAGTTTAAGTAAAACTCTAAGATTTGAGTTAACCAGGCATCAAATAAAACGAAAGGCTCAGTCGAAAGACTGGGCCTTTCGTTTTATCT<br>GTTGTTTGTCGGTGAACGCTCTCC |
| Promoterless-aptamer.FOR | ATGTCAATTTATCAAGAATTTGTT |
| Aptamer.REV | GGAGAGCGTTCACCGACAAA |
| *cas9* promoter aptamer | TTAAATTTATAACCTATAAAGAAATTTCAGTGAGTTATTTACTAAAATTCATAGAATCATTCATACTTTCTAGAAATCAAATAATATTTACATAAATCGTAGCTA<br>ATACTGACAATATTTCTTTCAAAATGATTTATAAAATTTGTAATTTTAGATTAGTTTTACTATTAGGAAAATTGATAGTTATATAGAGCTTTATGAAGTAGAATT<br>AATGTATAAGTTTAACTACTTTTCCATTTTTTAATTTGATTTTGATTTTAAATGCTGGTTTAATGCTTTATTAAACCAATGATAAACGTATGTGAAGCTAGTTC<br>CTAAAAGCTTTTATCCAAGTTTATAGATTTATGATTAAAAACTTATATCAAAATATTATTCTATACCGAGTAAAAGAGGTCATTTTATGTCAATTTATCAAGAA<br>TTTGTTAATAAATAGACGCGACCGAAATGGTGAAGGACGGGTCCAGTGCTTCGGCACTGTTGAGTAGAGTGTGAGCTCCGTAACTGGTCGCGTCTAGTT<br>TAAGTAAAACTCTAAGATTTGAGTTAACCAGGCATCAAATAAAACGAAAGGCTCAGTCGAAAGACTGGGCCTTTCGTTTTATCTGTTGTTTGTCGGTGAAC<br>GCTCTCC |
| FTN_0757-aptamer_1.FOR | TTAAATTTATAACCTATAAAGAAATTTCAG |
| FTN_0757-aptamer_1.REV | GATAAATTGACATAAAATGACCTCTTTT |
| FTN_0757-aptamer_2.FOR | AGAGGTCATTTTATGTCAATTTATCAAG |
| Aptamer.REV | GGAGAGCGTTCACCGACAAA |
| *cas12* promoter aptamer | CTAGTAGATGAAACGCTAAAAATATATGCCCTTGCACCAATGTCTGTGGCACCAAAATATCAACATCAAGGAATTGGTTCTAAGCTTATAGAAGCAATGAT<br>TAAGGAAGCCAAAAAAAATAATATTGATGCAATATTTGTCTTAGGTCATCCAAGTTATTATCCAAAATTTGGTTTTAAACCAGCCACAGAATATCAGATAAAA<br>TGTGAATATGATGTCCCAGCGGATGTTTTTATGGTACTAGATTTGTCAGCTAAACTAGCTAGTTTAAAAGGACAAACTGTCTACTATGCCGATGAGTTTGG<br>CAAAATTTTTTAGATCTACAAAATTATAAACTAAATAAAGATTCTTATAATAACTTTATATATAATCGAAATGTAGAGAATTTTATAAGGAGTCTTTATCATGTC<br>AATTTATCAAGAATTTGTTAATAAATAGACGCGACCGAAATGGTGAAGGACGGGTCCAGTGCTTCGGCACTGTTGAGTAGAGTGTGAGCTCCGTAACTGG<br>TCGCGTCTAGTTTAAGTAAAACTCTAAGATTTGAGTTAACCAGGCATCAAATAAAACGAAAGGCTCAGTCGAAAGACTGGGCCTTTCGTTTTATCTGTTGT<br>TTGTCGGTGAACGCTCTCC |
| FTN_1397-aptamer_1.FOR | GCGCGCGCGCAGATGAAACGCTAAAAATATATG |
| FTN_1397-aptamer_1.REV | ATAAATTGACATGATAAAGACTCCTTA |
| FTN_1397-aptamer_2.REV | GGAGTCTTTATCATGTCAATTTATCAAG |
| Aptamer.REV | GGAGAGCGTTCACCGACAAA |
